# Exploring Dynamic Modulation of Binding, Allostery and Immune Resistance in the SARS-CoV-2 Spike Complexes with Classes of Antibodies Targeting Cryptic Binding Sites: Antibody-Specific Augmentations of Conserved Allosteric Architecture Can Influence Evolution of Viral Escape

**DOI:** 10.1101/2025.06.28.662162

**Authors:** Mohammed Alshahrani, Vedant Parikh, Brandon Foley, Gennady Verkhivker

**Author notes:** Correspondence; (G.V). (M.A); (V.P.); (B.F.); (G.V).

## Abstract

The ongoing evolution of SARS-CoV-2 variants has underscored the need to understand not only the structural basis of antibody recognition but also the dynamic and allosteric mechanisms that could underlie complexity of broad and escape-resistant neutralization. In this study, we employed a multi-scale approach integrating structural analysis, hierarchical molecular simulations, mutational scanning and network-based allosteric modeling to dissect how Class 4 antibodies (represented by S2X35, 25F9, and SA55) and Class 5 antibodies (represented by S2H97, WRAIR-2063 and WRAIR-2134) can modulate conformational behavior, binding energetics, allosteric interactions and immune escape patterns of the SARS-CoV-2 spike protein. Using hierarchical simulations of the antibody complexes with the spike protein and ensemble-based mutational scanning of binding interactions we showed that these antibodies through targeting conserved cryptic sites can exert allosteric effects that influence global conformational dynamics in the RBD functional regions. The ensemble-based mutational scanning of binding interactions revealed an excellent agreement with experimentally derived deep mutational scanning (DMS) data accurately recapitulating the known binding hotspots and escape mutations across all studied antibodies. The predicted destabilization values in functional sites are consistent with experimentally observed reductions in antibody binding affinity and immune escape profiles demonstrating that computational models can robustly reproduce and forecast mutation-induced immune escape trends. Using dynamic network modeling we characterized the antibody-induced changes in residue interaction networks and long-range interactions. The results revealed that class 4 antibodies can exhibit distinct patterns of allosteric influence despite targeting overlapping regions, while class 5 antibodies elicit consistently dense and broadly distributed allosteric networks and long-range stabilization of the RBD conformations. Dynamic network analysis identifies a conserved allosteric network core that mediates long-range interactions and incudes antibody specific allosteric extensions that connect the binding interface hotspots with allosteric hubs. This study suggests that mechanisms of binding and immune escape for classes of antibodies targeting cryptic binding sites may be determined by confluence of multiple factors including high-affinity binding and long-range allosteric effects that modulate RBD adaptability and propagation of dynamic constraints which can influence mechanisms of viral evolution and escape.

## Introduction

Structural and biochemical studies of the SARS-CoV-2 Spike (S) glycoprotein have unveiled critical insights into the mechanisms driving viral transmission, immune evasion, and host-cell entry.^1–9^ The S glycoprotein is distinguished by its conformational flexibility and functional plasticity, particularly within the S1 subunit, which encompasses several key domains: the N-terminal domain (NTD), the receptor-binding domain (RBD), and two structurally conserved subdomains, SD1 and SD2. This intrinsic flexibility enables the S glycoprotein to dynamically adapt to different stages of the viral entry process, enhancing both its functionality and ability to evade immune detection.^1–9^ The RBD plays a pivotal role in binding to the angiotensin-converting enzyme 2 (ACE2) receptor. ^10–15^ Conformational changes in the NTD and RBD drive transitions between the closed and open states of the S protein, allowing the virus to efficiently engage with host receptors while leveraging structural variability to evade immune surveillance. Biophysical investigations revealed that mutations within the S protein, particularly in the S1 subunit, can induce structural alterations that affect its stability and conformational dynamics.^16–18^ The structural variability introduced by these mutations enhances the virus capacity to evade immune responses, complicating the host ability to mount an effective defense. The enormous wealth of cryo-electron microscopy (cryo-EM) and X-ray structures of SARS-CoV-2 S protein variants of concern (VOCs) in various functional states, along with their interactions with antibodies, highlighted how VOCs can induce structural changes in the dynamic equilibrium of the S protein and control balance between structural stability, immune evasion, and receptor binding that shapes the evolutionary trajectory of SARS-CoV-2 and its variants.^19–25^

The emergence and evolution of SARS-CoV-2 Omicron variants represented critical milestones in the evolutionary trajectory of the virus due to their enhanced growth advantages and transmissibility that optimize receptor binding while enhancing immune evasion capabilities.^26–28^

XBB variants, descendants of the BA.2 lineage, arose through recombination events introducing key mutations in the RBD enhancing ACE2 binding affinity and infectivity.^26–28^ XBB descendants EG.5 and EG.5.1 carry an additional F456L mutation, boosting immune escape.^29–32^ The “FLip” variants, featuring L455F and F456L mutations, exemplify convergent evolution, enabling XBB strains to outcompete others by balancing immune evasion with receptor binding efficiency.^33^ BA.2.86, another BA.2 descendant, demonstrated significant genetic divergence and heightened immune evasion against RBD-targeted antibodies, surpassing XBB.1.5 and EG.5.1 variants.^34–38^ Its descendant, JN.1, acquired an L455S mutation, further evading immune responses and reducing ACE2 binding affinity while resisting class-1 and class-3 antibodies.^39^ Subsequent variants including “SLip” (L455S + F456L) and “FLiRT” (additional R346T) enhanced immune escape and maintained ACE2 binding.^40–43^ JN.1 subvariants KP.2 and KP.3 independently gained mutations such as R346T, F456L, Q493E, and V1104L, improving transmissibility and immune evasion.^44^ Other JN.1 subvariants, including LB.1 and KP.2.3, have also emerged, sharing mutations S:R346T and S:F456L while acquiring unique changes such as S:S31- and S:Q183H (LB.1) or S:H146Q (KP.2.3). These convergent mutations contributed to increased immune evasion.^45^ KP.3 (“FLuQE”), carrying L455S, also emerged as highly immune-evasive.^46^ The convergence of mutations L455F, F456L, and R346T highlights the intense selective pressure driving viral evolution, as the virus continually seeks to optimize its fitness in the face of immune challenges Recent cryo-EM studies of JN.1, KP.2, and KP.3 RBD complexes revealed that F456L mutation enhances the binding potential of Q493E, leading to stronger interactions of KP.3 with the ACE2 receptor.^47^ This synergy provides an additional evolutionary advantage, enabling the virus to incorporate additional immune-evasive mutations while maintaining high infectivity. XEC, a recombinant KP.3 variant with NTD mutations F59S and T22N, showed higher infectivity and resistance to immune responses.^48–52^ KP.3.1.1, derived via S31 deletion, exhibited epistatic synergy between F456L and Q493E, restoring ACE2 affinity despite reduced binding from Q493E alone.^53,54^ LP.8 and its descendant LP.8.1 (“DeFLiRT”) feature mutations S31del, F186L, Q493E, and R190S, contributing to immune evasion and growth advantages.^55^ LF.7.2.1 (with A475V) and LP.8.1 (R190S) demonstrated significant immune evasion and enhanced ACE2 engagement, respectively.^55^ BA.3.2, with over 50 mutations, showed antibody resistance but reduced ACE2 binding.^56^ These findings underscore the rapid evolutionary dynamics of SARS-CoV-2 and its remarkable capacity to adapt through mutations that balance immune evasion with functional constraints, optimizing mutational landscapes for fitness and transmissibility.

The fundamental aspect of the immune response to SARS-CoV-2 is the production of antibodies that target various regions of the S protein, which plays a central role in viral entry into host cells. Most of the SARS-CoV-2 S antibodies target RBD and can be divided into different classes on the basis of their targeted epitopes. Although several different classification systems have been proposed, the most commonly referenced is that proposed by Barnes and colleagues who grouped RBD-targeting antibodies into four classes on the basis of their binding epitope and mode of binding to the S protein.^57^ The neutralizing antibodies target four key regions within the SARS-CoV-2 S protein, including NTD and RBD in the S1 subunit, and the stem helix region and the fusion peptide region in the S2 subunit.^58^ Class 1 and 2 RBD-targeting antibodies have the binding epitope that overlaps with the receptor-binding motif (RBM) in the RBD, and antibodies in this class mostly recognize only the ‘up’ RBD conformation. Class 3 RBD-targeting antibodies bind the outside the ACE2-binding region, and they can also bind to RBDs regardless of their ‘up’ and ‘down’ conformations.^58^ This class of antibodies includes REGN10987, COV2-2130, 2-7,1-57, A19-61.1, P2G3, S309 and LY-CoV1404, have demonstrated potent neutralizing activities against SARS-CoV-2 variants. S309 targets a highly conserved epitope near the N343 glycosylation site within the RBD but does not overlap with the ACE2-binding motif.^59,60^ Class 4 RBD-targeting antibodies target highly conserved region in the RBD that do not directly block ACE2–RBD binding. This epitope is conserved by up to 86% in SARS-CoV and SARS-CoV-2 and has also been described as a cryptic region, based on a cryptic epitope recognized by the CR3022 antibody.^61^

High-throughput yeast display screening and deep mutational scanning (DMS) have revolutionized our understanding of the escape mutation profiles associated with the RBD residues of the S protein. These advanced techniques have enabled a rigorous mapping of the functional epitopes targeted by human anti-RBD neutralizing antibodies leading to a comprehensive classification of these antibodies into distinct epitope groups (A–F).^62^ Notably, groups E and F correspond to class 3 and class 4 antibodies respectively in earlier classification.^57^ Through the identification of 241 broad sarbecovirus neutralizing antibodies and using high-throughput yeast-display mutational screening to determine the RBD escaping mutation profile, a total of 6 clusters of antibodies with diverse breadth and epitopes were identified.^63^ These clusters included group E1 (S309 site^59,60^), E3 (S2H97 site^64^), F1 (CR3022^61^, S304 site^65^), F2 (DH1047site^66^), F3 (ADG-2 site^67.68^). This pioneering approach was expanded to characterize the epitope distribution of antibodies elicited by post-vaccination BA.1 infection and using the mutational escape profiles for 1,640 RBD-binding antibodies, the antibodies were classified into 12 epitope groups.^69^ In this classification, groups A–C consist of antibodies targeting the ACE2-binding motif that are highly effective at blocking the interaction between the virus and its host receptor, making them crucial for neutralization. Group D antibodies such as REGN-10987, LY-CoV1404, and COV2-2130, bind to the epitope 440–449 on the RBD and are further divided into D1 and D2 subgroups. Groups E and F that are of consideration in our study and target regions outside the ACE2-binding motif were subdivided into E1–E3 and F1– F3 subgroups, respectively, covering the front and back of the RBD.^69^ Using DMS profiles of the monoclonal antibodies isolated from BA.2 and BA.5 convalescent individuals, Cao and colleagues embedded all antibodies using multidimensional scaling based on their DMS profiles, followed by *t*-distributed stochastic neighbor embedding (*t*-SNE) and *k*-nearest neighbors-based classification to determine the epitope groups of new antibodies. This resulted in a dataset containing the DMS profiles of 3,051 SARS-CoV-2 WT RBD-targeting antibodies in which BA.1, BA.2 and BA.5 breakthrough infections mainly elicit antibodies of groups E2.2, E3 and F1, which do not compete with ACE2 and demonstrate weak neutralizing activity, whereas WT-elicited antibodies are enriched in groups A, B and C, which compete with ACE2 and exhibit strong neutralization potency.^70^

The molecular mechanisms underlying the broadly neutralizing antibodies induced by XBB/JN.1 infections was conducted using high-throughput yeast-display-based DMS assays and the escape mutation profiles in screening of a total 2,688 antibodies, including 1,874 isolated from XBB/JN.1 infection cohorts, resulting in 22 clusters. ^71^ In this study, Cao and colleagues showed the possibility of accurately predicting SARS-CoV-2 RBD evolution by aggregating high-throughput antibody DMS results and constructing pseudoviruses that carry the predicted mutations as filters to screen for antibodies.^71^ This innovative approach enabled the identification of E1 group antibodies BD55-3546, BD55-3152, BD55-5585, BD55-5549 and BD55-5840 (SA58) antibodies a well as F3 antibodies BD55-4637, BD55-3372, BD55-5483, and BD55-5514 (SA55).^72,73^ In the recent pioneering investigation, Cao’s group leveraged DMS to predict viral evolution and to select for monoclonal antibodies neutralizing both existing and prospective variants.^74^ Through this screening process, a retrospective analysis of 1,103 SARS-CoV-2 wild type (WT)-elicited monoclonal antibodies identified BD55-1205, the only WT-elicited monoclonal antibody exhibiting exceptional neutralization breadth against the latest variants and prospective mutants.^74^ In another seminal study, pan-sarbecovirus binding assays, in vitro mapping of viral escape, structural analyses and DMS experiments provided a comprehensive characterization of a panel of antibodies targeting different epitopes, including conserved cryptic RBD regions.^75^

Several neutralizing antibodies with exceptional sarbecovirus breadth and a corresponding resistance to immune escape were discovered, including antibody S2H97 that binds with high affinity to a novel cryptic epitope (site 5) and S2H97 is respectively annotated as class 5 antibody.^75^ Subsequent studies identified several other class 5 antibodies such as WRAIR-2057, WRAIR-2063, and WRAIR-2134 that showed broad cross-reactivity across SARS-CoV-2 VOC’s^76,77^. Structural studies revealed that WRAIR-2063 binds to a cryptic epitope 5 within the RBD that is typically occluded by the NTD of an adjacent spike protomer in the closed conformation of the spike trimer and th.is epitope becomes accessible only when at least one RBD adopts the up conformation.^76^ A recent structural and functional study reported reporting the crystal structures of several broadly reactive monoclonal antibodies including class 5 WRAIR-2134 antibody.^77^ The class 5 antibodies WRAIR-2134, WRAIR-2057, WRAIR-2063 and S2H97 share a closely related binding footprint binding to a highly structurally conserved and functionally relevant neutralizing cryptic epitope that becomes exposed only when the RBD is in an open conformation, requiring extensive opening of the RBD for binding.^75–77^ These studies suggested that a potential neutralization mechanism of class 5 antibodies implies induction of rapid and premature refolding of the spike into the post-fusion state.

Computer simulations have significantly advanced our understanding of the dynamics and functions of the S protein and S complexes with ACE2 and antibodies at the atomic level. Molecular dynamics (MD) simulations and Markov state models (MSM) have systematically characterized the conformational landscapes of XBB.1 and XBB.1.5 Omicron variants and their complexes.^78^ Mutational scanning and binding analysis of the Omicron XBB spike variants with ACE2 and a panel of class 1 antibodies provided a quantitative rationale for the experimental evidence.^79,80^ We combined AlphaFold2-based atomistic predictions of structures and conformational ensembles of the S complexes with the ACE2 for the most dominant Omicron variants JN.1, KP.1, KP.2 and KP.3 to examine the mechanisms underlying the role of convergent evolution hotspots in balancing ACE2 binding and Ab evasion.^81^ Our recent studies suggested a mechanism in which the pattern of specific escape mutants for ultrapotent antibodies may be driven by a complex balance between the impact of mutations on structural stability, binding strength, and long-range communications.^82,83^ Moreover, convergent Omicron mutations can display epistatic couplings with the major stability and binding affinity hotspots which may allow for the observed broad antibody resistance.^82,83^ Computational studies examined mechanisms of broadly neutralizing antibodies of E1 and F3 groups of antibodies that leverage strong hydrophobic interactions with the binding epitope hotspots critical for the RBD stability and ACE2 binding, while escape mutations tend to emerge in sites associated with synergistically strong hydrophobic and electrostatic interactions.^84^ In the recent modeling study, a comparative analysis of the binding mechanisms and resistance profiles of S309, S304, CYFN1006, and VIR-7229 revealed distinct molecular strategies employed by these antibodies that can be broadly categorized into two paradigms: (1) conservation-driven binding, where antibodies exploit highly conserved residues critical for viral function, and (2) adaptability-driven binding.^85^ Computational and experimental studies provided a compelling evidence that at the molecular level the viral evolution is a complex process driven by a fine-tuned balance of immune evasion and receptor-binding affinity, influenced by both genetic mutations and the diversity of antibody responses.^86–88^ The SARS-CoV-2 spike protein interaction with antibodies can also involve a critical interplay between binding and allostery. Some antibodies, particularly targeting conserved cryptic binding sites on the RBD and of the rigid S2 domain can achieve neutralization not just by blocking ACE2 binding, but also by inducing allosteric changes that disrupt the S protein ability to undergo necessary conformational changes for viral entry. The recent structure-functional investigation revealed that the conformational dynamics and allosteric perturbations are linked to binding of novel human antibodies where antibody-induced dynamics can render weak, moderate and strong neutralizing antibodies.^89^ According to this study, some ‘weak’ antibodies can show only a mild effect on the dynamics and induce long-range destabilization, while the strong neutralizing antibodies can elicit large-scale conformational rigidity of the S protein.

In this study, we examine the interplay of dynamic, energetic and allosteric effects in determining molecular mechanisms of binding and immune escape for class 4 and class 5 antibodies targeting distinct cryptic binding sites on the RBD. We employed a combination of coarse-grained and atomistic MD simulations, mutational scanning of binding interactions and dynamic interrogation of allosteric residue interaction networks of conformational ensembles to dissect binding and allosteric mechanisms of immune resistance and viral evolution of the S protein. For this study, we used a representative panel of class 4 antibodies (group F3 S2X35 antibody,^90^ group F3 25F9 antibody^91^ and group F3 SA55 antibody^72,73^) and panel of class 5 antibodies S2H97^75^, WRAIR-2063^76^ and WRAIR-2134^77^. In this panel class 4 antibodies 25F9 SA55 antibody showed excellent neutralization against latest variants JN.1, KP.2. KP.3, KP.3.1.1, XEC and exceptional immune escape.^92^ Using dynamic ensembles of the antibody complexes and systematic mutational scanning of the RBD and antibody residues we characterize patterns of mutational sensitivity and compute mutational scanning heatmaps to identify binding hotspots and escape mutations. Through network-based modeling of conformational ensembles, we also focus on allosteric effects induced by antibodies in which binding to conserved regions distant from the ACE2 binding site can cause conformational changes in the RBD and induce antibody-specific dynamics of the RBD conformations that allosterically interferes with the receptor binding. We show that neutralization effects of these antibodies targeting conserved cryptic sites reflects a dynamic equilibrium shaped by multiple factors, including the energetic contributions of specific binding interactions, the architecture of allosteric network, the distribution of escape hotspots across the protein, and the selective pressures exerted by diverse antibody repertoires. The results provide a broad perspective on the interplay of binding and allostery in shaping up mechanisms of immune defense that are energetically nuanced and context-dependent. The antibody-dependent variability of immune responses adds another layer of complexity to understanding how SARS-CoV-2 continues to adapt under selective pressures imposed by both natural immunity and confluence of binding and allosteric effects induced by new classes of antibodies.

## Materials and Methods

### Coarse-Grained Molecular Simulations and Atomistic Reconstruction of Ensembles

Although all-atom MD simulations with the explicit inclusion of the glycosylation shield can in principle provide a rigorous assessment of conformational landscape of the SARS-CoV-2 S proteins, such direct simulations may become challenging due to the size of a complete SARS-CoV-2 S system embedded onto the membrane and this complexity can often obscure the main molecular determinants of the binding mechanisms and prevents efficient comparative analysis across large number of antibodies. Coarse-grained (CG) models are computationally effective approaches for rapid and efficient exploration of large systems over long timescales. The crystal and cryo-EM structures of the RBD-antibody are obtained from the Protein Data Bank.^93^ The following systems were used in this study : the structure of class 4 S2X35 with RBD (pdb id 7R6W), class 4 25F9 with RBD (pdb id 8GB5), class 4 SA55 with Omicron BA.1 RBD (pdb id 7Y0W), class 5 S2H97 with RBD (8S6M), class 5 WRAIR-2063 with RBD (pdb id 8EOO) and class 5 WRAIR-2134 with RBD (pdb id 8F2J).

We employed CABS-flex approach that efficiently combines a high-resolution coarse-grained model and efficient search protocol capable of accurately reproducing all-atom MD simulation trajectories and dynamic profiles of large biomolecules on a long time scale.^94–99^ In this high-resolution model, the amino acid residues are represented by Cα, Cβ, the center of mass of side chains and another pseudoatom placed in the center of the Cα-Cα pseudo-bond. In this model, the amino acid residues are represented by Cα, Cβ, the center of mass of side chains and the center of the Cα-Cα pseudo-bond. The CABS-flex approach implemented as a Python 2.7 object-oriented standalone package was used in this study to allow for robust conformational sampling proven to accurately recapitulate all-atom MD simulation trajectories of proteins on a long time scale. Conformational sampling in the CABS-flex approach is conducted with the aid of Monte Carlo replica-exchange dynamics and involves local moves of individual amino acids in the protein structure and global moves of small fragments. The default settings were used in which soft native-like restraints are imposed only on pairs of residues fulfilling the following conditions : the distance between their *C*_α_ atoms was smaller than 8 Å, and both residues belong to the same secondary structure elements. A total of 1000 independent CG-CABS simulations were performed for each of the systems studied. In each simulation, the total number of cycles was set to 10,000 and the number of cycles between trajectory frames was 100. MODELLER-based reconstruction of simulation trajectories to all-atom representation^100^ provided by the CABS-flex package was employed to produce atomistic models of the equilibrium ensembles for studied systems.

### All-Atom MD Simulations and Analysis of Equilibrium Ensembles

The missing regions for the studied structures of the RBD-antibody are reconstructed and optimized using template-based loop prediction approach ArchPRED.^101^ The side chain rotamers were refined and optimized by SCWRL4 tool.^102^ The protonation states for all the titratable residues of the antibody and RBD proteins were predicted at pH 7.0 using Propka 3.1 software and web server.^103,104^ The glycan chains were built using CHARMM-GUI Glycan Reader^105,106^ and Modeller^100^ at glycosylation sites N331 and N343 of RBD. NAMD 2.13-multicore-CUDA package^107^ with CHARMM36m force field^108^ employed to perform all-atom MD simulations for the RBD-antibody complexes. Each system was solvated with TIP3P water molecules and neutralizing 0.15 M NaCl in a periodic box that extended 10 Å beyond any protein atom in the system.^109^ All Na+ and Cl− ions were placed at least 8 Å away from any protein atoms and from each other. MD simulations are typically performed in an aqueous environment in which the number of ions remains fixed for the duration of the simulation, with a minimally neutralizing ion environment or salt pairs to match the macroscopic salt concentration.^110^

The heavy atoms in the complex were restrained using a force constant of 1000 kJ mol^−1^ nm^−1^ to perform 500 ps equilibration simulation. Long-range, non-bonded van der Waals interactions were computed using an atom-based cutoff of 12 Å, with the switching function beginning at 10 Å and reaching zero at 14 Å. The SHAKE method was used to constrain all the bonds associated with hydrogen atoms. The simulations were run using a leap-frog integrator with a 2 fs integration time step. The ShakeH algorithm in NAMD was applied for the water molecule constraints. A 310 K temperature was maintained using the Nóse-Hoover thermostat with 1.0 ps time constant and 1 atm pressure was maintained using isotropic coupling to the Parrinello-Rahman barostat with time constant of 5.0 ps.^111,112^ The long-range electrostatic interactions were calculated using the particle mesh Ewald method^113^ with a cut-off of 1.2 nm and a fourth-order (cubic) interpolation. The simulations were performed under an NPT ensemble with a Langevin thermostat and a Nosé–Hoover Langevin piston at 310 K and 1 atm. The damping coefficient (gamma) of the Langevin thermostat was 1/ps. In NAMD, the Nosé–Hoover Langevin piston method is a combination of the Nosé– Hoover constant pressure method^114^ and piston fluctuation control implemented using Langevin dynamics.^115^ An NPT production simulation was run on equilibrated structures for 1µs keeping the temperature at 310 K and a constant pressure (1 atm).

### Mutational Scanning of Binding Interactions in the RBD Complexes with Antibodies

We conducted a comprehensive mutational scanning analysis of the binding epitope residues. This approach systematically evaluated the effects of mutations on protein stability and binding free energy, providing insights into the structural and energetic determinants of RBD-antibody interactions. Each binding epitope residue in the RBD-antibody complexes was systematically mutated using all possible amino acid substitutions. The corresponding changes in protein stability and binding free energy were computed using the BeAtMuSiC approach.^116–119^ This method relies on statistical potential that describe pairwise inter-residue distances, backbone torsion angles, and solvent accessibility.^120^ The BeAtMuSiC approach evaluates the impact of mutations on both the strength of interactions at the protein-protein interface and the overall stability of the complex using statistical energy functions for ΔΔ*G* estimation, derived from the Boltzmann law which relates the frequency of occurrence of a structural pattern to its free energy.

BeAtMuSiC identifies a residue as part of the protein-protein interface if its solvent accessibility in the complex is at least 5% lower than its solvent accessibility in the individual protein partner(s).

The binding free energy of a protein-protein complex is expressed as the difference between the folding free energy of the complex and the folding free energies of the individual binding partners:

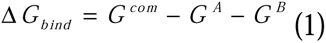

The change of the binding energy due to a mutation was calculated then as

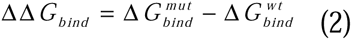

We also employed SAAMBE-3D machine learning-based predictor that utilizes knowledge-based features representing the physical environment surrounding the mutation site.^121,122^ Due to computational efficiency of BeAtMuSiC and SAAMBE-3D that enable large-scale computational mutagenesis experiments, we leveraged rapid calculations to compute ensemble-averaged binding free energy changes based on consensus estimates from BeAtMuSiC and SAAMBE-3D and using equilibrium samples from MD simulation trajectories. The binding free energy changes were averaged over 10,000 equilibrium samples for each system studied. We used 1000 ns of equilibrated trajectory data for each system, with snapshots collected at 100 ps intervals.

### Dynamic Network Analysis

To analyze protein structures, we employed a graph-based representation where residues are modeled as network nodes, and non-covalent interactions between residue side-chains define the edges. This approach captures the spatial and functional relationships between residues, providing insights into the protein structural and dynamic properties.^123–125^ The graph-based framework allows for the integration of both structural and evolutionary information, enabling a comprehensive analysis of residue interactions. The Residue Interaction Network Generator (RING) program^126–128^ was used to generate the initial residue interaction networks from the crystal structures of the antibody-RBD protein complexes. Network graph calculations were performed using the Python package NetworkX.^129,130^ To further characterize the structural and functional importance of individual residues within the RBD–antibody interaction networks, we computed multiple residue centrality metrics, including Short Betweenness Centrality (SPC), Closeness Centrality (CCA), Residual Centrality (RCA) and Z-score-based mutational perturbation profile of SPC and RCA parameters. These metrics provide complementary views into the role of each residue in mediating communication, stability, and binding within the protein complex. SPC parameter quantifies how frequently a node (residue) appears on the shortest paths between other nodes in the network. It reflects the extent to which a residue serves as a mediator of communication across the protein structure. SPC is a measure of the influence of a node in a network based on the number of shortest paths that pass through it.

The SPC of residue *i* is defined to be the sum of the fraction of shortest paths between all pairs of residues that pass through residue *i*:

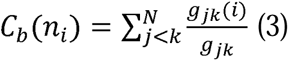

where *g_jk_* denotes the number of shortest geodesics paths connecting *j* and *k,* and *g_jk_* (*i*) is the number of shortest paths between residues *j* and *k* passing through the node *n_i_.* Residues with high occurrence in the shortest paths connecting all residue pairs have a higher betweenness values. For each node *n,* the betweenness value is normalized by the number of node pairs excluding *n* given as(*N* -1)(*N* - 2) / 2, where *N* is the total number of nodes in the connected component that node *n* belongs to.

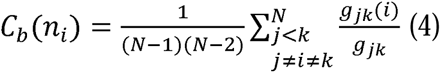

To account for differences in network size, the betweenness centrality of each residue ii was normalized by the number of node pairs excluding ii. The normalized short path betweenness of residue *i* can be expressed as : *g _jk_* is the number of shortest paths between residues *j* and k; *g _jk_* (*i*) is the fraction of these shortest paths that pass through residue *i*. Residues with high normalized betweenness centrality values were identified as key mediators of communication within the protein structure network.

### Network-Based Mutational Profiling of Allosteric Residue Interaction Networks

Residual Centrality (RCA) assesses the impact of removing a residue on the overall network connectivity by evaluating the change in the average shortest path length (ASPL) of the network. This metric helps identify residues whose removal significantly disrupts network integrity. We used this parameter to introduce mutation-based perturbations of protein residues and compute changes in the ASPL parameters averaged over all possible modifications in a given position. The change of ASPL upon mutational changes of each node is reminiscent to the calculation of residue centralities by systematically removing nodes from the network.

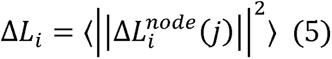

where *i* is a given site, *j* is a mutation and 〈⋯〉 denotes averaging over mutations. 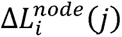 describes the change of RCA parameters upon mutation *j* in a residue node *i.* Δ*L_i_* is the average change of ASPL triggered by mutational changes in position *i*. Z-score is then calculated for each node as follows:

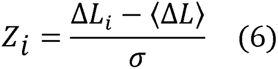

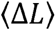 is the change of the ASPL network parameter under mutational scanning averaged over all protein residues and σ is the corresponding standard deviation. The ensemble-average Z score changes are computed from network analysis of the conformational ensembles of the antibody-RBD complexes using 1,000 snapshots of the simulation trajectory.

## Results

### Structural Analysis of the RBD Complexes with Class 4 and 5 Antibodies

We started with a focused structural analysis of the antibody-RBD complexes (Figures 1,2). Antibodies in Group F3 such as S2X35 (Figure 1A,B, Supporting Information, Table S1) and 25F9 (Figure 1C,D, Supporting Information, Table S2) employ long CDR3 loop (Complementarity Determining Region 3), particularly the heavy chain CDR3 (CDR-H3) as a crucial part in recognizing and binding to specific antigens. The light chain of an antibody has three CDRs, labeled CDRL1, CDRL2, and CDRL3. CDRL3, along with its heavy chain counterpart CDRH3, is critical for antigen binding because of its sequence variability and structural complexity. S2X35 forms a broad binding interface that extends from the ACE2-distal core RBD near Y369 position to the RBD residues D405 and R408 via close packing of its CDRH3, CDRL3 and CDRL1 loops (Figure 1A,B). 25F9 targets a similar region, with its heavy chain inserting into a hydrophobic pocket formed by conserved aromatic and aliphatic residues including RBD-Y365, F377, Y369, P384, and L387 (Figure 1C,D, Supporting Information, Table S2). Both S2X35 and 25F9 utilize extensive shape complementarity and hydrophobic stacking interactions reinforcing major determinants of their strong binding affinity and functional potency. The structural analysis of the SA55 antibody reveals that it recognizes a region encompassing residues 373–376, 404–408, 436–440, 445–446, and an extended segment spanning residues 498–508 (Figure 1E,F, Supporting Information, Table S3). At the heart of the SA55 epitope lies a highly conserved region centered around residues 436–440, which is crucial for RBD stability and proper folding. This region overlaps spatially with the ACE2-binding motif, particularly the 498–508 stretch, where several residues serve as energetic hotspots for viral attachment (Figure 1E,F, Supporting Information, Table S3). Within this segment, SA55 forms multiple contacts with conserved residues including Y501, G502, V503, G504, H505, and T508, reinforcing its ability to engage a structurally essential interface (Figure 1E,F, Supporting Information, Table S3). Among these, Y501 and H505 are especially important for ACE2 engagement, and by directly interacting with them, SA55 effectively competes with the host receptor for RBD binding. Because these residues are indispensable for viral entry, they are under strong evolutionary constraint, making them poor candidates for immune escape mutations. Consequently, the core SA55 epitope represents a structurally and functionally vulnerable site on the spike protein that is less likely to tolerate sequence variation without compromising viral fitness. SA55 also engages peripheral residues such as T376, D405, and R408, which lie at the edges of the binding interface (Figure 1E,F, Supporting Information, Table S3).

**Figure 1.**
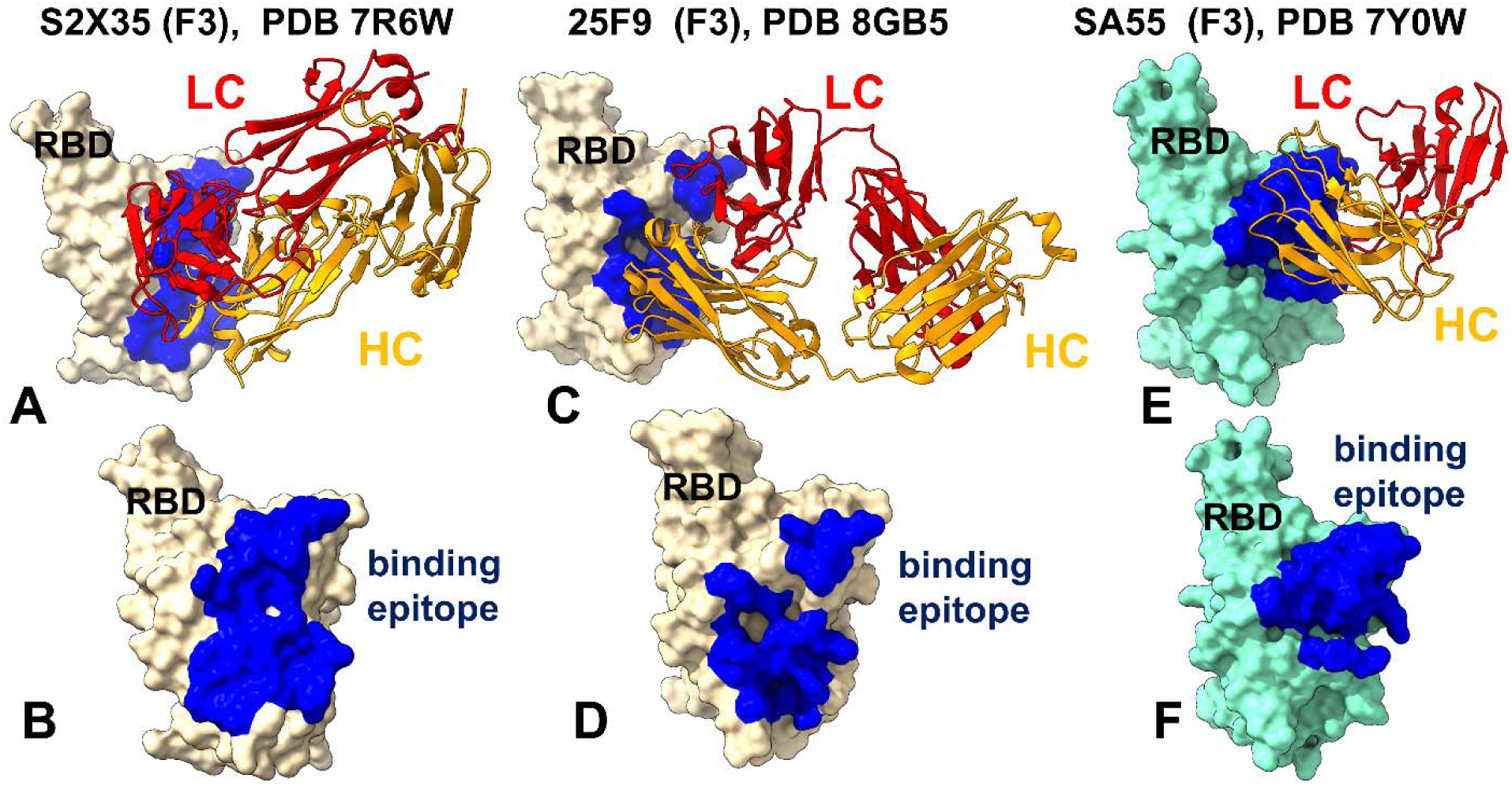
Structural organization of the RBD complexes and binding epitopes for class 4 antibodies. (A) The structure of S2X35 with RBD (pdb id 7R6W). The heavy chain in orange ribbons, the light chain in red ribbons. (B) The RBD and binding epitope footprint for S2X35. The binding epitope residues are shown in blue surface. (C) The structure of class 4 antibody 25F9 bound with RBD (pdb id 8GB5). The heavy chain in orange ribbons, the light chain in red ribbons. (D) The RBD and binding epitope footprint for 25F9. The binding epitope residues are shown in blue surface. (E) The structure of SA55 (BD55-3514) bound with BA.1 RBD (pdb id 7Y0W). The heavy chain in orange ribbons, the light chain in red ribbons. (F) The RBD and binding epitope footprint for SA55. The binding epitope residues are shown in blue surface.

**Figure 2.**
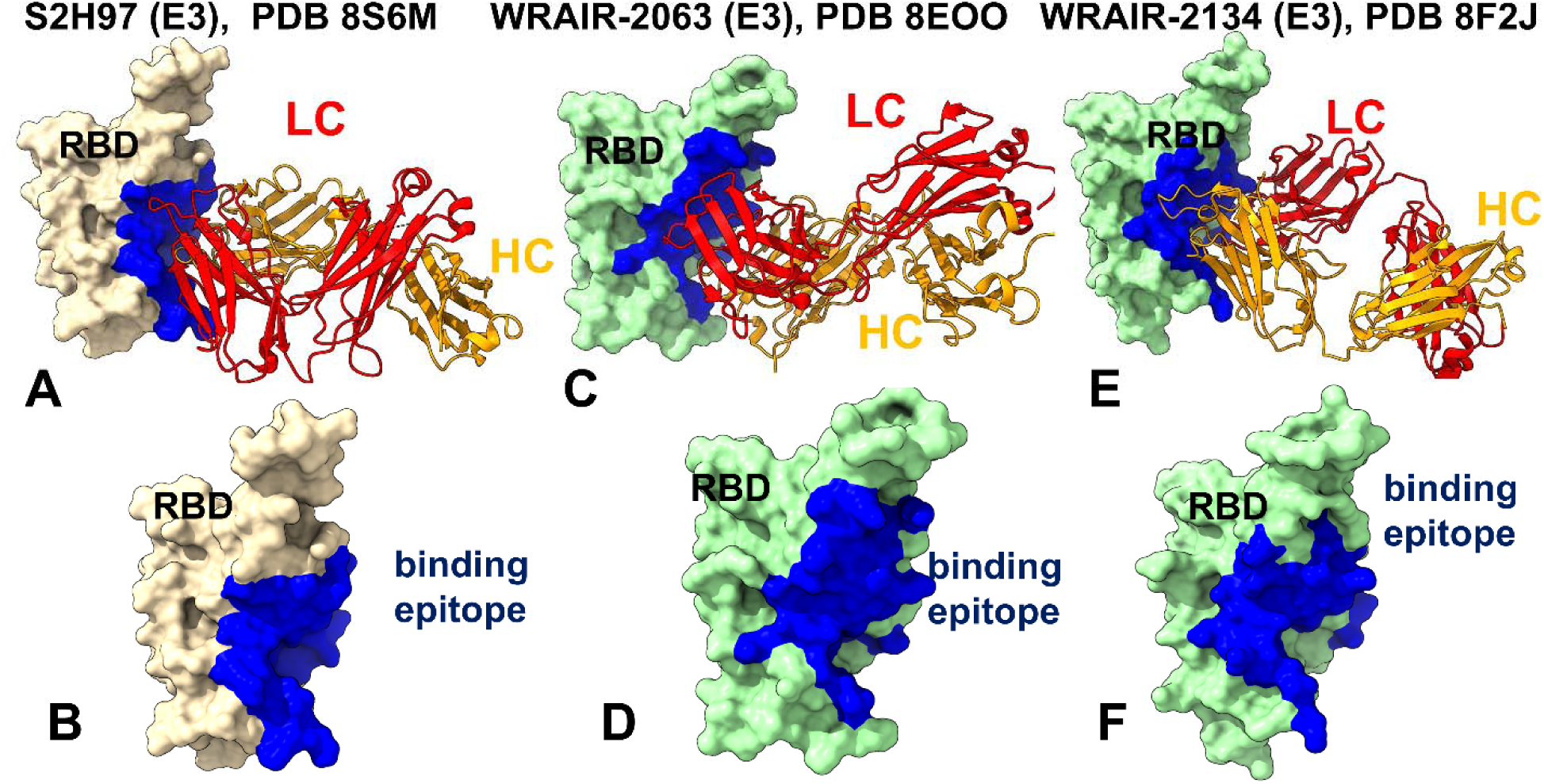
Structural organization of the RBD complexes and binding epitopes for class 5 antibodies. (A) The structure of S2H97 with RBD (pdb id 8S6M). The heavy chain in orange ribbons, the light chain in red ribbons. (B) The RBD and binding epitope footprint for S2H97. The binding epitope residues are shown in blue surface. (C) The structure of WRAIR-2063 antibody bound with RBD (pdb id 8EOO). The heavy chain in orange ribbons, the light chain in red ribbons. (D) The RBD and binding epitope footprint for WRAIR-263. The binding epitope residues are shown in blue surface. (E) The structure of WRAIR-2134 bound with RBD (pdb id 8F2J). The heavy chain in orange ribbons, the light chain in red ribbons. (F) The RBD and binding epitope footprint for WRAIR-2134. The binding epitope residues are shown in blue surface.

Despite its broad potency, SA55 is not entirely impervious to viral evolution. Mutations such as Y508H, G504S, and K440E have been shown to diminish SA55 binding, and complete escape has been observed with substitutions at V503E and G504D.^72,73^ This analysis emphasizes the dual nature of the SA55 epitope: a central, conserved core that is resistant to mutational drift due to its functional importance, flanked by more flexible peripheral regions that may accommodate escape mutations under selective pressure.

The structural analysis of the binding interfaces for class 5 antibodies S2H97 (Figure 2A,B, Supporting Information Table S4), WRAIR-2063 (Figure 2C,D, Supporting Information Table S5) and WRAIR-2134 (Figure 2E,F, Supporting Information Table S6) reveals their unique ability to target cryptic epitopes on the RBD. These epitopes are deeply buried within the spike trimer when the RBD is in the “down” conformation, requiring extensive opening of the RBD for antibody binding. The class 5 antibodies WRAIR-2134, WRAIR-2057, WRAIR-2063 and S2H97 share a closely related binding footprint binding to a highly conserved regions on the bottom and left flank of the RBD. These epitopes are distinct from those targeted by other classes of antibodies, such as class I antibodies, which often overlap with the ACE2-binding motif.

The epitopes recognized by class 5 antibodies include residues K462, E516, and L518, as well as S383, T385, and K386 (Figure 2, Supporting Information Tables S4-S6). These residues are functionally indispensable, as mutations at these positions predominantly impair protein folding, ACE2 binding, and viral infectivity.^75–77^ This cryptic epitope becomes exposed only when the RBD is in an open conformation, requiring extensive opening of the RBD for binding.^75–77^ This structural constraint limits accessibility but also renders their epitopes highly conserved across sarbecoviruses, providing a natural shield against escape mutations. According to the emerging from structural studies view, S2H97 locks the RBD into a rigid conformation, enhancing stability and strengthening binding interactions. This stabilization minimizes conformational fluctuations, making it harder for the virus to exploit flexibility for immune evasion.^75^ The structural analysis of the binding interfaces for class 5 antibodies S2H97, WRAIR-2063, and WRAIR-2134 underscores their unique mechanism of action, characterized by targeting cryptic conserved epitopes that balance immune evasion with receptor binding efficiency.

### Hierarchical Modeling of Conformational Dynamics of the RBD Complexes with Antibodies Using Coarse-Grained and Atomistic Simulations

We performed multiple CG-CABS and atomistic simulations of the RBD-antibody complexes. The root-mean-square fluctuation (RMSF) profiles provide a detailed view of the dynamic behavior of RBD residues upon antibody binding, highlighting both shared features and notable differences among the antibodies. The primary objective of this study was to investigate the dynamic and energetic contributions of RBD residues, as these residues play a pivotal role in mediating interactions with neutralizing antibodies. The key observation of this analysis is functionally relevant contrast in the conformational dynamics of the RBD when bound to class 4 antibodies (S2X35, 25F9, SA55) (Figure 3A) versus class 5 antibodies (e.g., S2H97, WRAIR-2063, WRAIR-2134). (Figure 3B). Specifically, the RMSF values of the RBD are consistently higher in complexes with class 4 antibodies compared to those with class 5 antibodies.

**Figure 3.**
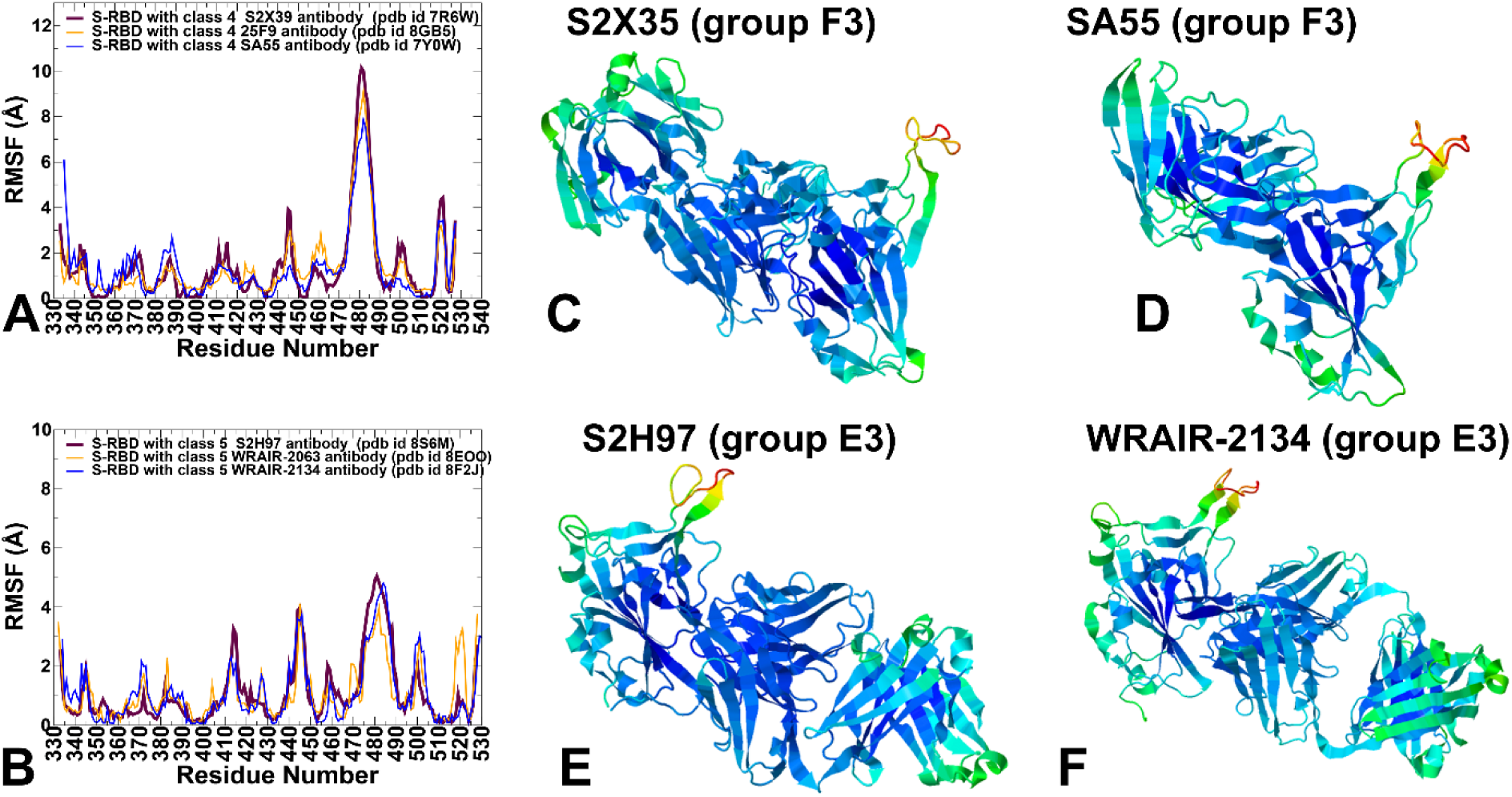
Conformational dynamics profiles obtained from CG-CABS simulations and atomistic reconstruction of the RBD-antibody complexes. (A) The RMSF profiles for the RBD residues obtained from simulations of the S-RBD complexes with class 4 antibodies : S2X35 with RBD, pdb id 7R6W (in thick maroon lines), 25F9 with RBD, pdb id 8GB5 (in orange lines), SA55 with BA.1 RBD, pdb id 7Y0W (in blue lines). (B) The RMSF profiles for the RBD residues obtained from simulations of the S-RBD complexes with class 5 antibodies : S2H97 with RBD, pdb id 8S6M (in maroon lines), WRAIRJ-2063 with RBD, pdb id 8EOO (in orange lines), WRAIR-2134 bound with RBD, pdb id 8F2J (in blue lines). Structural mapping of conformational mobility profiles along first three slow modes for the complex of class 4 (group F3) antibody S2X35 with RBD (C), class 4 (group F3) antibody SA55 with RBD (D), class 5 (group E3) antibody S2H97 with RBD (E) and class 5 (group E3) antibody WRAIR-2134 complex with RBD (F). The structures are shown in ribbons with the rigidity-to-flexibility scale colored from blue to red.

For class 4 antibodies, the RBD exhibits significantly larger RMSF values, particularly in flexible regions such as the 470–490 loop and residues 355–375 (Figure 3A). These regions are known to be highly dynamic and play a critical role in mediating interactions with both ACE2 and neutralizing antibodies. In some contrast, the RBD in complexes with class 5 antibodies shows appreciably lower RMSF values in the flexible 470-490 loop region, indicating reduced flexibility and greater stabilization (Figure 3B). The increased stabilization is wide spread and also evident in the central β-sheet and α-helices of the RBD, which exhibit minimal fluctuations across all complexes (Figure 3B). This difference could suggest potentially distinct mechanisms of interaction and stabilization between the two classes of antibodies even though both classes of antibodies target highly conserved RBD epitopes that are hidden in the closed spike form and become accessible only upon induction of a highly erected RBD form.

The RMSF analysis of RBD residues for class 4 antibodies provided some insights into the dynamic behavior of the RBD upon antibody binding. These profiles reveal both shared characteristics and particular features for each antibody (Figure 3A). The conserved structural core of the RBD include β1, β2, β3, β4, β5, β6, and β7 defined by the following residue ranges: β1: residues 354–358; β2: residues 376–379; β3: residues 394–403; β4: residues 432–437; β5: residues 452–454; β6: residues 492–494; β7: residues 507–516. This central β-sheet and α-helices exhibit low RMSF values across all class 4 antibodies, indicating minimal flexibility in these regions. This reflects the conserved structural integrity of the RBD core, which is critical for maintaining its overall stability. Residues such as 350–360, 375–380, and 394–403 show consistently low fluctuations, underscoring their role in stabilizing the RBD structure regardless of the antibody bound (Figure 3A). Residues 355–375 exhibit very moderate fluctuations, likely due to its proximity to the epitopes and their involvement in mediating interactions with the RBD. The 470-490 loop is a highly dynamic element within the RBD, playing a pivotal role in stabilizing the RBD-ACE2 interaction. It shows elevated RMSF values across class s4 antibodies, reflecting its importance in adaptive binding and immune evasion (Figure 3A). S2X35 demonstrates adaptability through increased flexibility in the 470–490 loop while 25F9 antibody tends to achieve more intermediate stabilization, balancing flexibility and rigidity to adapt to mutations while retaining binding efficacy. This is evident in regions 420–435, 450– 475, and 490–505 where fluctuations are moderately reduced (Figure 3A). The 470–490 loop showed high flexibility for 25F9 antibody but SA55 locks the RBD into a more rigid conformation, significantly reducing fluctuations in key regions (Figure 3A). This rigidity may help to prevent the RBD from adopting conformations favorable for ACE2 engagement. The 470–490 loop exhibits reduced flexibility, reflecting ability of SA55 antibody to stabilize the RBD in a conformation that sterically hinders ACE2 binding. Class 4 antibodies target cryptic epitopes on the RBD that are typically hidden in the “closed” conformation and become exposed only in the “open” state. Our results suggest that binding of class 4 antibodies to their cryptic site may allosterically promote increased flexibility in the 470–490 loop which plays a pivotal role in stabilizing the RBD-ACE2 interaction enabling escape from productive ACE2 engagement. According to this analysis class 4 antibodies S2X35, and 25F9 may prioritize adaptive flexibility in the 470-490 loop, allowing the RBD to adopt multiple conformations favorable for immune recognition but less conducive to ACE2 engagement. However, this flexibility may also make them more susceptible to escape mutations in dynamic regions of the 470–490 loop. At the same time, SA55 exemplifies a balance between considerable stabilization of the central core and part of the ACE2 binding interface (residues 500-510) leading to reduced flexibility of the RBM and potentially higher potency.^72,73^ Structural mapping of conformational mobility profiles along slow modes for class 4 S2X35 (Figure 3C) and SA55 antibodies (Figure 3D) highlighted the degree of RBD stabilization induced by antibodies and also emphasized the importance of RBM flexibility.

The RMSF profiles for class 5 antibodies revealed more significant stabilization of the RBD, particularly in regions near the ACE2-binding interface, such as residues 420–435, 450–475 and 490–505 where fluctuations are markedly reduced (Figure 3B). This stabilization reflects the ability of class 5 antibodies to lock the RBD into a more rigid conformation that is unfavorable for ACE2 engagement. However, the reduced flexibility also suggests that the antibody may restrict the RBD ability to transition between “up” and “down” conformations, which are critical for receptor-mediated viral entry. The dynamics results suggest a plausible explanation for its unique mechanism of neutralization, which involves a receptor-independent conversion of the spike protein to the post-fusion state.^75^ In this mechanism by stabilizing key regions of the RBD and reducing flexibility in dynamic 470–490 loop, S2H97 disrupts the S protein ability to adopt prefusion conformations, thereby triggering premature post-fusion conversion. While this rigid stabilization is highly effective against variants with minimal antigenic drift, it may also render class 5 antibodies become vulnerable to escape mutations at specific positions within their epitope, as they may rely more heavily on maintaining a precise conformation. Structural maps of conformational functional dynamics along the major slow modes for class 5 antibodies illustrated induction of long-range stabilization of the RBD conformation, including RBD binding interface with ACE2 (Figure 3E,F) which may force the flexible portion of the RBM loop to fluctuate only around specific open RBD-up position as any conformational changes towards down form would encounter steric hindrance with the RBD. In essence, S2H97 binding, which requires the RBD to open up and potentially impacts the mobility of the 470-490 loop, could also exert an allosteric effect that would disrupt the intrinsic conformational open-closed equilibrium of the S protein, enhance stabilization of the highly open RBD and induce receptor-independent conversion of the spike protein to the post-fusion state preventing viral entry. The proposed mechanism of S2H97 involves inducing a receptor-independent conversion of the spike protein to the post-fusion state.^75^ This process bypasses the need for ACE2 binding and directly triggers the conformational changes that lead to membrane fusion. Our results argue that the enhanced stabilization induced by class 5 antibodies in the highly flexible 470-490 region could disrupt the natural conformational landscape of the S protein involving stochastic transformations between the closed and open states. This disruption could promote the premature transition to the post-fusion conformation – a mechanism that was previously suggested as a potential driver of neutralization.^75^

The conformational dynamics of the RBD suggests differences in local and global dynamics effects induced by class 4 and 5 antibodies. Indeed, while class 4 antibodies may induce more flexibility to adapt to mutations, class 5 antibodies could rigidify the RBD in its highly specific open conformation that is incompatible with ACE2 binding. These findings underscore the importance of understanding RBD dynamics in designing broadly neutralizing antibodies capable of countering evolving viral variants. The results highlighted the adaptive role of the 470–490 loop that serves as a critical determinant of RBD flexibility, influencing both viral entry and immune recognition. On the other hand, class 5 antibodies that stabilize this loop in a specific conformation may be more effective at blocking ACE2 interactions but may be vulnerable to mutations at specific positions within their epitope.

### Mutational Profiling of Antibody-RBD Binding Interactions Interfaces Reveals Molecular Determinants of Immune Sensitivity and Emergence of Convergent Escape Hotspots

The mutational scanning analysis of F3 antibodies, S2X35, 25F9, and SA55 provides assessment of vulnerabilities to escape mutations, and resilience to viral evolution (Figure 4). S2X35 targets conserved residues in the RBD, particularly those involved in maintaining the structural integrity of the spike protein. Mutational heatmap of antibody S2X35 binding interface residues pointed to binding hotspots at RBD positions Y369, T376, F377, C379, T380, R408, V503, G504 and Y508 (Figure 4A). These findings are fully consistent with the DMS experiments according to which major escape sites for S2X35 and generally for F3 group antibodies include D405, R408, V503, G504, and Y508 (Figure 4A). The conserved hydrophobic RBD positions F377, C379 and T38 are functionally indispensable, as their mutations disrupt the structural integrity of the epitope and compromise RBD stability and antibody recognition. As a result, mutations in positions F377 and C379 are rarely observed due to requirements for RBD stability. The mutational map reflects the S2X35 binding footprint that includes packing of aromatic residues of heavy chain (Y54, Y106, G105 and L104) with Y369 on the RBD and network of interactions formed with D405, R408 and G504 by heavy chain residues Y112, Y93, W109 (Figure 4B). Indeed, these heavy chain residues on S2X35 correspond to the binding hotspots on the antibody (Figure 5A,B, Supporting Information, Figure S1A).

**Figure 4.**
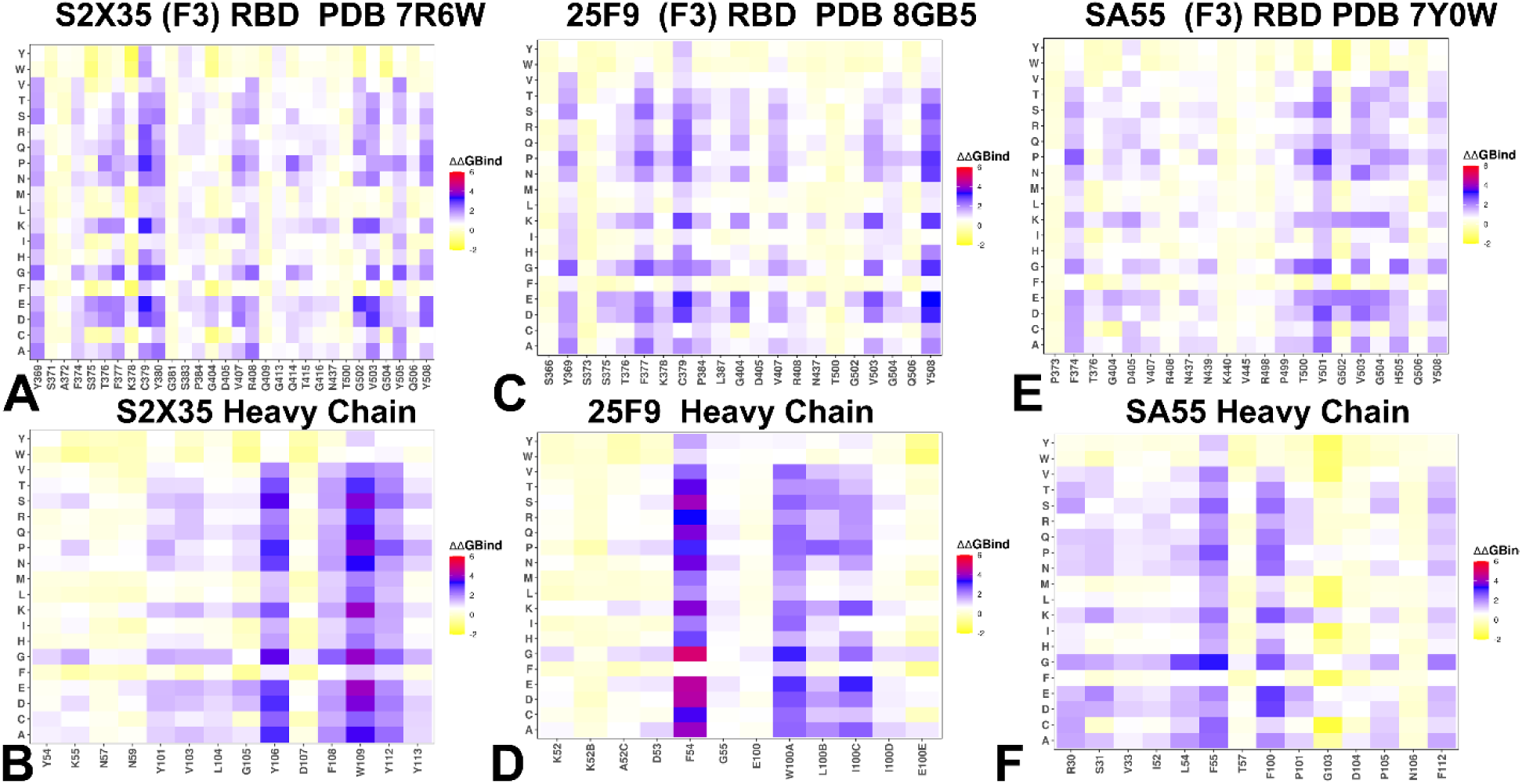
Ensemble-based dynamic mutational profiling of the RBD intermolecular interfaces in the RBD complex with class 4 S2X35 antibody (A,B), the RBD complex with class 4 25F9 antibody (C,D) and the RBD complex with class 4 SA55 antibody (E,F). The mutational scanning heatmaps are shown for the interfacial RBD residues (A,C,E) and interfacial heavy chain residues of respective class 4 antibodies (B,D,F). The heatmaps show the computed binding free energy changes for 20 single mutations of the interfacial positions. The standard errors of the mean for binding free energy changes using randomly selected 1,000 conformational samples (0.12-0.18 kcal/mol) obtained from the atomistic trajectories.

**Figure 5.**
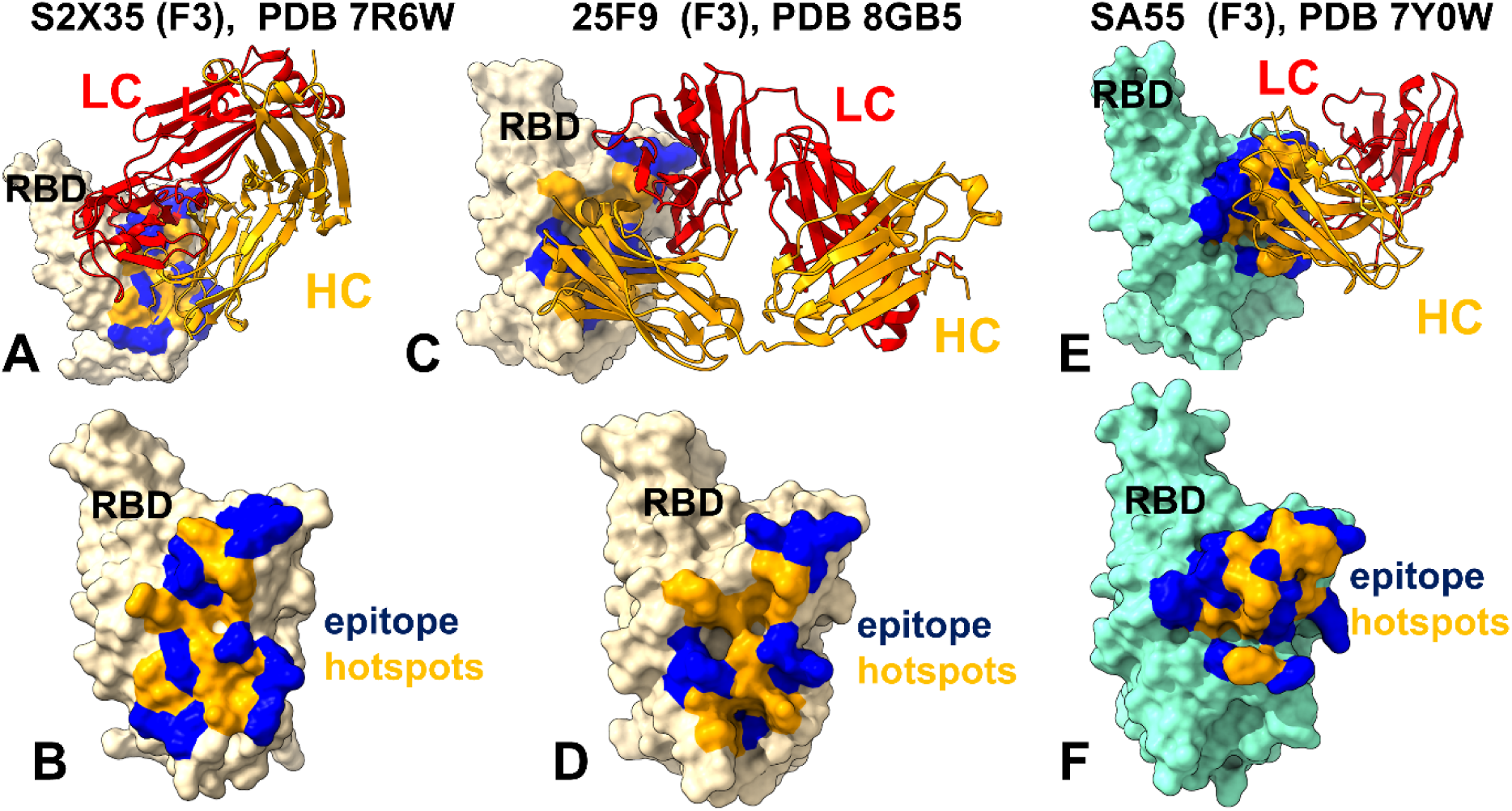
Structural mapping of binding hotspots for class 4 antibodies. (A) The structure of S2X35 with RBD (pdb id 7R6W). The heavy chain in orange ribbons, the light chain in red ribbons. (B) The RBD and binding hotspots footprint for S2X35. The binding epitope residues are shown in blue surface, and binding hotspots are in orange. (C) The structure of class 4 antibody 25F9 bound with RBD (pdb id 8GB5). (D) The RBD and binding hotspots footprint for 25F9. The binding epitope residues are shown in blue surface, and binding hotspots are in orange surface. (E) The structure of SA55 (BD55-3514) bound with BA.1 RBD (pdb id 7Y0W). (F) The RBD and binding hotspots for SA55. The binding epitope residues are shown in blue surface, and binding hotspots are in orange.

The experimental analysis showed that the strongest interactions are with D405 and R408 and escape at these position occurs via biochemically dramatic amino acid changes. ^75^ The relatively limited escape profile for S2X35 can be explained based on mutational scanning data, indicating that only D405, R408 and V503/G504 may represent potential points of vulnerability. Nonetheless, substitutions S371F, T376A, D405N, and R408S present in BA.2 could cause reduced binding which may account for the immune resistance against BA.2 variant.^131^

Mutational profiling of 25F9 antibody revealed similar vulnerabilities to mutations at residues such as Y369, T376, F377, C379, V503, and Y508 (Figure 4C). However, 25F9 demonstrates greater tolerance to mutations at positions T376, D405, and R408. The impact of T376A and R408S mutations on 25F9 binding is appreciably lower than on S2X35. Indeed, for S2X35 the destabilization energy for T376A is ΔΔG = 1.64 kcal/mol, for D405N ΔΔG = 0.74 kcal/mol and notably for R408S ΔΔG = 1.96 kcal/mol (Figure 4A). At the same time, for antibody 25F9 the destabilization energy ΔΔG values are : ΔΔG = 1.24 kcal/mol for T376A, ΔΔG= 0.32 kcal/mol for D405N, and ΔΔG= 0.77 kcal/mol for R408S indicating markedly reduced sensitivity to these mutations, particularly for R408S (Figure 4C). This relative tolerance explains a more robust neutralization activity of 25F9 against Omicron variants BA.2, BA.4/5, JN.1, KP.2, and KP.3.^91^ The hotspots on heavy chain of 25F9 are F54 and W100A residues (the crystal structure annotation) that make contacts with Y365, S366, Y369, F377, K378, C379 and L387 RBD residues (Figure 5C,D, Supporting Information, Figure S1B). This also points to the fact that strongest antibody binding anchors make interactions with conserved positions in the RBD core which are critical for RBD stability, while the interactions with mutation-vulnerable RBD sites are more moderate rendering greater immune resistance capability.

The mutational heatmap of SA55 interactions with the RBD reveals notably different pattern owing to a unique binding epitope of this broadly neutralizing antibody (Figure 4E). SA55 exhibits only mild sensitivity to mutations observed in recent variants, such as T376A (ΔΔG = 0.81 kcal/mol), D405N (ΔΔG = 0.79 kcal/mol), and R408S (ΔΔG = 0.34 kcal/mol) (Figure 4E). These modest destabilization effects reflect a limited loss in RBD stability and binding interactions, consistent with functional experiments showing that SA55 remains effective against a broad spectrum of variants, including BA.2.86, KP.2, and KP.3 subvariants.^72,73^

A significant destabilization upon mutations are observed primarily at Y501, V503, and G504 (Figure 4E). For example, mutations such as Y501D (ΔΔG = 2.74 kcal/mol), V503D (ΔΔG = 2.41 kcal/mol), and V503K (ΔΔG = 2.02 kcal/mol) may seriously impair binding affinity. However, these residues are fundamentally important for both ACE2 binding and RBD stability, making them highly constrained in terms of evolutionary flexibility. As a result, mutations at these sites are rare but impactful when they occur. The mutational heatmap of the SA55 heavy chain revealed a considerably larger number of strong binding hotspots including 30, S31, H32, L54, F55, and F112 (Figure 4F). These positions make multiple contacts with R405, R408 and most notably network of interactions with F374, T375, V503, G504, Y508. F112 hotspot position interacts strongly with T500 and Y501 (Figure 5E,F, Supporting Information Figure S1C).

These findings indicate that SA55 forms the largest interaction network among studied F3 antibodies and features strongest hotspots in T500, Y501, V503, G504 and H505 residues that are indispensable for RBD stability and ACE2 binding. Among vulnerable RBD positions are mostly V503 and G504 (Figure 4E,F) which is consistent with the DMS data showing that SA55 binding is slightly affected by Y508H; moderately affected by G504S; strongly affected by K440E; and escaped by V503E and G504D mutations.^72,73^

The mutational heatmap of the SA55 interactions showed large destabilization changes for Y501D (ΔΔG = 2.74 kcal/mol), Y501S (ΔΔG = 2.59 kcal/mol), V503D (ΔΔG = 2.41 kcal/mol), V503E (ΔΔG = 2.23 kcal/mol), and V503K mutations (ΔΔG = 2.02 kcal/mol) (Figure 4E,F). While these mutations are highly destabilizing for SA55 binding, these sites are fundamentally important for ACE2 binding and RBD functions. SA55 are sensitive to the changes on V503 and G504 but these mutations may interfere with the key RBD functions.^72,73^ As a result, evolution in these positions is highly constrained. Our mutational scanning heatmaps also showed that T376, D405 and R408 are tolerant to mutations even though these positions engage in interaction with SA55. A more detailed profiling of JN.1/KP.3 mutations against SA55 showed only small destabilization changes upon mutations T376A (ΔΔG = 0.81 kcal/mol), R403K (ΔΔG = 0.65 kcal/mol), D405N (ΔΔG = 0.79 kcal/mol), R408S (ΔΔG = 0.34 kcal/mol), L455S (ΔΔG = 0.7 kcal/mol) and F456L (ΔΔG = 0.51 kcal/mol) (Figure 4E, Supporting Information, Figure S2). These changes reflect a mild loss in the RBD stability and binding interactions, which is consistent with functional experiments showing group F3 SA55 are not sensitive to the D405N and R408S mutations of BA.2 making SA55 effective against a broad spectrum of recent variants from BA.2.86 to KP.2 and KP.3. These binding epitope sites are all located at the periphery of SA55 epitope (Figure 5E,F, Supporting Information, Figure S1C). As a result, this may rationalize the experimental observations that SA55 neutralization against BA.2/BA.3/BA.2.12.1/BA.4/BA.5 can be only moderately reduced compared with BA.1.^72,73^ In summary, the unique binding site and footprint of SA55, its focus on conserved viral regions, and its resistance to common escape mutations contribute to its ability to neutralize new variants while other group F3 antibodies may be less effective due to targeting more variable epitopes. The mutational scanning data underscores key differences in the escape profiles of F3 antibodies: S2X35 is sensitive to steric hindrance caused by mutations at residues G504 highlighting its reliance on precise structural interactions. 25F9 demonstrates better resilience to mutations, particularly at T376, D405 and R408 enabling it to maintain efficacy against diverse variants. SA55 shows high sensitivity to mutations at functionally critical residues V503 and G504, but its broader tolerance to other mutations may allow to neutralize recent variants effectively.

Class 5 antibodies, S2H97, WRAIR-2063, and WRAIR-21334, recognize highly conserved regions on the bottom and left flank of the RBD (Figure 2). Mutational scanning of the RBD binding interfaces with these antibodies generally revealed common binding hotspots K462, E516, and L518, as well as S383, T385, and K386 (Figure 6). These results are consistent with the DMS data showing that class 5 (group E3) S2H97 binds to highly conserved regions on the bottom of the RBD, interacting mainly with K462/E516/L518 and S383/T385/K386.^69^ Class 5 antibodies target cryptic epitopes that are less accessible and more conserved, providing a natural shield against escape mutations. Mutational scanning revealed that mutations at key positions K462, P463, and F464 result in significant destabilization manifested in decreased protein expression. In particular, mutations at Y396, F464 and L518 of S2H947 binding interface disrupt hydrophobic interactions critical for RBD stability, making such substitutions evolutionarily unfavorable (Figure 6A,B). These residues are functionally indispensable, as mutations at these positions predominantly impair protein folding, ACE2 binding, and viral infectivity.^69^ Similarly, L518 plays a pivotal role in maintaining RBD stability, making it less prone to mutational escape despite its importance in antibody recognition. Mutational scanning of S2H97 heavy chain showed consistent hotspots in positions W100, H102, Y103 (Figure 6B) that interact with L518, P426, D428, F429, P463, F464, F515 RBD sites (Figure 7A,B, Supporting Information, Figure S3A). Hence, class 5 epitopes are highly conserved during viral evolution, and antibodies targeting class 5 epitope, represented by S2H97, are not materially affected by Omicron mutations.^132^ Functional studies confirmed that S2H97 showed similar efficacy against all Omicron BA.2, BA.3, BA.4/5 subvariants but required high concentrations for efficient neutralization.^133^ While some studies indicate that S2H97 and other class 5 antibodies targeting similar sites retained relatively weak or moderate neutralizing activities against Omicron subvariants carrying R346T/K^25^, other studies suggested that S2H97 retained robust neutralizing activity against Omicron through recognition of the highly conserved cryptic site^75^ thus the impact of convergent mutations on S2H97 binding might be fairly moderate.

**Figure 6.**
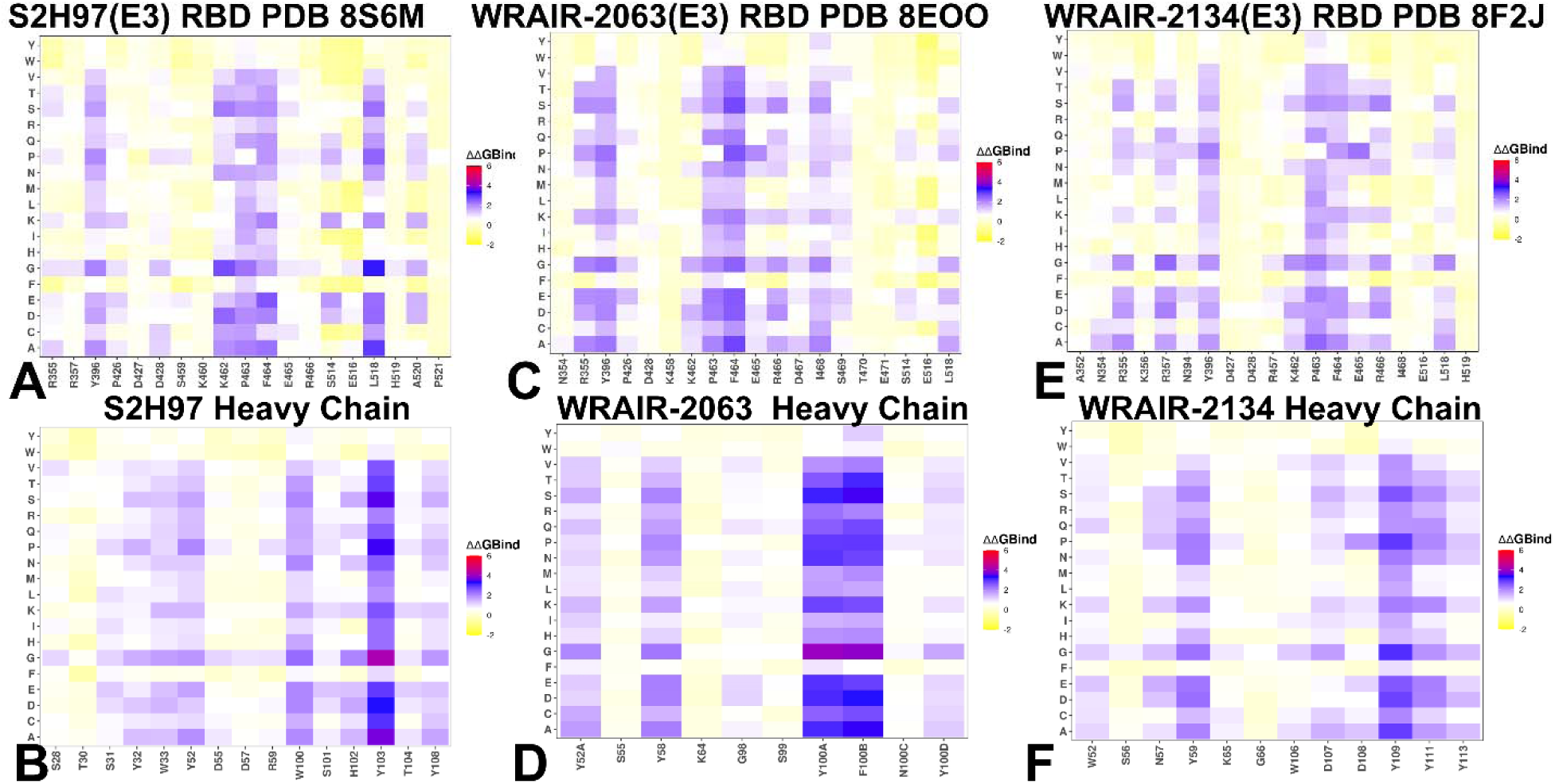
Ensemble-based dynamic mutational profiling of the RBD intermolecular interfaces in the RBD complex with class 5 S2H97 antibody (A,B), the RBD complex with class 5 antibody WRAR-2063 (C,D) and the RBD complex with class 5 antibody WRAIR-2134 (E,F). The mutational scanning heatmaps are shown for the interfacial RBD residues (A,C,E) and interfacial heavy chain residues of respective class 5 antibodies (B,D,F). The heatmaps show the computed binding free energy changes for 20 single mutations of the interfacial positions. The standard errors of the mean for binding free energy changes using randomly selected 1,000 conformational samples (0.08-0.15 kcal/mol) obtained from the atomistic trajectories.

**Figure 7.**
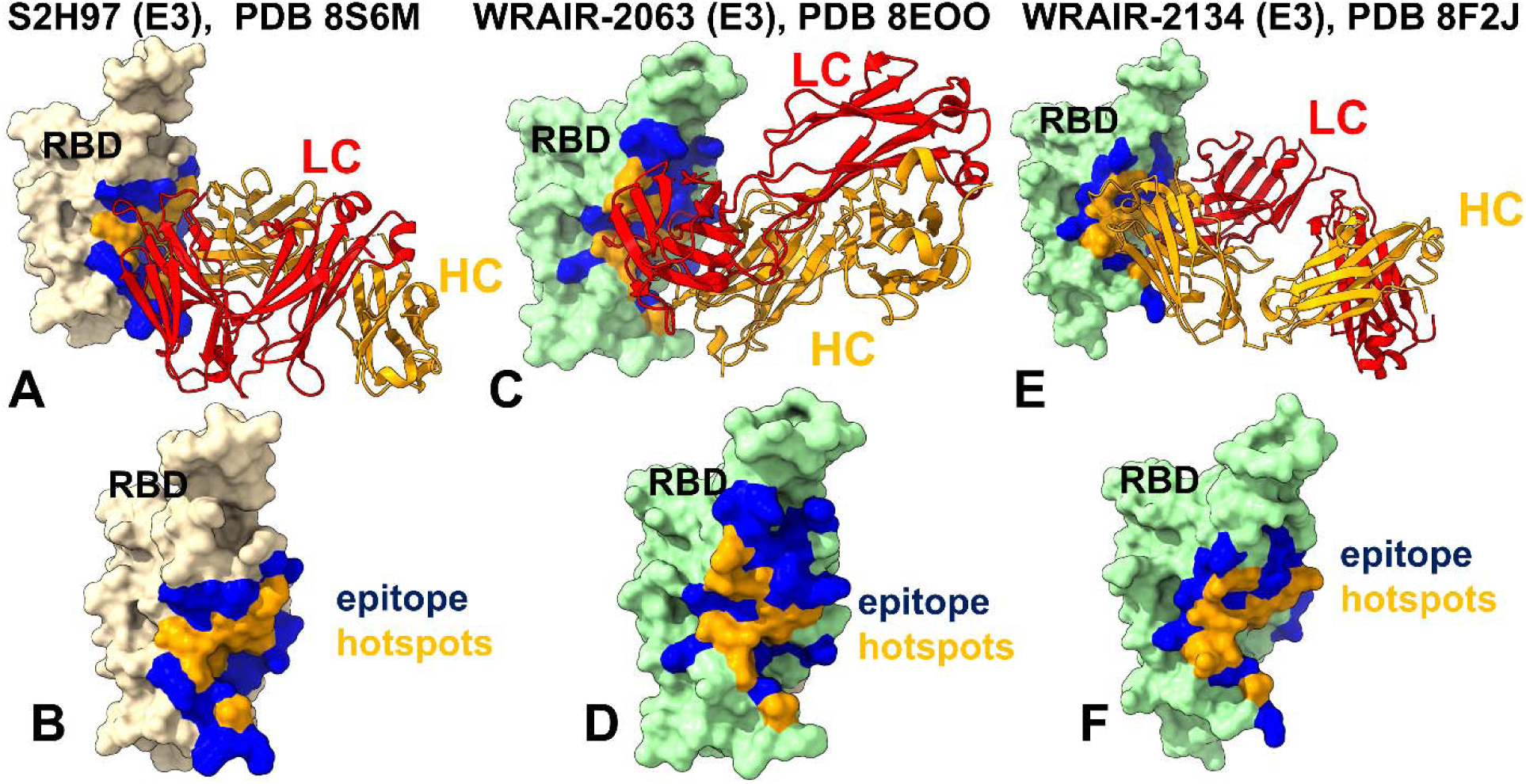
Structural mapping of binding hotspots for class 5 antibodies. (A) The structure of class 5 S2H97 with RBD (pdb id 8S6M). The heavy chain in orange ribbons, the light chain in red ribbons. (B) The RBD and binding hotspots footprint for S2H97. The binding epitope residues are shown in blue surface, and binding hotspots are in orange. (C) The structure of class 5 antibody WRAIR-2063 bound with RBD (pdb id 8EOO). (D) The RBD and binding hotspots footprint for WRAIR-2063. The binding epitope residues are shown in blue surface, and binding hotspots are in orange surface. (E) The structure of class 5 antibody WRAIR-2134 bound to RBD (pdb id 8F2J). (F) The RBD and binding hotspots for WRAIR-2134. The binding epitope residues are shown in blue surface, and binding hotspots are in orange.

Mutational scanning heatmap for class 5 WRAIR-2063 antibody showed a slightly broader range of binding hotspots at R355, Y396, P463, F464, D467, I468 and L518 positions (Figure 6C,D). The heavy chain hotspots L97, S99, F100 form strong packing hydrophobic contacts with the RBD F464 and L518 (Figure 7C,D, Supporting Information, Figure S3B). A similar mutational heatmap and binding hotspots were identified for the binding interface residues of WRAIR-2134 (Figure 6E). The binding hotspots on the antibody are concentrated on hydrophobic heavy chain sites W106, D107, D108, Y109, and Y111 (Figure 6F). These positions interact with R355, K356, Y396, P463 and F464 on the RBD (Figure 7E,F, Supporting Information, Figure S3C).

As the class 5 cryptic binding region is deeply buried within the S trimer, S2H97 can only bind when the RBD adopts a very special wide open conformation. It was argued that owing to this mechanism, neutralizing activities of class 5 (group E3) antibodies are relatively modest.^69^ Mutational scanning analysis of class 5 antibodies recapitulated the DMS experiments suggesting that critical for binding sites include R357, T393, Y396, D428, K462, S514, E516 and L518 that are all conserved in most sarbecoviruses and therefore resulting in a great neutralization breadth.^69^ To summarize, the mutational scanning analysis of class5 antibodies highlights the balance between epitope conservation, structural constraints, and viral evolution. These findings underscore the evolutionary constraints imposed by the structural and functional roles of the targeted residues, explaining why class 5 antibodies show robust resistance to escape mutations compared to antibodies targeting more exposed epitopes. In contrast, class 4 antibodies targeting different cryptic epitopes and ACE2-bnding portions of the RBD often exhibit higher potency but can be somewhat more vulnerable to a narrow range of escape mutations due to their reliance on evolutionary more vulnerable residues.

### Probing Antibody-Induced Allosteric Mechanisms of Binding Using Dynamic Network Analysis of Conformational Ensembles for Antibody-RBD Complexes

We used the ensemble-based network centrality analysis and the network-based mutational profiling of allosteric residue propensities that are computed using the topological network parameters particularly the SPC and Z-Score of the ASPL in the network where we compute changes in the metrics averaged over all possible modifications in a given position for all RBD residues in the complex (Figures 8,9). Through ensemble-based averaging over mutation-induced changes in the network parameters we identify positions in which mutations on average cause network changes. Allosteric hotspots are identified as residues in which mutations incur significant perturbations of the global residue interaction network that disrupt the network connectivity and cause a significant impairment of global network communications and compromise signaling. It is worth noting that allosteric effect of antibody binding refers to the process where binding at cryptic sites can induce long-range dynamic changes (enhancing flexibility or inducing greater RBD stabilization) and therefore affect the RBD binding interface with ACE2. The antibody-induced conformational changes in the RBD dynamics can contribute to the neutralization of the virus by interfering with the binding of ACE2 to the RBD, thus hindering viral entry into the host cell.

**Figure 8.**
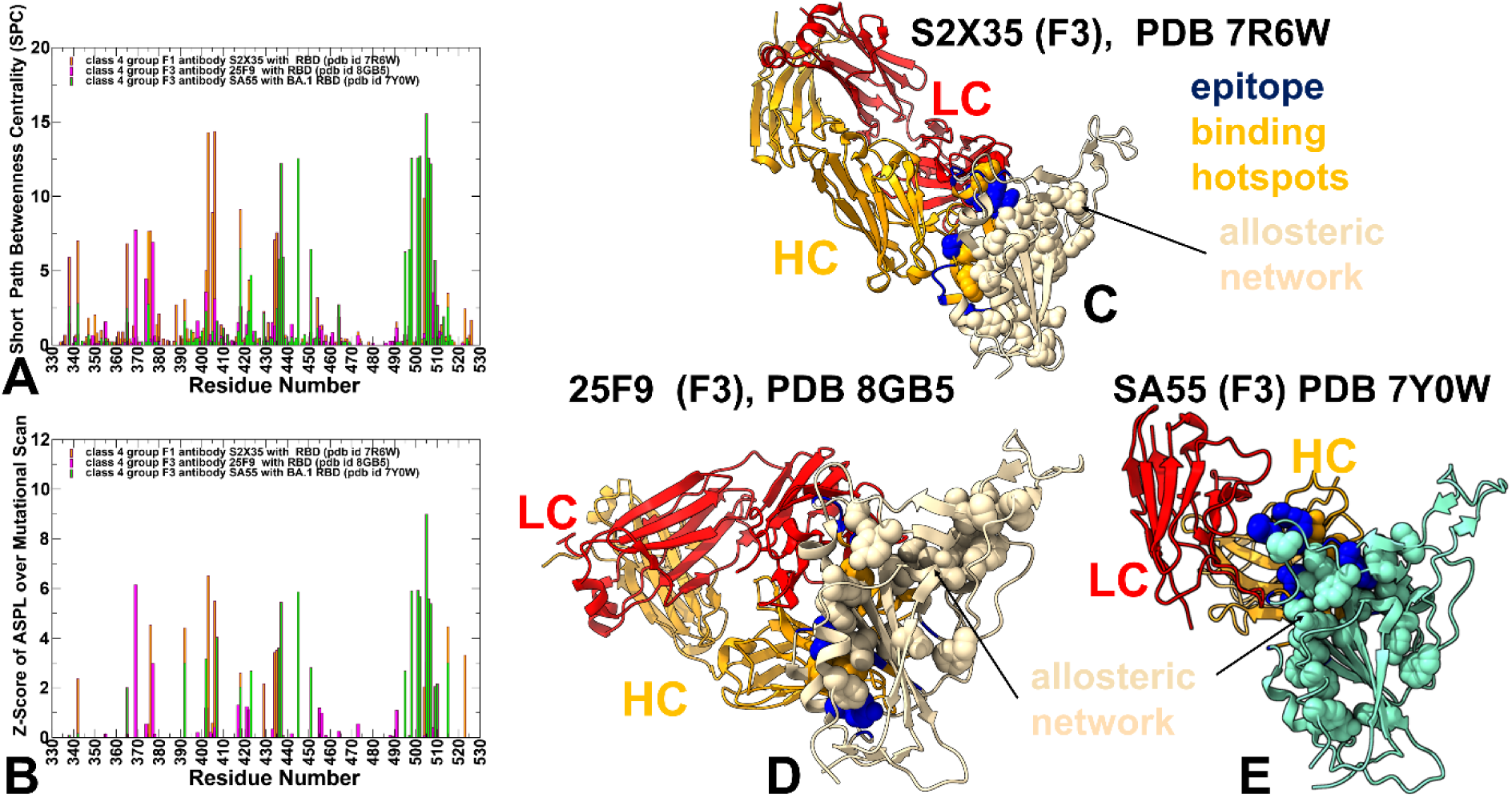
The ensemble-averaged SPC centrality (A) and the average Z-score of ASPL over mutational scan (B) for the RBD residues for the class 4 antibody complexes : S2X35 with RBD, pdb id 7R6W (in orange filled bars), 25F9 with RBD (in magenta filled bars) and SA55 with BA.1 RBD (in green filled bars). (C) Structural mapping of allosteric network centers for class 4 S2X35 with RBD. (D) Structural mapping of allosteric network sites for class 4 25F9 antibody with RBD. (E) Structural mapping of allosteric network sites for class 4 SA55 antibody with RBD.

**Figure 9.**
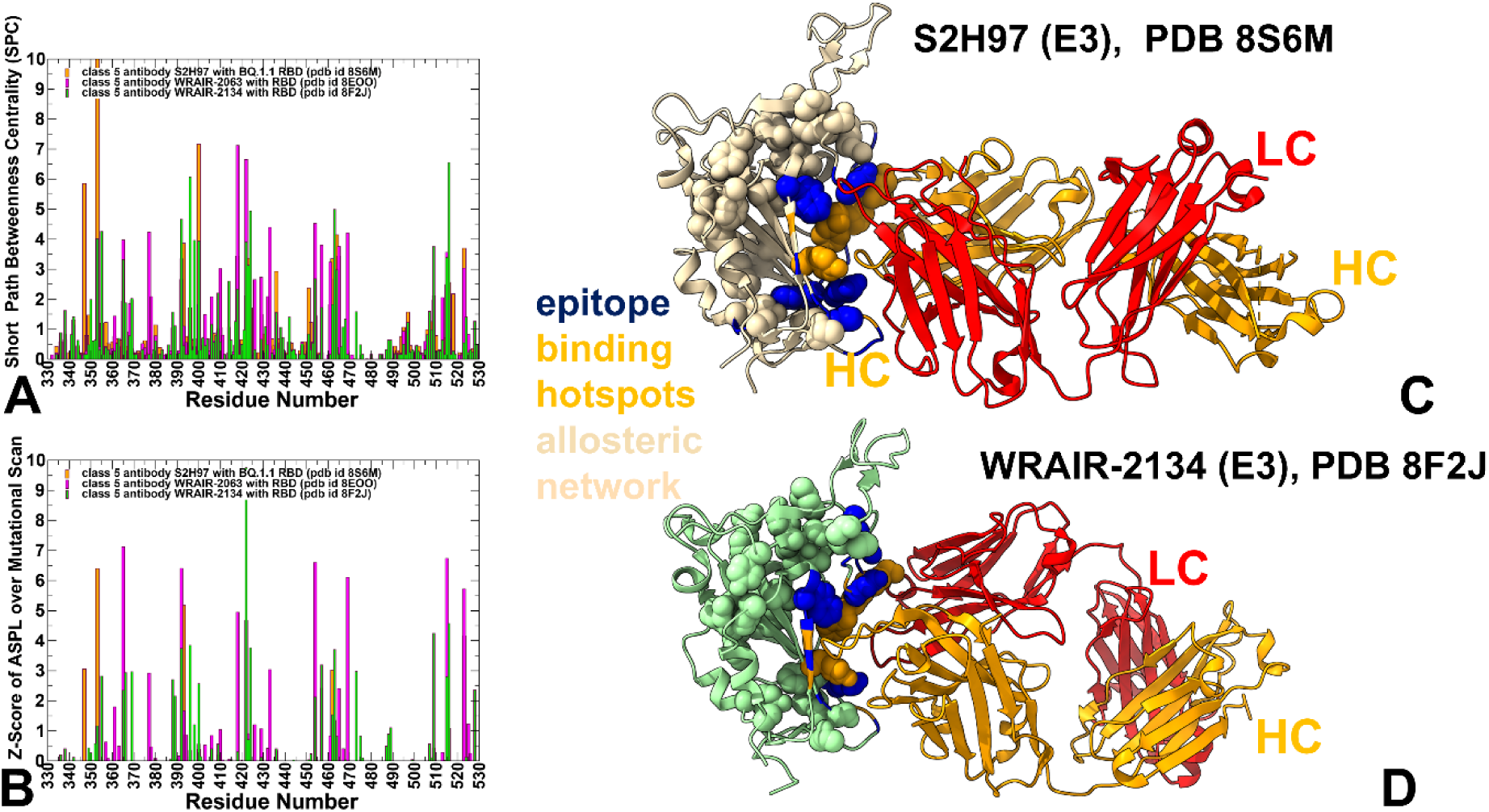
The ensemble-averaged SPC centrality (A) and the average Z-score of ASPL over mutational scan (B) for the RBD residues for the class 5 antibody complexes : S2H97 with RBD, pdb id 8S6M (in orange filled bars), WRAIR-2063 with RBD (in magenta filled bars) and WRAIR-2134 with RBD (in green filled bars). (C) Structural mapping of allosteric network centers for class 5 S297 antibody with RBD. (D) Structural mapping of allosteric network sites for class 5 WRAIR-2134 with RBD. The heavy chain in orange ribbons, the light chain in red ribbons. The binding epitope residues are shown in blue surface, and binding hotspots are in orange and allosteric residue interaction network with high SPC values in wheat-colored spheres.

We first analyzed the distribution of the SPC and Z-score ASPL parameter for class 4 antibodies S2X35, 25F9 and SA55 (Figure 8A,B). A comparative analysis showed both preservation of major allosteric positions in the RBD but also specific pattern of high centrality sites for each complex. For S2X35, we found that RBD positions E406, R403, G504, D405 and Y508 along with core sites T376, I434, A435 and W436 comprise the highest SPC and Z-score ASPLO positions. This analysis suggested that allosteric centers include some of the binding interface RBD hotspots along with important RBD core positions responsible for RBD stability (W436, I374). These sites may form a group of allosteric centers in which mutations and structural perturbations can cause significant detrimental effect on allosteric communications and are attributed to potential allosteric binding hotspots. For 25F9, the high SPC positions include Y369, F377, W436, I374, Y508, E406, I418 (Figure 8A), Mutational profiling of Z-ASPL for class 4 25F9 also revealed important role of other RBD residues Y421, L455 and F456 which are important convergent evolution positions emerged in recent Omicron variants (Figure 8A,B). Structural mapping of the high centrality network centers in the S2X35-RBD complex highlighted a potential network of mediating sites that expands from the binding interface towards the RBD regions that surrounds the conserved β-sheet core (Figure 8C). In essence, the conserved β-sheet framework of the RBD acts as a stable base from which the allosteric network extends. Notably for S2X35 the allosteric RBD network is weakly coupled to stochastic movements of 470-490 loop of the RBM which reflects the increased mobility of this region in the complexes. Hence, allosteric and dynamic analyses confluence to suggest that binding of S2X35 enables the RBM to adopt a broad ensemble of open conformations that are less conducive to productive ACE2 engagement. This adaptability allows the virus to evade immune recognition while reducing the likelihood of successful viral entry. At the same time, structural mapping of allosteric centers for 25F9 showed a network connecting the antibody binding interface with the RBM region (Figure 8D). This suggests that 25F9 may induce long-range stabilization effect on the RBM region and induce stronger binding and neutralization which is in fact consistent with the experiments. In this classification, S2X35 may be a weaker antibody that may induce long-range destabilization and increased RBM dynamics.^89^ Strikingly, a considerably different pattern of allosteric hotspots is revealed by the SPC and Z-ASPL distributions for SA55 antibody (Figure 8A,B). The distributions for both network parameters prioritized major allosteric centers at key RBD positions H505, Y501, R498, G502, K444, V445 as well as I418, W436, N437, Y453, L455. Importantly, the key allosteric centers form a subnetwork that strongly links the conserved sites at the SA55-RBD binding interface with important RBD residues involved in Omicron mutations such as K444, V445, Y453 and L455. Structural mapping of the allosteric sites for SA55-RBD complex highlighted the expansion of the allosteric network towards the RBD-ACE2 binding site, emphasizing antibody-induced stabilization of the RBM regio (Figure 8E). As a result, allosteric effect of SA55 may involve considerable stabilization of the central core and the ACE2 binding interface (residues 500-510) leading to reduced flexibility of the RBM and locking a specific RBD-up conformation that could sterically hinder productive ACE2 binding. We argue that mechanism of SA55 neutralization may be governed by confluence of binding and allosteric effects where strong binding to conserved RBD epitope is also linked with allosterically induced stabilization of the RBM region and interfering with ACE2 binding. It is worth stressing that we performed structural mapping of top 20 high SPC/Z-score consensus RBD positions. Importantly, these sites are not confined to specific RBD region, and the network is wide-spread beyond the antibody binding interface region and includes RBD core residues surrounding the central β-sheet core (Figure 8C-E).

The heavy chain in orange ribbons, the light chain in red ribbons. The binding epitope residues are shown in blue surface, and binding hotspots are in orange and allosteric residue interaction network with high SPC values depicted in wheat-colored spheres.

To summarize, our dynamic network analysis of antibody-bound RBD ensembles reveals that Class 4 neutralizing antibodies exert their inhibitory effects not only through direct binding but also via modulation of RBD conformational dynamics. Importantly, our results demonstrate that antibody binding at cryptic or conserved epitopes can induce significant allosteric changes in the RBD, which may either enhance its flexibility or impose structural stabilization, ultimately influencing the accessibility of the ACE2-binding interface. This allosteric modulation provides an additional layer of neutralization mechanism beyond simple steric hindrance, offering insights into how certain antibodies maintain broad efficacy despite ongoing viral evolution. Despite targeting overlapping regions on the RBD, each antibody exhibits a distinct pattern of high centrality residues, suggesting unique allosteric fingerprints that correlate with differences in neutralization potency and mutation tolerance. These observations suggest that while all three antibodies engage the RBD at conserved structural elements—such as the central β-sheet core — they differ in the spatial organization of their allosteric networks, which likely underlies their distinct functional outcomes.

Most strikingly, SA55 exhibits a highly centralized and conserved allosteric signature, with top-ranked residues forming a subnetwork that links the conserved antibody-binding interface to key ACE2-contacting residues such as Y501, H505, and G504, as well as emerging Omicron mutational sites K444, V445, Y453, and L455. Notably, SA55 appears to stabilize the RBD core and lock the RBM in a constrained conformation, effectively reducing its flexibility and impairing the ability of the RBD to undergo the dynamic transitions required for ACE2 recognition. This dual mechanism—comprising both high-affinity binding to a structurally critical region and allosterically driven stabilization of the RBD —likely contributes to SA55 remarkable breadth and resistance to escape mutations. The distribution of the network parameters SPC and Z-score of ASPL for class 5 antibodies showed a much denser and broader allocation of allosteric centers (Figure 9A,B). This may indicate the emergence of a larger and broadly distributed allosteric network that interlinks different RBD regions and can contribute to long-range stabilization of the RBD (Figure 9C,D). For S2H97, the high centrality positions include W353, F400, F347, F464, E465, I418, Y423, W436, F392, T393, F515, L518, T523 (Figure 9A,B). A similar group of RBD core residues corresponded to allosteric network centers for WRAIR-2063 antibody featuring F392, T393, F418, Y421, N422, Y423 F377, E365, L461, P463 Several other RBD residues R454,L455 (β5: residues 452–454) and residues F515, L518 (β7: residues 507–516) were among top allosteric centers for WRAIR-2063. While the major allosteric hotspots corresponding to conserved RBD core are preserved among all class 5 antibodies, for WRAIR-2134 complex with RBD, the high centrality allosteric sites additionally included RBD positions Y453, R454, L455 R457, Y473 that are located away from the direct antibody interface and correspond to important stabilization centers of RBD binding with the host receptor ACE (Figure 9A,B).

Structural mapping of top allosteric centers in the class 5 antibody complexes showed signatures of a broad allosteric network that encompassed the key elements of the RBD core including the central β-sheet as well as residues from the RBD regions proximally to ACE2 binding site (Figure 9C,D). The identified allosteric centers link the cryptic binding interface with the conserved elements of the RBD core and hydrophobic RBD regions near the ACE2 binding interface. Mutations of these residues may affect allosteric communications between the binding sites and compromise structural integrity and stability of the RBD. Indeed, the experiments showed that substitutions in these positions resulted in decreased protein expression, ACE2 binding, and viral infectivity, highlighting functional and structural constraints imposed on the allosteric centers.^76,77, 132^

Structural analysis of the projected allosteric centers highlighted the dense interconnectivity between the cryptic binding epitope, the RBD central core and the RBD region anchoring the RBM motif (Figure 9C,D). The architecture of the allosteric network suggests that class 5 antibodies induce long-range conformational stabilization of the RBD and S1 subunit which is a typical signature of strong neutralization and high binding affinity.^76,89^ Indeed, these class 5 antibodies demonstrate broad neutralization against known VOC’s with high-affinity binding to RBDs.^76^ According to the allosteric network analysis class 5 antibodies may induce considerable stabilization of the RBD in the specific open conformation which may trigger conformational changes in the S trimer that impact accessibility of other cryptic sites.^75^ As a result, class 5 antibodies by stabilizing the RBD in a specific conformation can disrupt its ability to transition between functional states required for viral entry and potentially promoting premature transition to the post-fusion conformation. The results point to a synergy of binding and allosteric effects where targeting conserved RBD epitope with strong local contacts is also linked with allosterically induced stabilization of the remote RBD regions and sequestering of a specific conformational state that compromises spike equilibrium and ACE2 engagement.

The observed interplay between binding-induced allosteric changes and RBD conformational equilibrium offers a mechanistic explanation for the variable neutralization profiles of these antibodies. Class 4 antibodies appear to modulate the intrinsic flexibility of the RBD, shifting the balance between closed and open states. The increased RBM mobility induced by S2X35 may allow explore alternative conformations, enabling partial immune evasion while still limiting ACE2 binding. Conversely, enhanced RBM stabilization, as with 25F9 and especially SA55, restricts conformational freedom, impairing ACE2 access and resulting in robust neutralization. We propose that the most effective neutralization by SA55 arises from a synergistic combination of strong epitope binding and allosterically mediated stabilization of the RBD. Specifically, SA55 appears to lock the RBD into a conformation that thereby sterically occluding host receptor binding. Class 5 antibodies form interactions with residues that are spatially distributed across multiple RBD subdomains, including the NTD-RBD interface, the β-sheet core, and the ACE2-binding motif. Dynamic network analysis revealed that these antibodies elicit a dense and broadly distributed allosteric network, connecting distant regions of the RBD and reinforcing overall structural rigidity. This long-range allosteric stabilization locks the RBD in a specific conformation, thereby impairing its ability to undergo the necessary transitions for viral entry. A unifying theme across both Class 4 and Class 5 antibodies is the synergistic relationship between epitope conservation and allosteric control. While direct binding to conserved residues ensures broad reactivity, the allosteric propagation of dynamic constraints enhances neutralization by modulating the functional conformational ensemble of the RBD.

Despite their shared emphasis on conserved RBD elements, mutational profiling of binding and allostery revealed that certain substitutions can indeed compromise network integrity. Interestingly, many of these escape-prone residues are located at topologically sensitive positions, where small structural perturbations can lead to widespread network disruption. Yet, in many cases, such mutations incur a fitness penalty, as evidenced by reduced protein expression and ACE2 binding affinity. This suggests that evolutionary paths toward complete escape are limited, particularly for antibodies that engage deeply conserved structural motifs. Moreover, the dense allosteric network formed by class 4 SA55 and Class 5 antibodies implies that mutations at any single node may not be sufficient to fully escape neutralization, due to the presence of redundant or compensatory communication routes which may underpin the remarkable breadth and durability of these antibodies.

Strikingly, quite similar to pioneering experimental studies by Lehner and colleagues mapping allosteric landscapes in various proteins^132–135^, we found that while the peaks of highest SPC/Z-score positions are located near the antibody binding interface, many conserved high centrality sites are situated away from the antibody binding site, and in the RBD regions surrounding the central β-sheet core. The results suggest a modular allosteric architecture with the conserved “allosteric ring” in the RBD core shared by all complexes and antibody-specific “extensions” of the allosteric network that expands towards ACE2 binding site affects the stability of the RBM regions. These results are consistent with the recently proposed generic architecture principles of allosteric interactions in proteins reported in a series of seminal studies from Lehner’s lab.^132–135^ According to these experiments, allostery primarily evolves via the gain-and-loss of peripheral extensions to a conserved allosteric core.^134,135^ In our allosteric network analysis, we found evidence that antibodies can modulate the composition of these “peripheral extensions” by specifically engaging functional RBM regions near the ACE2 binding site, modulating mobility and affecting the immune resistance responses.

## Discussion

The structural and dynamic analyses of Class 4 and Class 5 monoclonal antibodies reveal a functionally significant interplay between epitope recognition, conformational modulation of the RBD, and allosteric reprogramming of residue interaction networks. These findings provide a comprehensive atomistic view of how different antibodies engage RBD and modulate functional conformational ensemble offering insights into mechanisms of broad neutralization and resistance to immune escape mutations. Class 4 antibodies represented by S2X35, 25F9, and SA55 target conserved regions of the RBD that are accessible across multiple spike conformations. Despite overlapping epitopes, each antibody exhibits distinct impacts on RBD flexibility and mutation tolerance, underscoring the importance of not just where they bind, but how their binding propagates through the protein dynamic network. Our hierarchical modeling of conformational dynamics using CG-CABS and atomistic simulations reveals that Class 4 antibodies generally induce greater mobility in flexible regions such as the 470–490 loop and residues 355–375, compared to Class 5 antibodies. This suggests that while these antibodies recognize structurally conserved motifs, they can promote increased RBM mobility, potentially allowing the virus to explore alternative conformations that reduce ACE2 compatibility while enabling partial immune evasion. S2X35 appears to allow the broadest conformational freedom, particularly in the RBM, which may reflect an adaptive mechanism that balances immune recognition with functional constraint. In contrast, 25F9 demonstrates intermediate stabilization, retaining flexibility in dynamic regions of the 470–490 loop while preserving core interactions critical for binding. SA55, however, locks the RBD into a more rigid state, significantly reducing fluctuations in both the central β-sheet and at the ACE2-binding interface. This global stabilization likely underpins its exceptional breadth against Omicron subvariants, especially BA.2, KP.2, and KP.3, where it remains effective despite common mutations at positions such as T376A, D405N, and R408S.

Mutational scanning data further refine our understanding of these antibodies’ vulnerabilities. S2X35 shows high sensitivity to mutations at D405N, R408S, and V503E/G504D, consistent with its reliance on precise hydrophobic contacts. 25F9 engages a broader set of conserved RBD core residues, exhibits greater resilience to peripheral mutations. SA55 can tolerate many mutational changes at the periphery while remaining sensitive to substitutions at functionally essential sites Y501, V503, and G504, which serve as energetic hotspots for both ACE2 engagement and RBD folding. Mutations at these positions often result in reduced viral infectivity, reinforcing the idea that immunological pressure and functional constraints converge at key epitope regions. Our findings align closely with DMS evidence showing that SA55, despite being affected by substitutions like V503E and G504D, remains broadly effective against circulating variants due to its reliance on structurally critical and conserved RBD motifs. In contrast, Class 5 antibodies S2H97, WRAIR-2063, and WRAIR-2134 showed a distinct pattern of resilience. Our mutational scanning identified K462, F464, and L518 as central hubs in the antibody-RBD interface, consistent with DMS data indicating that these residues are rarely mutated in natural variants due to their structural importance. Class 5 antibodies S2H97, WRAIR-2063, and WRAIR-2134 exhibit a markedly different behavior. Their epitopes are deeply buried within the spike trimer and only become accessible when the RBD adopts a fully open conformation. This structural requirement imposes a unique constraint on viral evolution, as mutations in this region often compromise spike expression, ACE2 binding and viral infectivity making them poor candidates for immune escape.

Dynamic network analysis revealed that class 4 antibodies can exhibit distinct patterns of allosteric influence despite targeting overlapping regions. S2X35 induces a loosely connected network with weak coupling to the highly flexible 470–490 loop, promoting conformational diversity in the RBM and potentially enabling partial immune evasion while still impairing ACE2 engagement. In contrast, 25F9 forms a more coherent network linking its epitope to the RBM, suggesting a role in long-range stabilization and stronger neutralization. Notably, SA55 shows a highly centralized allosteric signature, with high centrality residues forming a subnetwork that connects the antibody-binding interface to critical ACE2-contacting sites like Y501, G504, and H505, as well as emerging mutational sites such as K444, V445, and L455. This dual mechanism—strong direct binding combined with allosterically induced rigidity may contribute to broad neutralization breadth of SA55 and resistance to escape mutations.

The network modeling of allostery highlights that Class 5 antibodies elicit dense and broadly distributed allosteric networks, linking cryptic binding interfaces to the RBD core and the ACE2-binding motif. This leads to significant reduction in RBD flexibility, particularly in the 470–490 loop and other dynamic elements involved in receptor engagement. Importantly, the allosteric effects are propagated through the conserved β-sheet framework, which acts as a structural backbone for long-range signaling. This mechanism provides a novel rationale to neutralization which includes direct binding effect combined with disruption of the natural conformational landscape of the RBD, thereby compromising viral entry even before ACE2 engagement. Mutational profiling confirms the evolutionary constraints acting on Class 5 epitopes. RBD positions K462, L518, and F464, which are top-ranked in network centrality metrics, show marked destabilization upon mutation, correlating with reduced protein expression and loss of neutralization potency. Mutations at these sites often incur fitness costs, reinforcing the idea that Class 5 antibodies target functionally indispensable residues which is a feature that enhances their durability across evolving variants. WRAIR-2134 extends its allosteric influence beyond direct binding sites, engaging additional stabilization centers such as Y453, R454, and Y473, which are important for anchoring the RBM. This broader footprint explains its robustness and its ability to sequester the RBD in a specific conformation incompatible with ACE2 binding. The comparative framework illustrates that Class 4 antibodies act primarily through a combination of strong local binding and moderate global stabilization, whereas Class 5 antibodies exert long-range effects, leveraging cryptic epitopes to impose widespread rigidity across the RBD and S1 subunit.

While all examined antibodies engage conserved RBD motifs, the extent to which they propagate dynamic effects varies, shaping the neutralization profile and escape vulnerability of each antibody. Despite their shared focus on conserved RBD regions, neither Class 4 nor Class 5 antibodies are entirely immune to escape. Class 4 antibodies, particularly S2X35, exhibit notable sensitivity to mutations at key binding sites such as G504 and R408, though many of these mutations still incur fitness penalties. Class 5 antibodies, due to their reliance on deeply conserved residues, face stronger evolutionary constraints. However, most of these positions are structurally irreplaceable making them poor targets for viable escape pathways. This network-based view supports a model in which the effective antibodies combine high-affinity binding to structurally critical residues, long-range allosteric effects that restrict RBD adaptability and propagation of dynamic constraints that can reshape the conformational equilibrium and determine efficacy and neutralization patterns. Antibodies SA55 and WRAIR-2134 may exemplify this paradigm, showing broad cross-reactivity and high resistance to escape mutations may be in part due to their engagement into central network hubs that are indispensable for RBD function. These results highlight the mechanistic synergy between direct binding and allosteric propagation where antibodies not only recognize conserved epitopes but also reshape the conformational landscape and operate through confluence of binding and allosteric control. The excellent agreement with DMS data reinforces the notion that integrating ensemble-based mutational scanning of binding and allostery may be deployed to assist in design of durable, broadly neutralizing antibodies that exploit evolutionary constraints and modulate dynamic communication pathways.

## Conclusion

This study reveals that broadly neutralizing monoclonal antibodies from Class 4 and Class 5 exert their effects through a combination of direct epitope recognition and dynamic modulation of RBD conformational ensembles. Despite targeting overlapping or cryptic regions, each antibody exhibits a unique pattern of interaction and allosteric influence, resulting in functionally divergent outcomes. We found that class 4 antibodies modulate the RBD conformational equilibrium in ways that either enhance flexibility (as seen with S2X35), promote intermediate stabilization (25F9), or impose stronger rigidity (SA55). The results suggested that the most efficient antibody SA55 may exemplify a dual mechanism as it binds a central, conserved region near the ACE2 interface and simultaneously stabilizes the RBD core, impairing the ability to transition into ACE2-compatible conformations. In contrast, class 5 antibodies engage deeply buried epitopes that become accessible only when the RBD adopts an open conformation. Their binding induces widespread stabilization across the RBD, particularly in flexible motifs and may trigger receptor-independent spike activation, effectively blocking viral entry. Mutational scanning confirms that escape at these sites often incurs fitness costs, reinforcing the functional constraints acting on this class of antibodies. A major finding of the allosteric network analysis of conformational ensembles is that Class 5 antibodies induce broader and more robust allosteric networks, which span multiple structural elements of the RBD and link distant functional motifs. This widespread stabilization explains their exceptional resistance to immune escape, as mutations at these topologically important nodes often lead to loss of RBD folding, decreased ACE2 binding, and reduced viral infectivity.

In contrast, Class 4 antibodies, while still engaging conserved structural elements, show more localized and variable allosteric effects, making them somewhat more vulnerable to specific escape mutation, particularly at peripheral sites V503 and G5034 of the RBD. Importantly, the ensemble-based network modeling suggested that binding and neutralization mechanisms extend beyond simple steric hindrance, involving binding-induced reprogramming of RBD long-range stabilization.

Together, these atomistic and network-level insights offer a new mechanistic framework where the effective antibodies combine local binding to conserved motifs with global dynamic reprogramming of RBD conformational ensembles, thereby enhancing breadth and durability against antigenically drifting variants. These findings may be helpful for engineering next-generation therapeutics that exploit both binding and allostery to counteract immune escape and stabilize functionally critical spike conformations.

## Author Contributions

Conceptualization, G.V.; methodology, G.V.; software, M.A., V.P., B.F., G.H., G.V.; validation, G.V.; formal analysis, G.V., M.A., G.H., investigation, G.V.; resources, G.V., M.A. and G.H.; data curation, G.V.; writing—original draft preparation, G.V.; writing—review and editing, G.V., M.A. and G.H.; visualization, V.P., B.F., G.V.; supervision, G.V.; project administration, G.V.; funding acquisition, G.V. All authors have read and agreed to the published version of the manuscript.

## Data Availability Statement

Data is fully contained within the article and Supplementary Materials. Crystal structures were obtained and downloaded from the Protein Data Bank (http://www.rcsb.org). The rendering of protein structures was done with UCSF ChimeraX package (https://www.rbvi.ucsf.edu/chimerax/) and Pymol (https://pymol.org/2/). All mutational heatmaps were produced using the developed software that is freely available at https://alshahrani.shinyapps.io/HeatMapViewerApp/.

## Acknowledgments

The authors acknowledge support from Schmid College of Science and Technology at Chapman University for providing computing resources at the Keck Center for Science and Engineering.

## References

1. Tai, W.; He, L.; Zhang, X.; Pu, J.; Voronin, D.; Jiang, S.; Zhou, Y.; Du, L. Characterization of the receptor-binding domain (RBD) of 2019 novel coronavirus: implication for development of RBD protein as a viral attachment inhibitor and vaccine. Cell. Mol. Immunol. 2020, 17, 613–620. doi: 10.1038/s41423-020-0400-4.

2. Wang, Q.; Zhang, Y.; Wu, L.; Niu, S.; Song, C.; Zhang, Z.; Lu, G.; Qiao, C.; Hu, Y.; Yuen, K. Y.; Wang, Q.; Zhou, H.; Yan, J.; Qi, J. Structural and functional basis of SARS-CoV-2 entry by using human ACE2. Cell 2020, 181, 894–904.e9. doi: 10.1016/j.cell.2020.03.045.

3. Walls, A. C.; Park, Y. J.; Tortorici, M. A.; Wall, A.; McGuire, A. T.; Veesler, D. Structure, Function, and Antigenicity of the SARS-CoV-2 Spike Glycoprotein. Cell 2020, 181, 281–292.e6. doi: 10.1016/j.cell.2020.02.058.

4. Wrapp, D.; Wang, N.; Corbett, K. S.; Goldsmith, J. A.; Hsieh, C. L.; Abiona, O.; Graham, B. S.; McLellan, J. S. Cryo-EM structure of the 2019-nCoV spike in the prefusion conformation. Science 2020, 367, 1260–1263. doi: 10.1126/science.abb2507.

5. Cai, Y.; Zhang, J.; Xiao, T.; Peng, H.; Sterling, S. M.; Walsh, R. M., Jr.; Rawson, S.; Rits-Volloch, S.; Chen, B. Distinct conformational states of SARS-CoV-2 spike protein. Science 2020, 369, 1586–1592. doi: 10.1126/science.abd4251.

6. Hsieh, C. L.; Goldsmith, J. A.; Schaub, J. M.; DiVenere, A. M.; Kuo, H. C.; Javanmardi, K.; Le, K. C.; Wrapp, D.; Lee, A. G.; Liu, Y., Chou, C.W.; Byrne, P.O.; Hjorth, C.K.; Johnson, N.V.; Ludes-Meyers J.; Nguyen, A.W.; Park, J.; Wang, N.; Amengor, D.; Lavinder, J.J.; Ippolito, G.C.; Maynard, J.A.; Finkelstein, I.J.; McLellan, J.S. Structure-based design of prefusion-stabilized SARS-CoV-2 spikes. Science 2020, 369, 1501–1505. doi: 10.1126/science.abd0826.

7. Henderson, R.; Edwards, R. J.; Mansouri, K.; Janowska, K.; Stalls, V.; Gobeil, S. M. C.; Kopp, M.; Li, D.; Parks, R.; Hsu, A. L., Borgnia, M.J.; Haynes, B.F.; Acharya, P. Controlling the SARS-CoV-2 spike glycoprotein conformation. Nat. Struct. Mol. Biol. 2020, 27, 925–933. doi: 10.1038/s41594-020-0479-4.

8. McCallum, M.; Walls, A. C.; Bowen, J. E.; Corti, D.; Veesler, D. Structure-guided covalent stabilization of coronavirus spike glycoprotein trimers in the closed conformation. Nat. Struct. Mol. Biol. 2020, 27, 942–949. doi: 10.1038/s41594-020-0483-8.

9. Xiong, X.; Qu, K.; Ciazynska, K. A.; Hosmillo, M.; Carter, A. P.; Ebrahimi, S.; Ke, Z.; Scheres, S. H. W.; Bergamaschi, L.; Grice, G. L., Zhang, Y.; CITIID-NIHR COVID-19 BioResource Collaboration, Nathan, J.A.; Baker, S.; James, L.C.; Baxendale, H.E.; Goodfellow, I.; Doffinger, R.; Briggs, J.A.G. A thermostable, closed SARS-CoV-2 spike protein trimer. Nat. Struct. Mol. Biol. 2020, 27, 934–941. doi: 10.1038/s41594-020-0478-5.

10. Costello, S.M.; Shoemaker, S.R.; Hobbs, H.T.; Nguyen, A.W.; Hsieh, C.L.; Maynard, J.A.; McLellan, J.S.; Pak, J.E.; Marqusee, S. The SARS-CoV-2 spike reversibly samples an open-trimer conformation exposing novel epitopes. Nat. Struct. Mol. Biol. 2022, 27, 229–238. doi: 10.1038/s41594-022-00735-5.

11. McCormick, K.D.; Jacobs, J.L.; Mellors, J.W. The emerging plasticity of SARS-CoV-2. Science 2021, 371, 1306–1308. doi: 10.1126/science.abg4493.

12. Ghimire, D.; Han, Y.; Lu, M. Structural Plasticity and Immune Evasion of SARS-CoV-2 Spike Variants. Viruses 2022, 14, 1255. 10.3390/v14061255.

13. Xu, C.; Wang, Y.; Liu, C.; Zhang, C.; Han, W.; Hong, X.; Wang, Y.; Hong, Q.; Wang, S.; Zhao, Q.; Wang, Y.; Yang, Y.; Chen, K.; Zheng, W.; Kong, L.; Wang, F.; Zuo, Q.; Huang, Z.; Cong, Y. Conformational dynamics of SARS-CoV-2 trimeric spike glycoprotein in complex with receptor ACE2 revealed by cryo-EM. Sci. Adv. 2021, 7, eabe5575. doi: 10.1126/sciadv.abe5575.

14. Benton, D. J.; Wrobel, A. G.; Xu, P.; Roustan, C.; Martin, S. R.; Rosenthal, P. B.; Skehel, J. J.; Gamblin, S. J. Receptor binding and priming of the spike protein of SARS-CoV-2 for membrane fusion. Nature 2020, 588, 327–330. doi: 10.1038/s41586-020-2772-0.

15. Turoňová, B.; Sikora, M.; Schuerman, C.; Hagen, W. J. H.; Welsch, S.; Blanc, F. E. C.; von Bülow, S.; Gecht, M.; Bagola, K.; Hörner, C.; van Zandbergen, G.; Landry, J.; de Azevedo, N. T. D.; Mosalaganti, S.; Schwarz, A.; Covino, R.; Mühlebach, M. D.; Hummer, G.; Krijnse Locker, J.; Beck, M. In situ structural analysis of SARS-CoV-2 spike reveals flexibility mediated by three hinges. Science 2020, 370, 203–208. doi: 10.1126/science.abd5223.

16. Lu, M.; Uchil, P. D.; Li, W.; Zheng, D.; Terry, D. S.; Gorman, J.; Shi, W.; Zhang, B.; Zhou, T.; Ding, S.; Gasser, R.; Prevost, J.; Beaudoin-Bussieres, G.; Anand, S. P.; Laumaea, A.; Grover, J. R.; Lihong, L.; Ho, D. D.; Mascola, J.R.; Finzi, A.; Kwong, P. D.; Blanchard, S. C.; Mothes, W. Real-time conformational dynamics of SARS-CoV-2 spikes on virus particles. Cell Host Microbe. 2020, 28, 880–891.e8. doi: 10.1016/j.chom.2020.11.001.

17. Yang, Z.; Han, Y.; Ding, S.; Shi, W.; Zhou, T.; Finzi, A.; Kwong, P.D.; Mothes, W.; Lu, M. SARS-CoV-2 Variants Increase Kinetic Stability of Open Spike Conformations as an Evolutionary Strategy. mBio 2022, 13, e0322721. doi: 10.1128/mbio.03227-21.

18. Díaz-Salinas, M.A.; Li, Q.; Ejemel, M.; Yurkovetskiy, L.; Luban, J.; Shen, K.; Wang, Y.; Munro, J.B. Conformational dynamics and allosteric modulation of the SARS-CoV-2 spike. Elife 2022, 11, e75433. doi: 10.7554/eLife.75433.

19. Wang, Y.; Liu, C.; Zhang, C.; Wang, Y.; Hong, Q.; Xu, S.; Li, Z.; Yang, Y.; Huang, Z.; Cong, Y. Structural Basis for SARS-CoV-2 Delta Variant Recognition of ACE2 Receptor and Broadly Neutralizing Antibodies. Nat. Commun. 2022, 13, 871. doi: 10.1038/s41467-022-28528-w.

20. Mannar, D.; Saville, J.W.; Zhu, X.; Srivastava, S.S.; Berezuk, A.M.; Tuttle, K.S.; Marquez, A.C.; Sekirov, I.; Subramaniam, S. SARS-CoV-2 Omicron Variant: Ab Evasion and Cryo-EM Structure of Spike Protein–ACE2 Complex. Science 2022, 375, 760–764. doi: 10.1126/science.abn7760.

21. Hong, Q.; Han, W.; Li, J.; Xu, S.; Wang, Y.; Xu, C.; Li, Z.; Wang, Y.; Zhang, C.; Huang, Z.; Cong, Y. Molecular Basis of Receptor Binding and Ab Neutralization of Omicron. Nature 2022. doi: 10.1038/s41586-022-04581-9.

22. McCallum, M.; Czudnochowski, N.; Rosen, L.E.; Zepeda, S.K.; Bowen, J.E.; Walls, A.C.; Hauser, K.; Joshi, A.; Stewart, C.; Dillen, J.R.; Powell, A.E.; Croll, T.I.; Nix, J.; Virgin, H.W.; Corti, D.; Snell, G.; Veesler, D. Structural Basis of SARS-CoV-2 Omicron Immune Evasion and Receptor Engagement. Science 2022, 375, 864–868. doi: 10.1126/science.abn8652.

23. Yin, W.; Xu, Y.; Xu, P.; Cao, X.; Wu, C.; Gu, C.; He, X.; Wang, X.; Huang, S.; Yuan, Q.; Wu, K.; Hu, W.; Huang, Z.; Liu, J.; Wang, Z.; Jia, F.; Xia, K.; Liu, P.; Wang, X.; Song, B.; Zheng, J.; Jiang, H.; Cheng, X.; Jiang, Y.; Deng, S.J.; Xu, H.E. Structures of the Omicron Spike Trimer with ACE2 and an Anti-Omicron Ab. Science 2022, 375, 1048–1053. doi: 10.1126/science.abn8863.

24. Gobeil, S. M.-C.; Henderson, R.; Stalls, V.; Janowska, K.; Huang, X.; May, A.; Speakman, M.; Beaudoin, E.; Manne, K.; Li, D.; Parks, R.; Barr, M.; Deyton, M.; Martin, M.; Mansouri, K.; Edwards, R. J.; Eaton, A.; Montefiori, D. C.; Sempowski, G. D.; Saunders, K. O.; Wiehe, K.; Williams, W.; Korber, B.; Haynes, B. F.; Acharya, P. Structural Diversity of the SARS-CoV-2 Omicron Spike. Mol Cell. 2022, 82, 2050–2068.e6. doi: 10.1016/j.molcel.2022.03.028.

25. Cui, Z.; Liu, P.; Wang, N.; Wang, L.; Fan, K.; Zhu, Q.; Wang, K.; Chen, R.; Feng, R.; Jia, Z.; Yang, M.; Xu, G.; Zhu, B.; Fu, W.; Chu, T.; Feng, L.; Wang, Y.; Pei, X.; Yang, P.; Xie, X.S.; Cao, L.; Cao, Y.; Wang, X. Structural and Functional Characterizations of Infectivity and Immune Evasion of SARS-CoV-2 Omicron. Cell 2022, 185, 860–871.e13. doi: 10.1016/j.cell.2022.01.019.

26. Parums DV. Editorial: The XBB.1.5 (’Kraken’) Subvariant of Omicron SARS-CoV-2 and its Rapid Global Spread. Med Sci Monit. 2023, 29, e939580. doi: 10.12659/MSM.939580.

27. Wang, Q.; Iketani, S.; Li, Z.; Liu, L.; Guo, Y.; Huang, Y.; Bowen, A. D.; Liu, M.; Wang, M.; Yu, J.; Valdez, R.; Lauring, A. S.; Sheng, Z.; Wang, H. H.; Gordon, A.; Liu, L.; Ho, D. D. Alarming Ab Evasion Properties of Rising SARS-CoV-2 BQ and XBB Subvariants. Cell 2023, 186, 279–286.e8. 10.1016/j.cell.2022.12.018.

28. Hoffmann, M.; Arora, P.; Nehlmeier, I.; Kempf, A.; Cossmann, A.; Schulz, S. R.; Morillas Ramos, G.; Manthey, L. A.; Jäck, H.-M.; Behrens, G. M. N.; Pöhlmann, S. Profound Neutralization Evasion and Augmented Host Cell Entry Are Hallmarks of the Fast-Spreading SARS-CoV-2 Lineage XBB.1.5. Cell Mol Immunol. 2023, 1–4. doi: 10.1038/s41423-023-00988-0.

29. Yamasoba, D.; Uriu, K.; Plianchaisuk, A.; Kosugi, Y.; Pan, L.; Zahradnik, J.; Ito, J.; Sato, K. Virological Characteristics of the SARS-CoV-2 Omicron XBB.1.16 Variant. Lancet Infect Dis. 2023, S1473-3099(23)00278-5. doi: 10.1016/S1473-3099(23)00278-5.

30. Tsujino, S.; Deguchi, S.; Nomai, T.; Padilla Blanco, M.; Plianchaisuk, A.; Wang, L.; Begum, M. M.; Uriu, K.; Mizuma, K.; Nao, N.; Kojima, I.; Tsubo, T.; Li, J.; Matsumura, Y.; Nagao, M.; Oda, Y.; Tsuda, M.; Anraku, Y.; Kita, S.; Yajima, H.; Sasaki Tabata, K.; Guo, Z.; Hinay, A. A., Jr.; Yoshimatsu, K.; Yamamoto, Y.; Nagamoto, T.; Asakura, H.; Nagashima, M.; Sadamasu, K.; Yoshimura, K.; Nasser, H.; Jonathan, M.; Putri, O.; Kim, Y.; Chen, L.; Suzuki, R.; Tamura, T.; Maenaka, K.; Irie, T.; Matsuno, K.; Tanaka, S.; Ito, J.; Ikeda, T.; Takayama, K.; Zahradnik, J.; Hashiguchi, T.; Fukuhara, T.; Sato, K. Virological Characteristics of the SARS CoV 2 Omicron EG.5.1 Variant. Microbiol Immunol. 2024, 68, 305–330. doi: 10.1111/1348-0421.13165

31. Wang, Q.; Guo, Y.; Zhang, R. M.; Ho, J.; Mohri, H.; Valdez, R.; Manthei, D. M.; Gordon, A.; Liu, L.; Ho, D. D. Ab Neutralization of Emerging SARS-CoV-2 Subvariants: EG.5.1 and XBC.1.6. Lancet Infect Dis. 2023, 23, e397–e398. doi: 10.1016/S1473-3099(23)00555-8.

32. Faraone, J. N.; Qu, P.; Goodarzi, N.; Zheng, Y.-M.; Carlin, C.; Saif, L. J.; Oltz, E. M.; Xu, K.; Jones, D.; Gumina, R. J.; Liu, S.-L. Immune Evasion and Membrane Fusion of SARS-CoV-2 XBB Subvariants EG.5.1 and XBB.2.3. Emerg. Microbes Infect. 2023, 12, 2270069. doi: 10.1080/22221751.2023.2270069.

33. Kosugi, Y.; Plianchaisuk, A.; Putri, O.; Uriu, K.; Kaku, Y.; Hinay, A. A., Jr; Chen, L.; Kuramochi, J.; Sadamasu, K.; Yoshimura, K.; Asakura, H.; Nagashima, M.; Ito, J.; Sato, K.; Misawa, N.; Guo, Z.; Tolentino, J. E. M.; Fujita, S.; Pan, L.; Suganami, M.; Chiba, M.; Yoshimura, R.; Yasuda, K.; Iida, K.; Ohsumi, N.; Strange, A. P.; Tanaka, S.; Fukuhara, T.; Tamura, T.; Suzuki, R.; Suzuki, S.; Ito, H.; Matsuno, K.; Sawa, H.; Nao, N.; Tanaka, S.; Tsuda, M.; Wang, L.; Oda, Y.; Ferdous, Z.; Shishido, K.; Nakagawa, S.; Shirakawa, K.; Takaori-Kondo, A.; Nagata, K.; Nomura, R.; Horisawa, Y.; Tashiro, Y.; Kawai, Y.; Takayama, K.; Hashimoto, R.; Deguchi, S.; Watanabe, Y.; Sakamoto, A.; Yasuhara, N.; Hashiguchi, T.; Suzuki, T.; Kimura, K.; Sasaki, J.; Nakajima, Y.; Yajima, H.; Irie, T.; Kawabata, R.; Tabata, K.; Ikeda, T.; Nasser, H.; Shimizu, R.; Begum, M. M.; Jonathan, M.; Mugita, Y.; Takahashi, O.; Ichihara, K.; Ueno, T.; Motozono, C.; Toyoda, M.; Saito, A.; Shofa, M.; Shibatani, Y.; Nishiuchi, T. Characteristics of the SARS-CoV-2 Omicron HK.3 Variant Harbouring the FLip Substitution. Lancet Microbe 2024, S2666-5247(23)00373-7. doi: 10.1016/S2666-5247(23)00373-7.

34. Wang, Q.; Guo, Y.; Liu, L.; Schwanz, L. T.; Li, Z.; Nair, M. S.; Ho, J.; Zhang, R. M.; Iketani, S.; Yu, J.; Huang, Y.; Qu, Y.; Valdez, R.; Lauring, A. S.; Huang, Y.; Gordon, A.; Wang, H. H.; Liu, L.; Ho, D. D. Antigenicity and Receptor Affinity of SARS-CoV-2 BA.2.86 Spike. Nature 2023. 10.1038/s41586-023-06750-w.

35. Yang, S.; Yu, Y.; Jian, F.; Song, W.; Yisimayi, A.; Chen, X.; Xu, Y.; Wang, P.; Wang, J.; Yu, L.; Niu, X.; Wang, J.; Xiao, T.; An, R.; Wang, Y.; Gu, Q.; Shao, F.; Jin, R.; Shen, Z.; Wang, Y.; Cao, Y. Antigenicity and Infectivity Characterization of SARS-CoV-2 BA.2.86. Lancet Infect Dis. 2023, 23, e457–e459. doi: 10.1016/S1473-3099(23)00573-X.

36. Tamura, T.; Mizuma, K.; Nasser, H.; Deguchi, S.; Padilla-Blanco, M.; Oda, Y.; Uriu, K.; Tolentino, J. E. M.; Tsujino, S.; Suzuki, R.; Kojima, I.; Nao, N.; Shimizu, R.; Wang, L.; Tsuda, M.; Jonathan, M.; Kosugi, Y.; Guo, Z.; Hinay, A. A., Jr.; Putri, O.; Kim, Y.; Tanaka, Y. L.; Asakura, H.; Nagashima, M.; Sadamasu, K.; Yoshimura, K.; Saito, A.; Ito, J.; Irie, T.; Tanaka, S.; Zahradnik, J.; Ikeda, T.; Takayama, K.; Matsuno, K.; Fukuhara, T.; Sato, K. Virological Characteristics of the SARS-CoV-2 BA.2.86 Variant. Cell Host Microbe. 2024, 32, 170–180.e12. doi: 10.1016/j.chom.2024.01.001.

37. Liu, C.; Zhou, D.; Dijokaite-Guraliuc, A.; Supasa, P.; Duyvesteyn, H. M. E.; Ginn, H. M.; Selvaraj, M.; Mentzer, A. J.; Das, R.; de Silva, T. I.; Ritter, T. G.; Plowright, M.; Newman, T. A. H.; Stafford, L.; Kronsteiner, B.; Temperton, N.; Lui, Y.; Fellermeyer, M.; Goulder, P.; Klenerman, P.; Dunachie, S. J.; Barton, M. I.; Kutuzov, M. A.; Dushek, O.; Fry, E. E.; Mongkolsapaya, J.; Ren, J.; Stuart, D. I.; Screaton, G. R. A Structure-Function Analysis SARS-CoV-2 BA.2.86 Balances Ab Escape and ACE2 Affinity. Cell Rep Med. 2024, 5, 101553. doi: 10.1016/j.xcrm.2024.101553.

38. Khan, K.; Lustig, G.; Römer, C.; Reedoy, K.; Jule, Z.; Karim, F.; Ganga, Y.; Bernstein, M.; Baig, Z.; Jackson, L.; Mahlangu, B.; Mnguni, A.; Nzimande, A.; Stock, N.; Kekana, D.; Ntozini, B.; van Deventer, C.; Marshall, T.; Manickchund, N.; Gosnell, B. I.; Lessells, R. J.; Karim, Q. A.; Abdool Karim, S. S.; Moosa, M.-Y. S.; de Oliveira, T.; von Gottberg, A.; Wolter, N.; Neher, R. A.; Sigal, A. Evolution and Neutralization Escape of the SARS-CoV-2 BA.2.86 Subvariant. Nat Commun. 2023, 14, 8078. doi: 10.1038/s41467-023-43703-3.

39. Yang, S.; Yu, Y.; Xu, Y.; Jian, F.; Song, W.; Yisimayi, A.; Wang, P.; Wang, J.; Liu, J.; Yu, L.; Niu, X.; Wang, J.; Wang, Y.; Shao, F.; Jin, R.; Wang, Y.; Cao, Y. Fast Evolution of SARS-CoV-2 BA.2.86 to JN.1 under Heavy Immune Pressure. Lancet Infect Dis. 2024, 24, e70–e72. doi: 10.1016/S1473-3099(23)00744-2.

40. Kaku, Y.; Okumura, K.; Padilla-Blanco, M.; Kosugi, Y.; Uriu, K.; Hinay, A. A., Jr; Chen, L.; Plianchaisuk, A.; Kobiyama, K.; Ishii, K. J.; Zahradnik, J.; Ito, J.; Sato, K., K. Virological Characteristics of the SARS-CoV-2 JN.1 Variant. Lancet Infect Dis. 2024, 24, e82. doi: 10.1016/S1473-3099(23)00813-7.

41. Li, P.; Faraone, J. N.; Hsu, C. C.; Chamblee, M.; Zheng, Y.-M.; Carlin, C.; Bednash, J. S.; Horowitz, J. C.; Mallampalli, R. K.; Saif, L. J.; Oltz, E. M.; Jones, D.; Li, J.; Gumina, R. J.; Xu, K.; Liu, S.-L. Neutralization Escape, Infectivity, and Membrane Fusion of JN.1-Derived SARS-CoV-2 SLip, FLiRT, and KP.2 Variants. Cell Rep. 2024, 43, 114520. doi: 10.1016/j.celrep.2024.114520.

42. Qu, P.; Faraone, J. N.; Evans, J. P.; Zheng, Y.-M.; Carlin, C.; Anghelina, M.; Stevens, P.; Fernandez, S.; Jones, D.; Panchal, A. R.; Saif, L. J.; Oltz, E. M.; Zhang, B.; Zhou, T.; Xu, K.; Gumina, R. J.; Liu, S.-L. Enhanced Evasion of Neutralizing Ab Response by Omicron XBB.1.5, CH.1.1, and CA.3.1 Variants. Cell Rep. 2023, 42, 112443. doi: 10.1016/j.celrep.2023.112443.

43. Kaku, Y.; Uriu, K.; Kosugi, Y.; Okumura, K.; Yamasoba, D.; Uwamino, Y.; Kuramochi, J.; Sadamasu, K.; Yoshimura, K.; Asakura, H.; Nagashima, M.; Ito, J.; Sato, K. Virological Characteristics of the SARS-CoV-2 KP.2 Variant. Lancet Infect Dis. 2024, 24, e416. doi: 10.1016/S1473-3099(24)00298-6.

44. Kaku, Y.; Yo, M. S.; Tolentino, J. E.; Uriu, K.; Okumura, K.; Ito, J.; Sato, K. Virological Characteristics of the SARS-CoV-2 KP.3, LB.1, and KP.2.3 Variants. Lancet Infect Dis. 2024, 24, e482–e483. doi: 10.1016/S1473-3099(24)00415-8.

45. Wang, Q.; Mellis, I. A.; Ho, J.; Bowen, A.; Kowalski-Dobson, T.; Valdez, R.; Katsamba, P. S.; Wu, M.; Lee, C.; Shapiro, L.; Gordon, A.; Guo, Y.; Ho, D. D.; Liu, L. Recurrent SARS-CoV-2 Spike Mutations Confer Growth Advantages to Select JN.1 Sublineages. Emerg Microbes Infect. 2024, 13, 2402880. doi: 10.1080/22221751.2024.2402880.

46. Yang, J.; He, X.; Shi, H.; He, C.; Lei, H.; He, H.; Yang, L.; Wang, W.; Shen, G.; Yang, J.; Zhao, Z.; Song, X.; Wang, Z.; Lu, G.; Li, J.; Wei, Y. Recombinant XBB.1.5 Boosters Induce Robust Neutralization against KP.2- and KP.3-Included JN.1 Sublineages. Sig Transduct Target Ther 2025, 10, 47. doi: 10.1038/s41392-025-02139-5.

47. Taylor, A. L.; Starr, T. N. Deep Mutational Scanning of SARS-CoV-2 Omicron BA.2.86 and Epistatic Emergence of the KP.3 Variant. Virus Evol. 2024, 10, veae067. doi: 10.1093/ve/veae067.

48. Feng, L.; Sun, Z.; Zhang, Y.; Jian, F.; Yang, S.; Xia, K.; Yu, L.; Wang, J.; Shao, F.; Wang, X.; Cao, Y. Structural and Molecular Basis of the Epistasis Effect in Enhanced Affinity between SARS-CoV-2 KP.3 and ACE2. Cell Discov. 2024, 10, 123. doi: 10.1038/s41421-024-00752-2.

49. Liu, J.; Yu, Y.; Jian, F.; Yang, S.; Song, W.; Wang, P.; Yu, L.; Shao, F.; Cao, Y. Enhanced Immune Evasion of SARS-CoV-2 Variants KP.3.1.1 and XEC through N-Terminal Domain Mutations. Lancet Infect Dis. 2025, 25, e6–e7. doi: 10.1016/S1473-3099(24)00738-2.

50. Kaku, Y.; Uriu, K.; Okumura, K.; Ito, J.; Sato, K. Virological Characteristics of the SARS-CoV-2 KP.3.1.1 Variant. Lancet Infect Dis. 2024, 24, e609. doi: 10.1016/S1473-3099(24)00505-X.

51. Kaku, Y.; Okumura, K.; Kawakubo, S.; Uriu, K.; Chen, L.; Kosugi, Y.; Uwamino, Y.; Begum, M. M.; Leong, S.; Ikeda, T.; Sadamasu, K.; Asakura, H.; Nagashima, M.; Yoshimura, K.; Ito, J.; Sato, K. Virological Characteristics of the SARS-CoV-2 XEC Variant. Lancet Infect Dis. 2024, S1473-3099(24)00731-X. doi: 10.1016/S1473-3099(24)00731-X.

52. Wang, Q.; Guo, Y.; Mellis, I. A.; Wu, M.; Mohri, H.; Gherasim, C.; Valdez, R.; Purpura, L. J.; Yin, M. T.; Gordon, A.; Ho, D. D. Antibody Evasiveness of SARS-CoV-2 Subvariants KP.3.1.1 and XEC. Cell Rep. 2025, 44,115543. doi: 10.1016/j.celrep.2025.115543.

53. Feng, Z.; Huang, J.; Baboo, S.; Diedrich, J. K.; Bangaru, S.; Paulson, J. C.; Yates, J. R., III; Yuan, M.; Wilson, I. A.; Ward, A. B. Structural and Functional Insights into the Evolution of SARS-CoV-2 KP.3.1.1 Spike Protein, bioRxiv 2024. doi: 10.1101/2024.12.10.627775.

54. Kaku, Y.; Uriu, K.; Okumura, K.; Ito, J.; Sato, K. Virological Characteristics of the SARS-CoV-2 KP.3.1.1 Variant. Lancet Infect Dis. 2024, 24, e609. doi: 10.1016/S1473-3099(24)00505-X.

55. Liu, J.; Yu, Y.; Yang, S.; Jian, F.; Song, W.; Yu, L.; Shao, F.; Cao, Y. Virological and Antigenic Characteristics of SARS-CoV-2 Variants LF.7.2.1, NP.1, and LP.8.1. Lancet Infect Dis. 2025, 25, e128–e130. doi: 10.1016/S1473-3099(25)00015-5.

56. Guo, C.; Yu, Y.; Liu, J.; Jian, F.; Yang, S.; Song, W.; Yu, L.; Shao, F.; Cao, Y. Antigenic and Virological Characteristics of SARS-CoV-2 Variants BA.3.2, XFG, and NB.1.8.1. Lancet Infect Dis. 2025. 10.1016/s1473-3099(25)00308-1.

57. Barnes, C. O.; Jette, C. A.; Abernathy, M. E.; Dam, K.-M. A.; Esswein, S. R.; Gristick, H. B.; Malyutin, A. G.; Sharaf, N. G.; Huey-Tubman, K. E.; Lee, Y. E.; Robbiani, D. F.; Nussenzweig, M. C.; West, A. P., Jr; Bjorkman, P. J. SARS-CoV-2 Neutralizing Antibody Structures Inform Therapeutic Strategies. Nature 2020, 588, 682–687. doi: 10.1038/s41586-020-2852-1.

58. Chen, Y., Zhao, X., Zhou, H. et al. Broadly neutralizing antibodies to SARS-CoV-2 and other human coronaviruses. Nat Rev Immunol 23, 189–199 (2023). 10.1038/s41577-022-00784-3

59. Pinto, D.; Park, Y.-J.; Beltramello, M.; Walls, A. C.; Tortorici, M. A.; Bianchi, S.; Jaconi, S.; Culap, K.; Zatta, F.; De Marco, A.; Peter, A.; Guarino, B.; Spreafico, R.; Cameroni, E.; Case, J. B.; Chen, R. E.; Havenar-Daughton, C.; Snell, G.; Telenti, A.; Virgin, H. W.; Lanzavecchia, A.; Diamond, M. S.; Fink, K.; Veesler, D.; Corti, D. Cross-Neutralization of SARS-CoV-2 by a Human Monoclonal SARS-CoV Antibody. Nature 2020, 583, 290–295. doi: 10.1038/s41586-020-2349-y.

60. Tortorici, M. A.; Beltramello, M.; Lempp, F. A.; Pinto, D.; Dang, H. V.; Rosen, L. E.; McCallum, M.; Bowen, J.; Minola, A.; Jaconi, S.; Zatta, F.; De Marco, A.; Guarino, B.; Bianchi, S.; Lauron, E. J.; Tucker, H.; Zhou, J.; Peter, A.; Havenar-Daughton, C.; Wojcechowskyj, J. A.; Case, J. B.; Chen, R. E.; Kaiser, H.; Montiel-Ruiz, M.; Meury, M.; Czudnochowski, N.; Spreafico, R.; Dillen, J.; Ng, C.; Sprugasci, N.; Culap, K.; Benigni, F.; Abdelnabi, R.; Foo, S.-Y. C.; Schmid, M. A.; Cameroni, E.; Riva, A.; Gabrieli, A.; Galli, M.; Pizzuto, M. S.; Neyts, J.; Diamond, M. S.; Virgin, H. W.; Snell, G.; Corti, D.; Fink, K.; Veesler, D. Ultrapotent Human Antibodies Protect against SARS-CoV-2 Challenge via Multiple Mechanisms. Science 2020, 370, 950–957. doi: 10.1126/science.abe3354. Yuan, M.; Wu, N. C.; Zhu, X.; Lee, C.-C. D.; So, R. T. Y.; Lv, H.; Mok, C. K. P.; Wilson, I. A. A Highly Conserved Cryptic Epitope in the Receptor Binding Domains of SARS-CoV-2 and SARS-CoV. Science. 2020, 368, 630–633. doi: 10.1126/science.abb7269.

61. Cao, Y.; Wang, J.; Jian, F.; Xiao, T.; Song, W.; Yisimayi, A.; Huang, W.; Li, Q.; Wang, P.; An, R.; Wang, J.; Wang, Y.; Niu, X.; Yang, S.; Liang, H.; Sun, H.; Li, T.; Yu, Y.; Cui, Q.; Liu, S.; Yang, X.; Du, S.; Zhang, Z.; Hao, X.; Shao, F.; Jin, R.; Wang, X.; Xiao, J.; Wang, Y.; Xie, X. S. Omicron Escapes the Majority of Existing SARS-CoV-2 Neutralizing Antibodies. Nature 2022, 602, 657–663. doi: 10.1038/s41586-021-04385-3. Cao, Y.; Yisimayi, A.; Jian, F.; Xiao, T.; Song, W.; Wang, J.; Du, S.; Zhang, Z.; Liu, P.; Hao, X.; Li, Q.; Chen, X.; Wang, L.; Wang, P.; An, R.; Wang, Y.; Wang, J.; Yang, P.; Sun, H.; Zhao, L.; Zhang, W.; Zhao, D.; Zheng, J.; Yu, L.; Li, C.; Zhang, N.; Wang, R.; Niu, X.; Yang, S.; Song, X.; Zheng, L.; Li, Z.; Gu, Q.; Shao, F.; Huang, W.; Wang, Y.; Shen, Z.; Wang, X.; Jin, R.; Xiao, J.; Xie, X. S. Omicron BA.2 Specifically Evades Broad Sarbecovirus Neutralizing Antibodies, bioRxiv 2022. 10.1101/2022.02.07.479349.

62. Rosen, L. E.; Tortorici, M. A.; De Marco, A.; Pinto, D.; Foreman, W. B.; Taylor, A. L.; Park, Y.-J.; Bohan, D.; Rietz, T.; Errico, J. M.; Hauser, K.; Dang, H. V.; Chartron, J. W.; Giurdanella, M.; Cusumano, G.; Saliba, C.; Zatta, F.; Sprouse, K. R.; Addetia, A.; Zepeda, S. K.; Brown, J.; Lee, J.; Dellota, E., Jr.; Rajesh, A.; Noack, J.; Tao, Q.; DaCosta, Y.; Tsu, B.; Acosta, R.; Subramanian, S.; de Melo, G. D.; Kergoat, L.; Zhang, I.; Liu, Z.; Guarino, B.; Schmid, M. A.; Schnell, G.; Miller, J. L.; Lempp, F. A.; Czudnochowski, N.; Cameroni, E.; Whelan, S. P. J.; Bourhy, H.; Purcell, L. A.; Benigni, F.; di Iulio, J.; Pizzuto, M. S.; Lanzavecchia, A.; Telenti, A.; Snell, G.; Corti, D.; Veesler, D.; Starr, T. N. A Potent Pan-Sarbecovirus Neutralizing Antibody Resilient to Epitope Diversification. Cell 2024, 187, 7196–7213.e26. 10.1016/j.cell.2024.09.026.

63. Piccoli, L.; Park, Y.-J.; Tortorici, M. A.; Czudnochowski, N.; Walls, A. C.; Beltramello, M.; Silacci-Fregni, C.; Pinto, D.; Rosen, L. E.; Bowen, J. E.; Acton, O. J.; Jaconi, S.; Guarino, B.; Minola, A.; Zatta, F.; Sprugasci, N.; Bassi, J.; Peter, A.; De Marco, A.; Nix, J. C.; Mele, F.; Jovic, S.; Rodriguez, B. F.; Gupta, S. V.; Jin, F.; Piumatti, G.; Lo Presti, G.; Pellanda, A. F.; Biggiogero, M.; Tarkowski, M.; Pizzuto, M. S.; Cameroni, E.; Havenar-Daughton, C.; Smithey, M.; Hong, D.; Lepori, V.; Albanese, E.; Ceschi, A.; Bernasconi, E.; Elzi, L.; Ferrari, P.; Garzoni, C.; Riva, A.; Snell, G.; Sallusto, F.; Fink, K.; Virgin, H. W.; Lanzavecchia, A.; Corti, D.; Veesler, D. Mapping Neutralizing and Immunodominant Sites on the SARS-CoV-2 Spike Receptor-Binding Domain by Structure-Guided High-Resolution Serology. Cell 2020, 183, 1024–1042.e21. doi: 10.1016/j.cell.2020.09.037.

64. Martinez, D. R.; Schäfer, A.; Gobeil, S.; Li, D.; De la Cruz, G.; Parks, R.; Lu, X.; Barr, M.; Stalls, V.; Janowska, K.; Beaudoin, E.; Manne, K.; Mansouri, K.; Edwards, R. J.; Cronin, K.; Yount, B.; Anasti, K.; Montgomery, S. A.; Tang, J.; Golding, H.; Shen, S.; Zhou, T.; Kwong, P. D.; Graham, B. S.; Mascola, J. R.; Montefiori, D. C.; Alam, S. M.; Sempowski, G. D.; Khurana, S.; Wiehe, K.; Saunders, K. O.; Acharya, P.; Haynes, B. F.; Baric, R. S. A Broadly Cross-Reactive Antibody Neutralizes and Protects against Sarbecovirus Challenge in Mice. Sci Transl Med. 2022, 14, eabj7125. doi: 10.1126/scitranslmed.abj7125.

65. Rappazzo, C. G.; Tse, L. V.; Kaku, C. I.; Wrapp, D.; Sakharkar, M.; Huang, D.; Deveau, L. M.; Yockachonis, T. J.; Herbert, A. S.; Battles, M. B.; O’Brien, C. M.; Brown, M. E.; Geoghegan, J. C.; Belk, J.; Peng, L.; Yang, L.; Hou, Y.; Scobey, T. D.; Burton, D. R.; Nemazee, D.; Dye, J. M.; Voss, J. E.; Gunn, B. M.; McLellan, J. S.; Baric, R. S.; Gralinski, L. E.; Walker, L. M. Broad and Potent Activity against SARS-like Viruses by an Engineered Human Monoclonal Antibody. Science. 2021, 371, 823–829. doi: 10.1126/science.abf4830.

66. Yuan, M.; Zhu, X.; He, W.; Zhou, P.; Kaku, C. I.; Capozzola, T.; Zhu, C. Y.; Yu, X.; Liu, H.; Yu, W.; Hua, Y.; Tien, H.; Peng, L.; Song, G.; Cottrell, C. A.; Schief, W. R.; Nemazee, D.; Walker, L. M.; Andrabi, R.; Burton, D. R.; Wilson, I. A. A Broad and Potent Neutralization Epitope in SARS-Related Coronaviruses. Proc. Natl. Acad. Sci. U.S.A. 2022, 119, e2205784119. doi: 10.1073/pnas.2205784119.

67. Cao, Y.; Yisimayi, A.; Jian, F.; Song, W.; Xiao, T.; Wang, L.; Du, S.; Wang, J.; Li, Q.; Chen, X.; Yu, Y.; Wang, P.; Zhang, Z.; Liu, P.; An, R.; Hao, X.; Wang, Y.; Wang, J.; Feng, R.; Sun, H.; Zhao, L.; Zhang, W.; Zhao, D.; Zheng, J.; Yu, L.; Li, C.; Zhang, N.; Wang, R.; Niu, X.; Yang, S.; Song, X.; Chai, Y.; Hu, Y.; Shi, Y.; Zheng, L.; Li, Z.; Gu, Q.; Shao, F.; Huang, W.; Jin, R.; Shen, Z.; Wang, Y.; Wang, X.; Xiao, J.; Xie, X. S. BA.2.12.1, BA.4 and BA.5 Escape Antibodies Elicited by Omicron Infection. Nature 2022, 608, 593-602. doi: 10.1038/s41586-022-04980-y.

68. Cao, Y.; Jian, F.; Wang, J.; Yu, Y.; Song, W.; Yisimayi, A.; Wang, J.; An, R.; Chen, X.; Zhang, N.; Wang, Y.; Wang, P.; Zhao, L.; Sun, H.; Yu, L.; Yang, S.; Niu, X.; Xiao, T.; Gu, Q.; Shao, F.; Hao, X.; Xu, Y.; Jin, R.; Shen, Z.; Wang, Y.; Xie, X. S. Imprinted SARS-CoV-2 Humoral Immunity Induces Convergent Omicron RBD Evolution. Nature 2023, 614, 521–529. doi: 10.1038/s41586-022-05644-7.

69. Jian, F.; Wang, J.; Yisimayi, A.; Song, W.; Xu, Y.; Chen, X.; Niu, X.; Yang, S.; Yu, Y.; Wang, P.; Sun, H.; Yu, L.; Wang, J.; Wang, Y.; An, R.; Wang, W.; Ma, M.; Xiao, T.; Gu, Q.; Shao, F.; Wang, Y.; Shen, Z.; Jin, R.; Cao, Y. Evolving Antibody Response to SARS-CoV-2 Antigenic Shift from XBB to JN.1. Nature 2025, 637, 921–929. doi: 10.1038/s41586-024-08315-x.

70. Cao, Y.; Jian, F.; Zhang, Z.; Yisimayi, A.; Hao, X.; Bao, L.; Yuan, F.; Yu, Y.; Du, S.; Wang, J.; Xiao, T.; Song, W.; Zhang, Y.; Liu, P.; An, R.; Wang, P.; Wang, Y.; Yang, S.; Niu, X.; Zhang, Y.; Gu, Q.; Shao, F.; Hu, Y.; Yin, W.; Zheng, A.; Wang, Y.; Qin, C.; Jin, R.; Xiao, J.; Xie, X. S. Rational Identification of Potent and Broad Sarbecovirus-Neutralizing Antibody Cocktails from SARS Convalescents. Cell Rep. 2022, 41, 111845. doi: 10.1016/j.celrep.2022.111845.

71. Yisimayi, A.; Song, W.; Wang, J.; Jian, F.; Yu, Y.; Chen, X.; Xu, Y.; Yang, S.; Niu, X.; Xiao, T.; Wang, J.; Zhao, L.; Sun, H.; An, R.; Zhang, N.; Wang, Y.; Wang, P.; Yu, L.; Lv, Z.; Gu, Q.; Shao, F.; Jin, R.; Shen, Z.; Xie, X. S.; Wang, Y.; Cao, Y. Repeated Omicron Exposures Override Ancestral SARS-CoV-2 Immune Imprinting. Nature 2024, 625, 148–156. doi: 10.1038/s41586-023-06753-7.

72. Jian, F.; Wec, A. Z.; Feng, L.; Yu, Y.; Wang, L.; Wang, P.; Yu, L.; Wang, J.; Hou, J.; Berrueta, D. M.; Lee, D.; Speidel, T.; Ma, L.; Kim, T.; Yisimayi, A.; Song, W.; Wang, J.; Liu, L.; Yang, S.; Niu, X.; Xiao, T.; An, R.; Wang, Y.; Shao, F.; Wang, Y.; Pecetta, S.; Wang, X.; Walker, L. M.; Cao, Y. Viral Evolution Prediction Identifies Broadly Neutralizing Antibodies to Existing and Prospective SARS-CoV-2 Variants. Nat Microbiol 2025. doi: 10.1038/s41564-025-02030-7.

73. Starr, T. N.; Czudnochowski, N.; Liu, Z.; Zatta, F.; Park, Y.-J.; Addetia, A.; Pinto, D.; Beltramello, M.; Hernandez, P.; Greaney, A. J.; Marzi, R.; Glass, W. G.; Zhang, I.; Dingens, A. S.; Bowen, J. E.; Tortorici, M. A.; Walls, A. C.; Wojcechowskyj, J. A.; De Marco, A.; Rosen, L. E.; Zhou, J.; Montiel-Ruiz, M.; Kaiser, H.; Dillen, J. R.; Tucker, H.; Bassi, J.; Silacci-Fregni, C.; Housley, M. P.; di Iulio, J.; Lombardo, G.; Agostini, M.; Sprugasci, N.; Culap, K.; Jaconi, S.; Meury, M.; Dellota Jr, E.; Abdelnabi, R.; Foo, S.-Y. C.; Cameroni, E.; Stumpf, S.; Croll, T. I.; Nix, J. C.; Havenar-Daughton, C.; Piccoli, L.; Benigni, F.; Neyts, J.; Telenti, A.; Lempp, F. A.; Pizzuto, M. S.; Chodera, J. D.; Hebner, C. M.; Virgin, H. W.; Whelan, S. P. J.; Veesler, D.; Corti, D.; Bloom, J. D.; Snell, G. SARS-CoV-2 RBD Antibodies That Maximize Breadth and Resistance to Escape. Nature 2021, 597, 97–102. doi: 10.1038/s41586-021-03807-6.

74. Jensen, J. L., Sankhala, R. S., Dussupt, V., Bai, H., Hajduczki, A., Lal, K. G., Chang, W. C., Martinez, E. J., Peterson, C. E., Golub, E. S., Rees, P. A., Mendez-Rivera, L., Zemil, M., Kavusak, E., Mayer, S. V., Wieczorek, L., Kannan, S., Doranz, B. J., Davidson, E., … Joyce, M. G. (2023). Targeting the Spike Receptor Binding Domain Class V Cryptic Epitope by an Antibody with Pan-Sarbecovirus Activity. J Virol. 2023, 97, e0159622. doi: 10.1128/jvi.01596-22.

75. Sankhala, R. S.; Dussupt, V.; Chen, W.-H.; Bai, H.; Martinez, E. J.; Jensen, J. L.; Rees, P. A. ; Hajduczki, A.; Chang, W. C.; Choe, M.; Yan, L.; Sterling, S. L.; Swafford, I.; Kuklis, C.; Soman, S.; King, J.; Corbitt, C.; Zemil, M.; Peterson, C. E.; Mendez-Rivera, L.; Townsley, S. M.; Donofrio, G. C.; Lal, K. G.; Tran, U.; Green, E. C.; Smith, C.; de Val, N.; Laing, E. D.; Broder, C. C.; Currier, J. R.; Gromowski, G. D.; Wieczorek, L.; Rolland, M.; Paquin-Proulx, D.; van Dyk, D.; Britton, Z.; Rajan, S.; Loo, Y. M.; McTamney, P. M.; Esser, M. T.; Polonis, V. R.; Michael, N. L.; Krebs, S. J.; Modjarrad, K.; Joyce, M. G. Antibody Targeting of Conserved Sites of Vulnerability on the SARS-CoV-2 Spike Receptor-Binding Domain. Structure 2024, 32, 131–147.e7. doi: 10.1016/j.str.2023.11.015.

76. Xiao, S.; Alshahrani, M.; Gupta, G.; Tao, P.; Verkhivker, G. Markov State Models and Perturbation-Based Approaches Reveal Distinct Dynamic Signatures and Hidden Allosteric Pockets in the Emerging SARS-Cov-2 Spike Omicron Variant Complexes with the Host Receptor: The Interplay of Dynamics and Convergent Evolution Modulates Allostery and Functional Mechanisms. J. Chem. Inf. Model. 2023, 63, 5272–5296. doi: 10.1021/acs.jcim.3c00778

77. Raisinghani, N.; Alshahrani, M.; Gupta, G.; Xiao, S.; Tao, P.; Verkhivker, G. AlphaFold2 Predictions of Conformational Ensembles and Atomistic Simulations of the SARS-CoV-2 Spike XBB Lineages Reveal Epistatic Couplings between Convergent Mutational Hotspots That Control ACE2 Affinity. J. Phys. Chem. B. 2024, 128, 4696–4715. doi: 10.1021/acs.jpcb.4c01341.

78. Raisinghani, N.; Alshahrani, M.; Gupta, G.; Verkhivker, G. Ensemble-Based Mutational Profiling and Network Analysis of the SARS-CoV-2 Spike Omicron XBB Lineages for Interactions with the ACE2 Receptor and Antibodies: Cooperation of Binding Hotspots in Mediating Epistatic Couplings Underlies Binding Mechanism and Immune Escape. Int. J. Mol. Sci. 2024, 25, 4281. doi: 10.3390/ijms25084281.

79. Raisinghani, N.; Alshahrani, M.; Gupta, G.; Verkhivker, G. AlphaFold2 Modeling and Molecular Dynamics Simulations of the Conformational Ensembles for the SARS-CoV-2 Spike Omicron JN.1, KP.2 and KP.3 Variants: Mutational Profiling of Binding Energetics Reveals Epistatic Drivers of the ACE2 Affinity and Escape Hotspots of Antibody Resistance. Viruses 2024, 16, 1458. doi: 10.3390/v16091458.

80. Verkhivker, G.; Alshahrani, M.; Gupta, G. Balancing Functional Tradeoffs between Protein Stability and ACE2 Binding in the SARS-CoV-2 Omicron BA.2, BA.2.75 and XBB Lineages: Dynamics-Based Network Models Reveal Epistatic Effects Modulating Compensatory Dynamic and Energetic Changes. Viruses 2023, 15, 1143. 10.3390/v15051143.

81. Verkhivker, G.; Agajanian, S.; Kassab, R.; Krishnan, K. Integrating Conformational Dynamics and Perturbation-Based Network Modeling for Mutational Profiling of Binding and Allostery in the SARS-CoV-2 Spike Variant Complexes with Antibodies: Balancing Local and Global Determinants of Mutational Escape Mechanisms. Biomolecules 2022, 12, 964. doi: 10.3390/biom12070964.

82. Alshahrani, M.; Parikh, V.; Foley, B.; Raisinghani, N.; Verkhivker, G. Quantitative Characterization and Prediction of the Binding Determinants and Immune Escape Hotspots for Groups of Broadly Neutralizing Antibodies Against Omicron Variants: Atomistic Modeling of the SARS-CoV-2 Spike Complexes with Antibodies. Biomolecules 2025, 15, 249. doi: 10.3390/biom15020249.

83. Alshahrani, M.; Parikh, V.; Foley, B.; Verkhivker, G. Integrative Computational Modeling of Distinct Binding Mechanisms for Broadly Neutralizing Antibodies Targeting SARS-CoV-2 Spike Omicron Variants: Balance of Evolutionary and Dynamic Adaptability in Shaping Molecular Determinants of Immune Escape. Viruses 2025, 17, 741; 10.3390/v17060741.

84. Yang, H.; Guo, H.; Wang, A.; Cao, L.; Fan, Q.; Jiang, J.; Wang, M.; Lin, L.; Ge, X.; Wang, H.; Zhang, R.; Liao, M.; Yan, R.; Ju, B.; Zhang, Z. Structural Basis for the Evolution and Antibody Evasion of SARS-CoV-2 BA.2.86 and JN.1 Subvariants. Nat Commun. 2024, 15, 7715. doi: 10.1038/s41467-024-51973-8.

85. Yajima, H.; Nomai, T.; Okumura, K.; Maenaka, K.; Ito, J.; Hashiguchi, T.; Sato, K.; Matsuno, K.; Nao, N.; Sawa, H.; Mizuma, K.; Li, J.; Kida, I.; Mimura, Y.; Ohari, Y.; Tanaka, S.; Tsuda, M.; Wang, L.; Oda, Y.; Ferdous, Z.; Shishido, K.; Mohri, H.; Iida, M.; Fukuhara, T.; Tamura, T.; Suzuki, R.; Suzuki, S.; Tsujino, S.; Ito, H.; Kaku, Y.; Misawa, N.; Plianchaisuk, A.; Guo, Z.; Hinay, A. A., Jr.; Usui, K.; Saikruang, W.; Lytras, S.; Uriu, K.; Yoshimura, R.; Kawakubo, S.; Nishumura, L.; Kosugi, Y.; Fujita, S.; M. Tolentino, J. E.; Chen, L.; Pan, L.; Li, W.; Yo, M. S.; Horinaka, K.; Suganami, M.; Chiba, M.; Yasuda, K.; Iida, K.; Strange, A. P.; Ohsumi, N.; Tanaka, S.; Ogawa, E.; Fukuda, T.; Osujo, R.; Yoshimura, K.; Sadamas, K.; Nagashima, M.; Asakura, H.; Yoshida, I.; Nakagawa, S.; Takayama, K.; Hashimoto, R.; Deguchi, S.; Watanabe, Y.; Nakata, Y.; Futatsusako, H.; Sakamoto, A.; Yasuhara, N.; Suzuki, T.; Kimura, K.; Sasaki, J.; Nakajima, Y.; Irie, T.; Kawabata, R.; Sasaki-Tabata, K.; Ikeda, T.; Nasser, H.; Shimizu, R.; Begum, M. M.; Jonathan, M.; Mugita, Y.; Leong, S.; Takahashi, O.; Ueno, T.; Motozono, C.; Toyoda, M.; Saito, A.; Kosaka, A.; Kawano, M.; Matsubara, N.; Nishiuchi, T.; Zahradnik, J.; Andrikopoulos, P.; Padilla-Blanco, M.; Konar, A. Molecular and Structural Insights into SARS-CoV-2 Evolution: From BA.2 to XBB Subvariants. mBio. 2024, 15, e0322023. doi: 10.1128/mbio.03220-23.

86. Xue, S.; Han, Y.; Wu, F.; Wang, Q. Mutations in the SARS-CoV-2 Spike Receptor Binding Domain and Their Delicate Balance between ACE2 Affinity and Antibody Evasion. Protein Cell. 2024, 15, 403–418. doi: 10.1093/procel/pwae007.

87. Tulsian, N. K.; Palur, R. V.; Qian, X.; Gu, Y.; D/O Shunmuganathan, B.; Samsudin, F.; Wong, Y. H.; Lin, J.; Purushotorman, K.; Kozma, M. M.; Wang, B.; Lescar, J.; Wang, C.-I.; Gupta, R. K.; Bond, P. J.; MacAry, P. A. Defining Neutralization and Allostery by Antibodies against COVID-19 Variants. Nat Commun. 2023, 14, 6967. doi: 10.1038/s41467-023-42408-x.

88. Tortorici, M. A.; Czudnochowski, N.; Starr, T. N.; Marzi, R.; Walls, A. C.; Zatta, F.; Bowen, J. E.; Jaconi, S.; Di Iulio, J.; Wang, Z.; De Marco, A.; Zepeda, S. K.; Pinto, D.; Liu, Z.; Beltramello, M.; Bartha, I.; Housley, M. P.; Lempp, F. A.; Rosen, L. E.; Dellota, E., Jr; Kaiser, H.; Montiel-Ruiz, M.; Zhou, J.; Addetia, A.; Guarino, B.; Culap, K.; Sprugasci, N.; Saliba, C.; Vetti, E.; Giacchetto-Sasselli, I.; Fregni, C. S.; Abdelnabi, R.; Foo, S.-Y. C.; Havenar-Daughton, C.; Schmid, M. A.; Benigni, F.; Cameroni, E.; Neyts, J.; Telenti, A.; Virgin, H. W.; Whelan, S. P. J.; Snell, G.; Bloom, J. D.; Corti, D.; Veesler, D.; Pizzuto, M. S. Broad Sarbecovirus Neutralization by a Human Monoclonal Antibody. Nature 2021, 597, 103–108. doi:10.1038/s41586-021-03817-4.

89. Feng, Y.; Yuan, M.; Powers, J. M.; Hu, M.; Munt, J. E.; Arunachalam, P. S.; Leist, S. R.; Bellusci, L.; Kim, J.; Sprouse, K. R.; Adams, L. E.; Sundaramurthy, S.; Zhu, X.; Shirreff, L. M.; Mallory, M. L.; Scobey, T. D.; Moreno, A.; O’Hagan, D. T.; Kleanthous, H.; Villinger, F. J.; Veesler, D.; King, N. P.; Suthar, M. S.; Khurana, S.; Baric, R. S.; Wilson, I. A.; Pulendran, B. Broadly Neutralizing Antibodies against Sarbecoviruses Generated by Immunization of Macaques with an AS03-Adjuvanted COVID-19 Vaccine. Sci Transl Med. 2023, 15, eadg7404. doi: 10.1126/scitranslmed.adg7404.

90. Wang, Q.; Guo, Y.; Mellis, I. A.; Wu, M.; Mohri, H.; Gherasim, C.; Valdez, R.; Purpura, L. J.; Yin, M. T.; Gordon, A.; Ho, D. D. Antibody Evasiveness of SARS-CoV-2 Subvariants KP.3.1.1 and XEC. Cell Rep. 2025, 44, 115543. doi: 10.1016/j.celrep.2025.115543.

91. Rose, P. W.; Prlic, A.; Altunkaya, A.; Bi, C.; Bradley, A. R.; Christie, C. H.; Costanzo, L. D.; Duarte, J. M.; Dutta, S.; Feng, Z.; Green, R. K.; Goodsell, D. S.; Hudson, B.; Kalro, T.; Lowe, R.; Peisach, E.; Randle, C.; Rose, A. S.; Shao, C.; Tao, Y. P.; Valasatava, Y.; Voigt, M.; Westbrook, J. D.; Woo, J.; Yang, H.; Young, J. Y.; Zardecki, C.; Berman, H. M.; Burley, S. K. The RCSB protein data bank: integrative view of protein, gene and 3D structural information. Nucleic Acids Res. 2017, 45, D271–D281. doi: 10.1093/nar/gkw1000.

92. Kmiecik, S.; Kolinski, A. Characterization of protein-folding pathways by reduced-space modeling. Proc. Natl. Acad. Sci. U. S. A. 2007, 104, 12330–12335. doi: 10.1073/pnas.0702265104.

93. Kmiecik, S.; Gront, D.; Kolinski, M.; Wieteska, L.; Dawid, A.E.; Kolinski, A. Coarse-grained protein models and their applications. Chem. Rev. 2016, 116, 7898–7936. doi: 10.1021/acs.chemrev.6b00163.

94. Kmiecik, S.; Kouza, M.; Badaczewska-Dawid, A.E.; Kloczkowski, A.; Kolinski, A. Modeling of protein structural flexibility and large-scale dynamics: Coarse-grained simulations and elastic network models. Int. J. Mol. Sci. 2018, 19, 3496. doi: 10.3390/ijms19113496.

95. Ciemny, M.P.; Badaczewska-Dawid, A.E.; Pikuzinska, M.; Kolinski, A.; Kmiecik, S. Modeling of disordered protein structures using monte carlo simulations and knowledge-based statistical force fields. Int. J. Mol. Sci. 2019, 20, 606. doi: 10.3390/ijms20030606.

96. Kurcinski, M.; Oleniecki, T.; Ciemny, M.P.; Kuriata, A.; Kolinski, A.; Kmiecik, S. CABS-flex standalone: A simulation environment for fast modeling of protein flexibility. Bioinformatics 2019, 35, 694–695. doi: 10.1093/bioinformatics/bty685.

97. Badaczewska-Dawid, A. E.; Kolinski, A.; Kmiecik, S. Protocols for fast simulations of protein structure flexibility using CABS-Flex and SURPASS. Methods Mol. Biol. 2020, 2165, 337–353. doi: 10.1007/978-1-0716-0708-4_20.

98. Marti-Renom, M. A.; Stuart, A. C.; Fiser, A.; Sanchez, R.; Melo, F.; Sali, A. Comparative protein structure modeling of genes and genomes. Annu. Rev. Biophys. Biomol. Struct. 2000, 29, 291–325.

99. Fernandez-Fuentes, N.; Zhai, J.; Fiser, A. ArchPRED: A template based loop structure prediction server. Nucleic Acids Res. 2006, 34, W173–W176. 10.1093/nar/gkl113.

100. Krivov, V.P.,B.F.; Shapovalov, M.V.; Dunbrack, R.L., Jr. Improved prediction of protein side-chain conformations with SCWRL4. Proteins 2009, 77, 778–795. doi10.1002/prot.22488.

101. Søndergaard C. R.; Olsson M. H.; Rostkowski M.; Jensen J. H. Improved treatment of ligands and coupling effects in empirical calculation and rationalization of pKa values. J. Chem. Theory Comput. 2011, 7, 2284–2295. 10.1021/ct200133y.

102. Olsson M. H.; Søndergaard C. R.; Rostkowski M.; Jensen J. H. PROPKA3: consistent treatment of internal and surface residues in empirical pKa predictions. J. Chem. Theory Comput. 2011, 7, 525–537. 10.1021/ct100578z.

103. Lee, J.; Cheng, X.; Swails, J. M.; Yeom, M. S.; Eastman, P. K.; Lemkul, J. A.; Wei, S.; Buckner, J.; Jeong, J. C.; Qi, Y.; Jo, S.; Pande, V. S.; Case, D. A.; Brooks, C. L., III; MacKerell, A. D., Jr.; Klauda, J. B.; Im, W. CHARMM-GUI Input Generator for NAMD, GROMACS, AMBER, OpenMM, and CHARMM/OpenMM Simulations Using the CHARMM36 Additive Force Field. J Chem Theory Comput. 2016, 12, 405–413. doi: 10.1021/acs.jctc.5b00935.

104. Park, S.-J.; Lee, J.; Qi, Y.; Kern, N. R.; Lee, H. S.; Jo, S.; Joung, I.; Joo, K.; Lee, J.; Im, W. CHARMM-GUI Glycan Modeler for Modeling and Simulation of Carbohydrates and Glycoconjugates. Glycobiology. 2019, 29, 320–331. doi: 10.1093/glycob/cwz003.

105. Phillips, J.C.; Hardy, D.J.; Maia, J.D.C.; Stone, J.E.; Ribeiro, J.V.; Bernardi, R.C.; Buch, R.; Fiorin, G.; Hénin, J.; Jiang, W.;, et al. Scalable Molecular Dynamics on CPU and GPU Architectures with NAMD. J. Chem. Phys. 2020, 153, 044130. 10.1063/5.0014475.

106. Huang, J.; Rauscher, S.; Nawrocki, G.; Ran, T.; Feig, M.; de Groot, B.L.; Grubmüller, H.; MacKerell, A.D., Jr. CHARMM36m: An improved force field for folded and intrinsically disordered proteins. Nat. Methods 2017, 14, 71–73. 10.1038/nmeth.4067.

107. Jorgensen, W.L.; Chandrasekhar, J.; Madura, J.D.; Impey, R.W.; Klein, M.L. Comparison of Simple Potential Functions for Simulating Liquid Water. J. Chem. Phys. 1983, 79, 926–935. 10.1063/1.445869.

108. Ross, G.A.; Rustenburg, A.S.; Grinaway, P.B.; Fass, J.; Chodera, J.D. Biomolecular Simulations under Realistic Macroscopic Salt Conditions. J. Phys. Chem. B 2018, 122, 5466–5486. 10.1021/acs.jpcb.7b11734.

109. Hoover, W. G. Canonical Dynamics: Equilibrium Phase-Space Distributions. Phys. Rev. A 1985, 31, 1695–1697. doi: 10.1103/physreva.31.1695.

110. Parrinello, M.; Rahman, A. Polymorphic transitions in single crystals: a new molecular dynamics method. J. Appl. Phys. 1981, 52, 7182–7190. doi: 10.1063/1.328693

111. Di Pierro, M.; Elber, R.; Leimkuhler, B. A Stochastic Algorithm for the Isobaric-Isothermal Ensemble with Ewald Summations for All Long Range Forces. J. Chem. Theory Comput. 2015, 11, 5624–5637. 10.1021/acs.jctc.5b00648.

112. Martyna, G.J.; Tobias, D.J.; Klein, M.L. Constant pressure molecular dynamics algorithms. J. Chem. Phys. 1994, 101, 4177–4189. 10.1063/1.467468.

113. Feller, S.E.; Zhang, Y.; Pastor, R.W.; Brooks, B.R. Constant pressure molecular dynamics simulation: The Langevin piston method. J. Chem. Phys. 1995, 103, 4613–4621. 10.1063/1.470648.

114. Davidchack, R.L.; Handel, R.; Tretyakov, M.V. Langevin thermostat for rigid body dynamics. J. Chem. Phys. 2009, 130, 234101. 10.1063/1.3149788.

115. Dehouck, Y.; Kwasigroch, J. M.; Rooman, M.; Gilis, D. BeAtMuSiC: Prediction of changes in protein-protein binding affinity on mutations. Nucleic Acids Res. 2013, 41, W333–W339. doi: 10.1093/nar/gkt450.

116. Dehouck, Y.; Gilis, D.; Rooman, M. A new generation of statistical potentials for proteins. Biophys. J. 2006, 90, 4010–4017. doi: 10.1529/biophysj.105.079434.

117. Dehouck, Y.; Grosfils, A.; Folch, B.; Gilis, D.; Bogaerts, P.; Rooman, M. Fast and accurate predictions of protein stability changes upon mutations using statistical potentials and neural networks:PoPMuSiC-2.0. Bioinformatics 2009, 25, 2537–2543. doi:10.1093/bioinformatics/btp445.

118. Tsishyn, M.; Pucci, F.; Rooman, M. Quantification of Biases in Predictions of Protein– Protein Binding Affinity Changes upon Mutations. Brief Bioinform. 2023, 25, bbad491. doi: 10.1093/bib/bbad491.

119. Pahari, S.; Li, G.; Murthy, A. K.; Liang, S.; Fragoza, R.; Yu, H.; Alexov, E. SAAMBE-3D: Predicting Effect of Mutations on Protein–Protein Interactions. Int J Mol Sci. 2020, 21, 2563. doi: 10.3390/ijms21072563.

120. Rimal, P.; Paul, S.; Panday, S.; Alexov, E. Further Development of SAMPDI-3D: A Machine Learning Method for Predicting Binding Free Energy Changes Caused by Mutations in Either Protein or DNA. Genes 2025, 16, 101. doi: 10.3390/genes16010101.

121. Brinda, K. V.; Vishveshwara, S. A Network Representation of Protein Structures: Implications for Protein Stability. Biophys. J. 2005, 89, 4159–4170. 10.1529/biophysj.105.067231

122. Vijayabaskar, M. S.; Vishveshwara, S. Interaction Energy Based Protein Structure Networks. Biophys. J. 2010, 99, 3704–3715. 10.1016/j.bpj.2010.08.079.

123. Sethi, A.; Eargle, J.; Black, A. A.; Luthey-Schulten, Z. Dynamical Networks in tRNA:Protein Complexes. Proc. Natl. Acad. Sci. U.S.A. 2009, 106, 6620–6625. 10.1073/pnas.0810961106.

124. Piovesan, D.; Minervini, G.; Tosatto, S. C. The RING 2.0 Web Server for High Quality Residue Interaction Networks. Nucleic Acids Res. 2016, 44, W367–W374. 10.1093/nar/gkw315.

125. Clementel, D.; Del Conte, A.; Monzon, A. M.; Camagni, G. F.; Minervini, G.; Piovesan, D.; Tosatto, S. C. E. RING 3.0: Fast Generation of Probabilistic Residue Interaction Networks from Structural Ensembles. Nucleic Acids Res. 2022, 50, W651–W656. 10.1093/nar/gkac365.

126. Del Conte, A.; Camagni, G. F.; Clementel, D.; Minervini, G.; Monzon, A. M.; Ferrari, C.; Piovesan, D.; Tosatto, S. C. E. RING 4.0: Faster Residue Interaction Networks with Novel Interaction Types across over 35,000 Different Chemical Structures. Nucleic Acids Res. 2024, 52, W306–W312. 10.1093/nar/gkae337.

127. Floyd, R. W. Algorithm 97: Shortest Path. Commun. ACM 1962, 5, 345. 10.1145/367766.368168.

128. Hagberg, A. A.; Schult, D. A.; Swart, P. J. Exploring Network Structure, Dynamics, and Function Using NetworkX. In Proceedings of the 7th Python in Science Conference (SciPy2008); Varoquaux, G., Vaught, T., Millman, J., Eds.; Pasadena, 2008; pp 11–15.

129. Javanmardi, K.; Segall-Shapiro, T. H.; Chou, C.-W.; Boutz, D. R.; Olsen, R. J.; Xie, X.; Xia, H.; Shi, P.-Y.; Johnson, C. D.; Annapareddy, A.; Weaver, S.; Musser, J. M.; Ellington, A. D.; Finkelstein, I. J.; Gollihar, J. D. Antibody Escape and Cryptic Cross-Domain Stabilization in the SARS-CoV-2 Omicron Spike Protein. Cell Host Microbe. 2022, 30, 1242–1254.e6. doi: 10.1016/j.chom.2022.07.016.

130. Cui, L.; Li, T.; Lan, M.; Zhou, M.; Xue, W.; Zhang, S.; Wang, H.; Hong, M.; Zhang, Y.; Yuan, L.; Sun, H.; Ye, J.; Zheng, Q.; Guan, Y.; Gu, Y.; Xia, N.; Li, S. A Cryptic Site in Class 5 Epitope of SARS-CoV-2 RBD Maintains Highly Conservation across Natural Isolates. iScience 2024, 27, 110208. doi: 10.1016/j.isci.2024.110208.

131. Arora, P.; Kempf, A.; Nehlmeier, I.; Schulz, S. R.; Cossmann, A.; Stankov, M. V.; Jäck, H.-M.; Behrens, G. M. N.; Pöhlmann, S.; Hoffmann, M. Augmented Neutralisation Resistance of Emerging Omicron Subvariants BA.2.12.1, BA.4, and BA.5. Lancet Infect Dis. 2022, 22, 1117–1118. doi: 10.1016/S1473-3099(22)00422-4.

132. Weng, C.; Faure, A. J.; Escobedo, A.; Lehner, B. The Energetic and Allosteric Landscape for KRAS Inhibition. Nature 2024, 626, 643–652. 10.1038/s41586-023-06954-0.

133. Bendel, A. M.; Faure, A. J.; Klein, D.; Shimada, K.; Lyautey, R.; Schiffelholz, N.; Kempf, G.; Cavadini, S.; Lehner, B.; Diss, G. The Genetic Architecture of Protein Interaction Affinity and Specificity. Nat Commun. 2024, 15, 8868. doi: 10.1038/s41467-024-53195-4.

134. Martí-Aranda, A.; Lehner, B. The Evolution of Allostery in a Protein Family, bioRxiv 2025. 10.1101/2025.06.20.660748.

135. Liao, X.; Lehner, B. Allostery Is a Widespread Cause of Loss-of-Function Variant Pathogenicity, bioRxiv 2025. 10.1101/2025.06.20.660737.

