## Supplemental Figures S1-S3, Tables S1-S6 for "Exploring Dynamic Modulation of Binding, Allostery and Immune Resistance in the SARS-CoV-2 Spike Complexes with Classes of Antibodies Targeting Cryptic Binding Sites: Antibody-Specific Augmentations of Conserved Allosteric Architecture Can Influence Evolution of Viral Escape"

**Supp****lementary Materials**

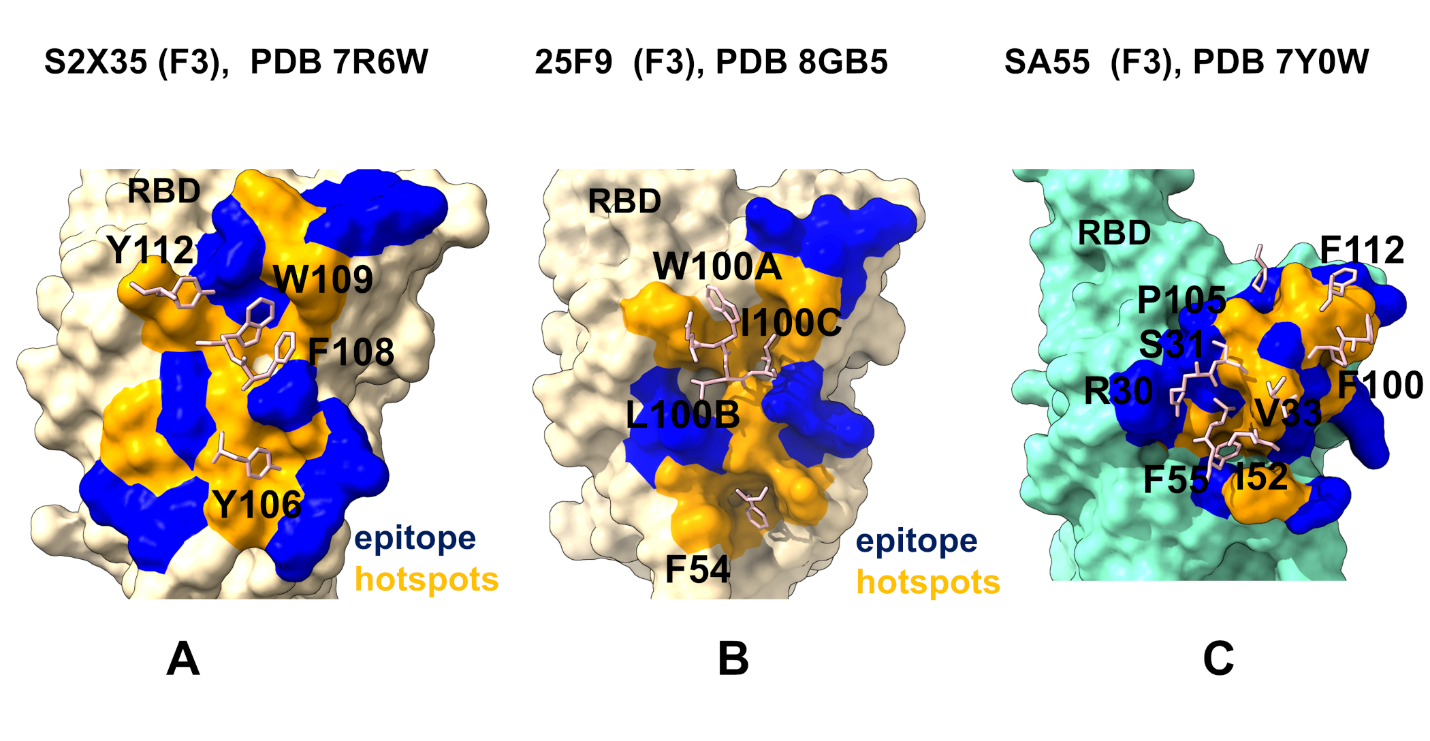

**Figure S1.** Structural maps of the close-ups of binding interactions formed by energetic hotspots of class 4 S2X35 antibody with the RBD (A), class 4 25F9 antibody with RBD (B), and class 4 SA55 antibody with BA.1 RBD (C). The hotspots from the heavy chain of class 4 antibodies are shown and annotated in pink sticks on panels (A-C). The RBD is shown in wheat-colored surface. The binding epitope residues are in blue surface and binding energy hotspots on the RBD are in orange surface.

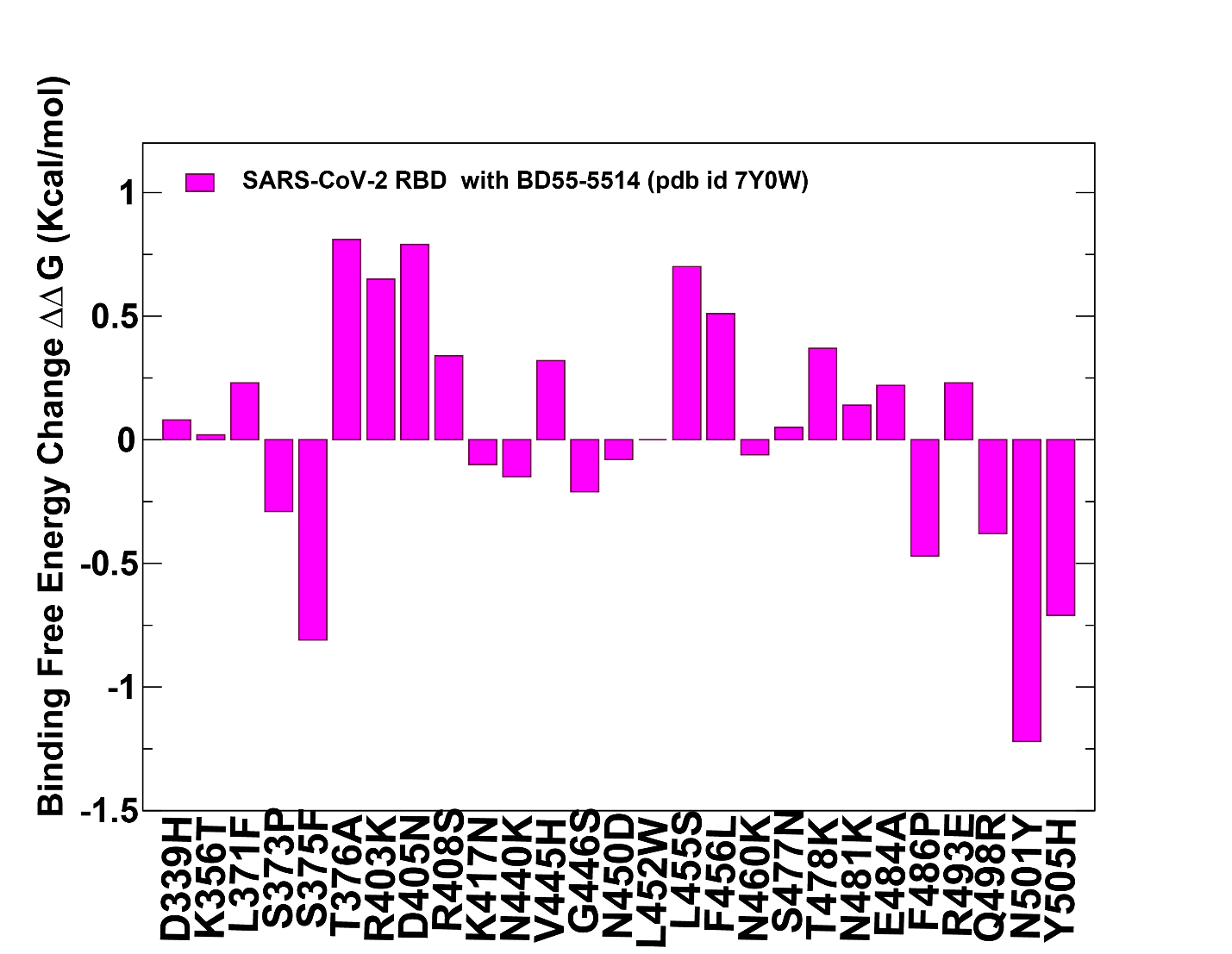

**Figure S2.** Structure-based mutational profiling of the S complexes with BD55-5514 (SA55; class 4) antibodies. The mutational screening evaluates binding energy changes induced by BA.2.86/JN.1/KP.2/KP.3 mutations in the RBD-antibody complexes. Mutational profiling of the S complex with SA55 (pdb id 7Y0W). The binding free energy changes are shown in magenta-colored filled bars. The positive binding free energy values ΔΔG correspond to destabilizing changes and negative binding free energy changes are associated with stabilizing changes.

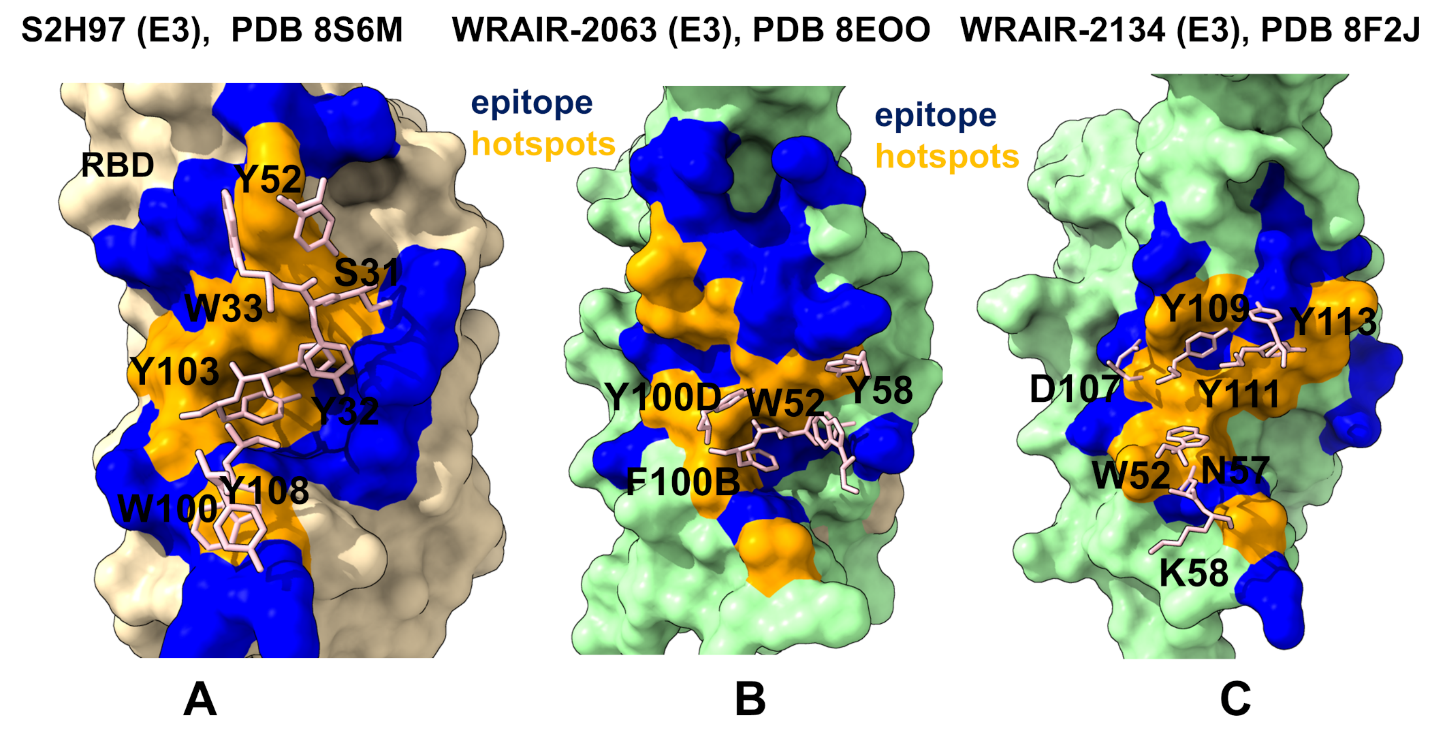

**Figure S1.** Structural maps of the close-us of binding interactions formed by energetic hotspots of class 5 S2H97 antibody with the RBD (A), class 5 WRAIR-063 antibody with RBD (B), and class 5 WRAIR antibody with RBD (C). The hotspots from the heavy chain of class 5 antibodies are shown and annotated in pink sticks on panels (A-C). The RBD is shown in wheat-colored surface. The binding epitope residues are in blue surface and binding energy hotspots on the RBD are in orange surface.

**Table S1**. The list of the intermolecular contacts in the structure of the Class 4 S2X35 antibody complex with RBD (pdb id 7R6W). The interfacial contacts in the structure are defined by counting the number of interatomic contacts within a 5.5 Å distance threshold between atoms of the interacting proteins.

| **RBD Residue** | **RBD Residue Number** | **RBD chain** | **Ab Residue** | **Ab Residue Number** | **Ab chain** |
| --- | --- | --- | --- | --- | --- |
| LEU | 368 | R | TYR | 106 | H |
| TYR | 369 | R | TYR | 54 | H |
| TYR | 369 | R | TYR | 106 | H |
| TYR | 369 | R | GLY | 105 | H |
| TYR | 369 | R | LEU | 104 | H |
| TYR | 369 | R | LYS | 55 | H |
| ASN | 370 | R | TYR | 54 | H |
| ASN | 370 | R | TYR | 106 | H |
| ASN | 370 | R | LYS | 55 | H |
| SER | 371 | R | LYS | 55 | H |
| SER | 371 | R | TYR | 106 | H |
| ALA | 372 | R | LYS | 55 | H |
| SER | 373 | R | LYS | 55 | H |
| PHE | 374 | R | LYS | 55 | H |
| PHE | 374 | R | ASN | 57 | H |
| PHE | 374 | R | TYR | 106 | H |
| SER | 375 | R | PHE | 108 | H |
| SER | 375 | R | TRP | 109 | H |
| SER | 375 | R | TYR | 106 | H |
| SER | 375 | R | ASN | 57 | H |
| SER | 375 | R | SER | 52 | H |
| SER | 375 | R | ASP | 107 | H |
| SER | 375 | R | LYS | 55 | H |
| THR | 376 | R | PHE | 108 | H |
| THR | 376 | R | TRP | 109 | H |
| THR | 376 | R | TYR | 106 | H |
| THR | 376 | R | ASP | 107 | H |
| PHE | 377 | R | TYR | 106 | H |
| PHE | 377 | R | LEU | 104 | H |
| PHE | 377 | R | VAL | 103 | H |
| PHE | 377 | R | ASP | 107 | H |
| PHE | 377 | R | GLY | 105 | H |
| LYS | 378 | R | ASP | 107 | H |
| LYS | 378 | R | TYR | 106 | H |
| LYS | 378 | R | GLY | 105 | H |
| LYS | 378 | R | LEU | 104 | H |
| LYS | 378 | R | VAL | 103 | H |
| LYS | 378 | R | TYR | 101 | H |
| CYS | 379 | R | GLY | 105 | H |
| CYS | 379 | R | LEU | 104 | H |
| CYS | 379 | R | VAL | 103 | H |
| TYR | 380 | R | TYR | 101 | H |
| TYR | 380 | R | LEU | 104 | H |
| TYR | 380 | R | VAL | 103 | H |
| GLY | 381 | R | LEU | 104 | H |
| VAL | 382 | R | LEU | 104 | H |
| SER | 383 | R | LEU | 104 | H |
| PRO | 384 | R | GLY | 105 | H |
| PRO | 384 | R | LEU | 104 | H |
| ARG | 403 | R | TYR | 33 | L |
| ARG | 403 | R | SER | 95 | L |
| GLY | 404 | R | TRP | 109 | H |
| GLY | 404 | R | TYR | 112 | H |
| ASP | 405 | R | TYR | 112 | H |
| ASP | 405 | R | SER | 95 | L |
| ASP | 405 | R | TYR | 93 | L |
| ASP | 405 | R | TRP | 109 | H |
| ASP | 405 | R | TYR | 33 | L |
| ASP | 405 | R | SER | 96 | L |
| ASP | 405 | R | SER | 100 | L |
| ASP | 405 | R | ASP | 94 | L |
| GLU | 406 | R | TYR | 112 | H |
| VAL | 407 | R | ASP | 107 | H |
| VAL | 407 | R | TRP | 109 | H |
| ARG | 408 | R | TYR | 112 | H |
| ARG | 408 | R | TYR | 113 | H |
| ARG | 408 | R | GLY | 32 | L |
| ARG | 408 | R | ASP | 34 | L |
| ARG | 408 | R | TRP | 109 | H |
| ARG | 408 | R | TYR | 33 | L |
| GLY | 496 | R | LEU | 97 | L |
| GLN | 498 | R | LEU | 97 | L |
| THR | 500 | R | LEU | 97 | L |
| ASN | 501 | R | LEU | 97 | L |
| GLY | 502 | R | GLY | 99 | L |
| GLY | 502 | R | SER | 96 | L |
| GLY | 502 | R | SER | 100 | L |
| GLY | 502 | R | LEU | 97 | L |
| GLY | 502 | R | SER | 98 | L |
| VAL | 503 | R | GLY | 99 | L |
| VAL | 503 | R | SER | 100 | L |
| VAL | 503 | R | PHE | 108 | H |
| VAL | 503 | R | TRP | 109 | H |
| VAL | 503 | R | ASN | 59 | H |
| GLY | 504 | R | SER | 95 | L |
| GLY | 504 | R | GLY | 99 | L |
| GLY | 504 | R | SER | 96 | L |
| GLY | 504 | R | SER | 100 | L |
| GLY | 504 | R | TRP | 109 | H |
| TYR | 505 | R | LEU | 97 | L |
| TYR | 505 | R | SER | 95 | L |
| TYR | 505 | R | SER | 96 | L |
| TYR | 505 | R | SER | 100 | L |
| TYR | 508 | R | PHE | 108 | H |
| TYR | 508 | R | TRP | 109 | H |

**Table S2.** The list of the intermolecular contacts in the structure of the Class 4 25F9 antibody complex with RBD (pdb id 8GB5). The interfacial contacts in the structure are defined by counting the number of interatomic contacts within a 5.5 Å distance threshold between atoms of the interacting proteins.

| **RBD Residue** | **RBD Residue Number** | **RBD chain** | **Ab Residue** | **Ab Residue Number** | **Ab chain** |
| --- | --- | --- | --- | --- | --- |
| TYR | 365 | A | PHE | 54 | B |
| SER | 366 | A | PHE | 54 | B |
| TYR | 369 | A | LYS | 52 | B |
| TYR | 369 | A | PHE | 54 | B |
| TYR | 369 | A | GLY | 55 | B |
| TYR | 369 | A | THR | 56 | B |
| SER | 373 | A | ARG | 93 | C |
| PHE | 374 | A | ILE | 100 | B |
| SER | 375 | A | ILE | 100 | B |
| SER | 375 | A | LEU | 100 | B |
| THR | 376 | A | ASP | 53 | B |
| THR | 376 | A | LEU | 100 | B |
| THR | 376 | A | ILE | 100 | B |
| PHE | 377 | A | ASP | 53 | B |
| PHE | 377 | A | PHE | 54 | B |
| LYS | 378 | A | ASP | 53 | B |
| LYS | 378 | A | PHE | 54 | B |
| LYS | 378 | A | LEU | 100 | B |
| CYS | 379 | A | ASP | 53 | B |
| CYS | 379 | A | PHE | 54 | B |
| TYR | 380 | A | LEU | 100 | B |
| PRO | 384 | A | LYS | 52 | B |
| PRO | 384 | A | ASP | 53 | B |
| PRO | 384 | A | PHE | 54 | B |
| LEU | 387 | A | PHE | 54 | B |
| VAL | 407 | A | LEU | 100 | B |
| ARG | 408 | A | LEU | 100 | B |
| ARG | 408 | A | GLU | 100 | B |
| ALA | 411 | A | LEU | 100 | B |
| THR | 500 | A | ARG | 66 | C |
| ASN | 501 | A | ALA | 29 | C |
| ASN | 501 | A | GLY | 30 | C |
| ASN | 501 | A | ARG | 66 | C |
| GLY | 502 | A | ALA | 29 | C |
| GLY | 502 | A | GLY | 30 | C |
| GLY | 502 | A | TYR | 32 | C |
| GLY | 502 | A | ARG | 66 | C |
| VAL | 503 | A | ALA | 29 | C |
| VAL | 503 | A | GLY | 30 | C |
| VAL | 503 | A | HIS | 31 | C |
| VAL | 503 | A | TYR | 32 | C |
| VAL | 503 | A | ALA | 100 | B |
| GLY | 504 | A | TYR | 32 | C |
| TYR | 505 | A | TYR | 32 | C |
| GLN | 506 | A | ALA | 29 | C |
| GLN | 506 | A | GLY | 30 | C |

**Table S3.** The list of the intermolecular contacts in the structure of the Class 4 SA55 antibody complex with RBD (pdb id 7Y0W). The interfacial contacts in the structure are defined by counting the number of interatomic contacts within a 5.5 Å distance threshold between atoms of the interacting proteins.

| **RBD Residue** | **RBD Residue Number** | **RBD chain** | **Ab Residue** | **Ab Residue Number** | **Ab chain** |
| --- | --- | --- | --- | --- | --- |
| PRO | 373 | R | LEU | 94 | B |
| PHE | 374 | R | PHE | 55 | A |
| PHE | 374 | R | THR | 57 | A |
| THR | 376 | R | PHE | 55 | A |
| ARG | 403 | R | PRO | 105 | A |
| ARG | 403 | R | ASN | 106 | A |
| GLY | 404 | R | ARG | 30 | A |
| GLY | 404 | R | LEU | 54 | A |
| GLY | 404 | R | PHE | 55 | A |
| ASP | 405 | R | THR | 28 | A |
| ASP | 405 | R | ARG | 30 | A |
| ASP | 405 | R | SER | 31 | A |
| ASP | 405 | R | LEU | 54 | A |
| GLU | 406 | R | ARG | 30 | A |
| VAL | 407 | R | ARG | 30 | A |
| VAL | 407 | R | LEU | 54 | A |
| VAL | 407 | R | PHE | 55 | A |
| ARG | 408 | R | ARG | 30 | A |
| ASN | 437 | R | ASP | 93 | B |
| ASN | 439 | R | TYR | 91 | B |
| ASN | 439 | R | ASP | 93 | B |
| LYS | 440 | R | ASP | 93 | B |
| VAL | 445 | R | HIS | 53 | B |
| TYR | 495 | R | PRO | 105 | A |
| SER | 496 | R | PRO | 105 | A |
| ARG | 498 | R | TYR | 49 | B |
| ARG | 498 | R | PHE | 112 | A |
| PRO | 499 | R | ASP | 50 | B |
| PRO | 499 | R | TYR | 91 | B |
| PRO | 499 | R | PHE | 100 | A |
| PRO | 499 | R | PRO | 101 | A |
| THR | 500 | R | TYR | 49 | B |
| THR | 500 | R | ASP | 50 | B |
| THR | 500 | R | PHE | 100 | A |
| THR | 500 | R | PRO | 101 | A |
| THR | 500 | R | ASN | 102 | A |
| THR | 500 | R | GLY | 103 | A |
| THR | 500 | R | ASP | 104 | A |
| THR | 500 | R | PHE | 112 | A |
| TYR | 501 | R | PRO | 101 | A |
| TYR | 501 | R | ASN | 102 | A |
| TYR | 501 | R | GLY | 103 | A |
| TYR | 501 | R | ASP | 104 | A |
| TYR | 501 | R | PRO | 105 | A |
| TYR | 501 | R | PHE | 112 | A |
| GLY | 502 | R | SER | 31 | A |
| GLY | 502 | R | HIS | 32 | A |
| GLY | 502 | R | PRO | 101 | A |
| GLY | 502 | R | ASN | 102 | A |
| GLY | 502 | R | GLY | 103 | A |
| GLY | 502 | R | ASP | 104 | A |
| VAL | 503 | R | SER | 31 | A |
| VAL | 503 | R | HIS | 32 | A |
| VAL | 503 | R | VAL | 33 | A |
| VAL | 503 | R | ILE | 52 | A |
| VAL | 503 | R | LEU | 54 | A |
| VAL | 503 | R | PHE | 55 | A |
| VAL | 503 | R | PRO | 95 | B |
| VAL | 503 | R | PRO | 101 | A |
| VAL | 503 | R | ASN | 102 | A |
| GLY | 504 | R | ARG | 30 | A |
| GLY | 504 | R | SER | 31 | A |
| GLY | 504 | R | HIS | 32 | A |
| GLY | 504 | R | LEU | 54 | A |
| HIS | 505 | R | SER | 31 | A |
| HIS | 505 | R | HIS | 32 | A |
| HIS | 505 | R | GLY | 103 | A |
| HIS | 505 | R | ASP | 104 | A |
| HIS | 505 | R | PRO | 105 | A |
| GLN | 506 | R | TYR | 91 | B |
| GLN | 506 | R | ASP | 93 | B |
| GLN | 506 | R | PRO | 101 | A |
| TYR | 508 | R | LEU | 54 | A |
| TYR | 508 | R | PHE | 55 | A |

**Table S4.** The list of the intermolecular contacts in the structure of the Class 5 S2H97 antibody complex with RBD (pdb id 8S6M). The interfacial contacts in the structure are defined by counting the number of interatomic contacts within a 5.5 Å distance threshold between atoms of the interacting proteins.

| **RBD Residue** | **RBD Residue Number** | **RBD chain** | **Ab Residue** | **Ab Residue Number** | **Ab chain** |
| --- | --- | --- | --- | --- | --- |
| TRP | 353 | R | TYR | 32 | M |
| ARG | 355 | R | TYR | 32 | M |
| ARG | 355 | R | HIS | 102 | I |
| ARG | 355 | R | TYR | 103 | I |
| ARG | 355 | R | TYR | 105 | I |
| ARG | 357 | R | ASP | 52 | M |
| ARG | 357 | R | ASN | 55 | M |
| THR | 393 | R | TYR | 51 | M |
| ASN | 394 | R | TYR | 51 | M |
| ASN | 394 | R | THR | 104 | I |
| TYR | 396 | R | TYR | 103 | I |
| PRO | 426 | R | SER | 31 | I |
| PRO | 426 | R | HIS | 102 | I |
| PRO | 426 | R | TYR | 103 | I |
| ASP | 427 | R | SER | 28 | I |
| ASP | 427 | R | THR | 30 | I |
| ASP | 427 | R | SER | 31 | I |
| ASP | 428 | R | TYR | 27 | I |
| ASP | 428 | R | SER | 28 | I |
| ASP | 428 | R | SER | 31 | I |
| ASP | 428 | R | TYR | 32 | I |
| ASP | 428 | R | HIS | 102 | I |
| PHE | 429 | R | HIS | 102 | I |
| PHE | 429 | R | TYR | 103 | I |
| THR | 430 | R | TYR | 103 | I |
| SER | 459 | R | ASP | 55 | I |
| SER | 459 | R | ASP | 57 | I |
| SER | 459 | R | ARG | 59 | I |
| LYS | 460 | R | ASP | 55 | I |
| LYS | 460 | R | ASP | 57 | I |
| LYS | 462 | R | TRP | 33 | I |
| LYS | 462 | R | TYR | 52 | I |
| LYS | 462 | R | ASP | 55 | I |
| LYS | 462 | R | ASP | 57 | I |
| LYS | 462 | R | ARG | 59 | I |
| PRO | 463 | R | SER | 31 | I |
| PRO | 463 | R | TYR | 52 | I |
| PRO | 463 | R | HIS | 102 | I |
| PHE | 464 | R | TYR | 32 | M |
| PHE | 464 | R | HIS | 102 | I |
| PHE | 464 | R | TYR | 103 | I |
| GLU | 465 | R | TYR | 32 | M |
| GLU | 465 | R | ARG | 59 | I |
| ARG | 466 | R | GLY | 30 | M |
| ARG | 466 | R | TYR | 32 | M |
| SER | 514 | R | TYR | 103 | I |
| PHE | 515 | R | TYR | 103 | I |
| GLU | 516 | R | SER | 101 | I |
| GLU | 516 | R | HIS | 102 | I |
| GLU | 516 | R | TYR | 103 | I |
| GLU | 516 | R | THR | 104 | I |
| LEU | 518 | R | TYR | 51 | M |
| LEU | 518 | R | TRP | 100 | I |
| LEU | 518 | R | SER | 101 | I |
| LEU | 518 | R | THR | 104 | I |
| HIS | 519 | R | ARG | 56 | M |
| HIS | 519 | R | PRO | 57 | M |
| HIS | 519 | R | SER | 58 | M |
| HIS | 519 | R | GLY | 59 | M |
| HIS | 519 | R | TRP | 100 | I |
| HIS | 519 | R | TYR | 108 | I |
| ALA | 520 | R | TYR | 51 | M |
| ALA | 520 | R | ARG | 56 | M |
| ALA | 520 | R | PRO | 57 | M |
| ALA | 520 | R | SER | 58 | M |
| ALA | 520 | R | TRP | 100 | I |
| PRO | 521 | R | ARG | 56 | M |
| PRO | 521 | R | PRO | 57 | M |
| PRO | 521 | R | SER | 58 | M |

**Table S5.** The list of the intermolecular contacts in the structure of the Class 5 WRAIR-2063 antibody complex with RBD (pdb id 8EOO). The interfacial contacts in the structure are defined by counting the number of interatomic contacts within a 5.5 Å distance threshold between atoms of the interacting proteins.

| **RBD Residue** | **RBD Residue Number** | **RBD chain** | **Ab Residue** | **Ab Residue Number** | **Ab chain** |
| --- | --- | --- | --- | --- | --- |
| TRP | 353 | D | TYR | 27 | J |
| ASN | 354 | D | ASP | 28 | J |
| ARG | 355 | D | TYR | 27 | J |
| ARG | 355 | D | ASP | 28 | J |
| ARG | 355 | D | GLY | 29 | J |
| ARG | 355 | D | PHE | 100 | E |
| ARG | 355 | D | TYR | 100 | E |
| LYS | 356 | D | ASP | 28 | J |
| TYR | 396 | D | PHE | 100 | E |
| ASP | 428 | D | SER | 55 | E |
| ASP | 428 | D | ASN | 56 | E |
| ARG | 457 | D | PRO | 95 | J |
| LYS | 458 | D | ASP | 1 | J |
| LEU | 461 | D | TRP | 94 | J |
| LYS | 462 | D | TYR | 58 | E |
| LYS | 462 | D | LYS | 64 | E |
| LYS | 462 | D | TRP | 94 | J |
| PRO | 463 | D | TYR | 58 | E |
| PRO | 463 | D | TRP | 94 | J |
| PRO | 463 | D | TYR | 100 | E |
| PHE | 464 | D | TYR | 27 | J |
| PHE | 464 | D | TRP | 94 | J |
| PHE | 464 | D | TYR | 100 | E |
| PHE | 464 | D | PHE | 100 | E |
| GLU | 465 | D | TYR | 27 | J |
| GLU | 465 | D | THR | 92 | J |
| GLU | 465 | D | HIS | 93 | J |
| GLU | 465 | D | TRP | 94 | J |
| GLU | 465 | D | PRO | 95 | J |
| GLU | 465 | D | TYR | 100 | E |
| ARG | 466 | D | TYR | 27 | J |
| ARG | 466 | D | GLN | 27 | J |
| ARG | 466 | D | HIS | 93 | J |
| ASP | 467 | D | GLN | 27 | J |
| ASP | 467 | D | HIS | 93 | J |
| ILE | 468 | D | LEU | 27 | J |
| ILE | 468 | D | GLN | 27 | J |
| ILE | 468 | D | HIS | 93 | J |
| SER | 469 | D | SER | 26 | J |
| SER | 469 | D | GLN | 27 | J |
| THR | 470 | D | SER | 26 | J |
| THR | 470 | D | GLN | 27 | J |
| GLU | 471 | D | ASP | 1 | J |
| GLU | 471 | D | VAL | 3 | J |
| GLU | 471 | D | SER | 26 | J |
| GLU | 471 | D | GLN | 27 | J |
| SER | 514 | D | PHE | 100 | E |
| GLU | 516 | D | GLY | 98 | E |
| GLU | 516 | D | SER | 99 | E |
| GLU | 516 | D | PHE | 100 | E |
| LEU | 518 | D | LEU | 97 | E |
| LEU | 518 | D | GLY | 98 | E |
| LEU | 518 | D | SER | 99 | E |

**Table S6.** The list of the intermolecular contacts in the structure of the Class 5 WRAIR-2134 antibody complex with RBD (pdb id 8F2J). The interfacial contacts in the structure are defined by counting the number of interatomic contacts within a 5.5 Å distance threshold between atoms of the interacting proteins.

| **RBD Residue** | **RBD Residue Number** | **RBD chain** | **Ab Residue** | **Ab Residue Number** | **Ab chain** |
| --- | --- | --- | --- | --- | --- |
| ALA | 352 | C | TRP | 106 | A |
| ALA | 352 | C | ASP | 107 | A |
| TRP | 353 | C | TRP | 106 | A |
| TRP | 353 | C | ASP | 107 | A |
| TRP | 353 | C | TYR | 109 | A |
| ASN | 354 | C | TRP | 106 | A |
| ASN | 354 | C | ASP | 107 | A |
| ARG | 355 | C | ASP | 107 | A |
| ARG | 355 | C | ASP | 108 | A |
| ARG | 355 | C | TYR | 109 | A |
| ARG | 355 | C | TYR | 110 | A |
| LYS | 356 | C | ASP | 107 | A |
| LYS | 356 | C | ASP | 108 | A |
| ARG | 357 | C | TRP | 52 | A |
| ARG | 357 | C | ASP | 54 | A |
| ARG | 357 | C | SER | 56 | A |
| ARG | 357 | C | ASN | 57 | A |
| ARG | 357 | C | ASP | 108 | A |
| ASN | 394 | C | ASN | 57 | A |
| TYR | 396 | C | TRP | 52 | A |
| TYR | 396 | C | ASN | 57 | A |
| TYR | 396 | C | TYR | 59 | A |
| TYR | 396 | C | ASP | 108 | A |
| ASP | 427 | C | ASP | 37 | B |
| ASP | 428 | C | SER | 101 | B |
| ARG | 457 | C | TYR | 39 | B |
| ARG | 457 | C | GLN | 57 | B |
| ASN | 460 | C | TYR | 39 | B |
| LEU | 461 | C | TYR | 39 | B |
| LYS | 462 | C | LEU | 35 | B |
| LYS | 462 | C | GLY | 36 | B |
| LYS | 462 | C | ASP | 37 | B |
| LYS | 462 | C | LYS | 38 | B |
| LYS | 462 | C | TYR | 39 | B |
| LYS | 462 | C | ASP | 58 | B |
| LYS | 462 | C | ASN | 73 | B |
| LYS | 462 | C | TYR | 111 | A |
| PRO | 463 | C | ASP | 37 | B |
| PRO | 463 | C | SER | 100 | B |
| PRO | 463 | C | TYR | 111 | A |
| PHE | 464 | C | SER | 100 | B |
| PHE | 464 | C | TYR | 109 | A |
| PHE | 464 | C | TYR | 111 | A |
| GLU | 465 | C | TYR | 39 | B |
| GLU | 465 | C | GLN | 57 | B |
| GLU | 465 | C | TYR | 109 | A |
| GLU | 465 | C | TYR | 111 | A |
| GLU | 465 | C | TYR | 113 | A |
| ARG | 466 | C | TRP | 106 | A |
| ARG | 466 | C | ASP | 107 | A |
| ARG | 466 | C | TYR | 109 | A |
| ARG | 466 | C | TYR | 113 | A |
| ILE | 468 | C | TRP | 106 | A |
| GLU | 516 | C | ASN | 57 | A |
| GLU | 516 | C | TYR | 59 | A |
| GLU | 516 | C | THR | 102 | B |
| GLU | 516 | C | SER | 103 | B |
| LEU | 518 | C | LYS | 58 | A |
| LEU | 518 | C | TYR | 59 | A |
| LEU | 518 | C | THR | 102 | B |
| HIS | 519 | C | LYS | 65 | A |
| HIS | 519 | C | GLY | 66 | A |
